# Structure Activity Relationship of USP5 Allosteric Inhibitors

**DOI:** 10.1101/2021.05.17.444542

**Authors:** Mandeep K. Mann, Carlos A. Zepeda-Velázquez, Hector G. Alvarez, Aiping Dong, Taira Kiyota, Ahmed Aman, Cheryl H. Arrowsmith, Rima Al-Awar, Rachel J. Harding, Matthieu Schapira

## Abstract

USP5 is a deubiquitinase that has been implicated in a range of diseases, including cancer, but no USP5-targeting chemical probe has been reported to date. Here, we present the progression of a chemical series that occupies the C-terminal ubiquitin-binding site of a poorly characterized zinc-finger ubiquitin binding domain (ZnF-UBD) of USP5 and allosterically inhibits the catalytic activity of the enzyme. Systematic exploration of the structure-activity relationship, complemented with crystallographic characterization of the ZnF-UBD bound to multiple ligands, led to the identification of **64**, which binds to the USP5 ZnF-UBD with a K_D_ of 2.8 µM. **64** is selective over the structurally similar ZnF-UBD domain of HDAC6 and inhibits USP5 catalytic activity *in vitro* with an IC_50_ of 26 µM. This study provides a chemical and structural framework for the discovery of a chemical probe to delineate USP5 function in cells.

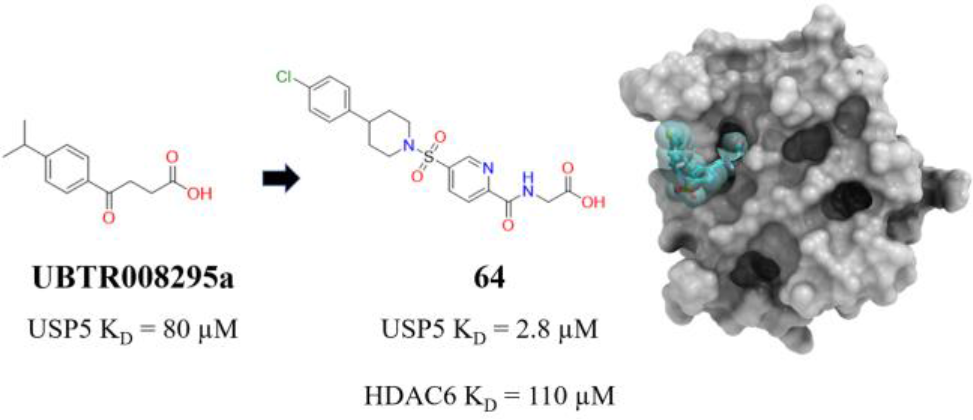
Table of Contents Graphic.

## INTRODUCTION

Ubiquitin specific proteases (USP) are the largest subfamily of deubiquitinating enzymes (DUB), consisting of more than 50 proteases with distinct roles in ubiquitin (Ub) biology. USP5 (also known as isopeptidase T, IsoT) cleaves unanchored ubiquitin chains, leading to the regeneration of monoUb^1, 2^ and removes Ub chains to stabilize post-translationally modified proteins in cells^3–12^. Deletion of UBP14, the gene encoding a yeast USP5 orthologue, leads to the accumulation of free polyUb chains and proteasome inhibition^13^. In addition to the proper functioning of the proteasome, USP5 has been shown to be important in neuropathic pain^4, 14–17^ and cancer^3, 7, 12, 18–24^. A genome-scale CRISPR-Cas9 knockout screen across 804 cells lines and 28 cancer types from the BROAD Institute reveals that, with USP36, USP5 is the most essential USP^25, 26^ (**Figure 1**). Interestingly, independent studies show that knocking-down USP5 is toxic in pancreatic cancer cell lines but has no effect in non-cancerous HEK293 cells^19^ and is well tolerated in mice^4^, suggesting that USP5 may be a valid therapeutic target in oncology.

**Figure 1.**
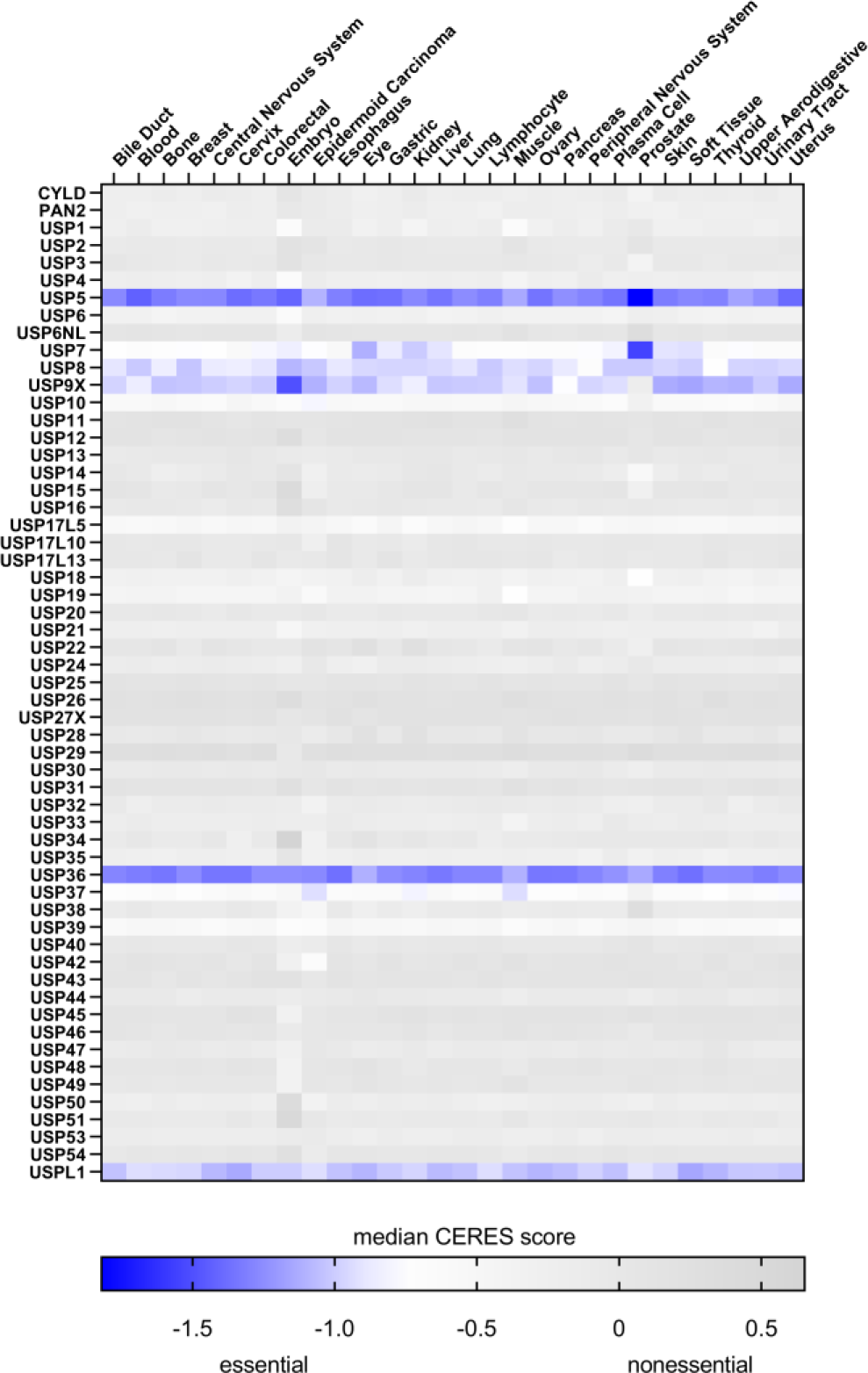
USP5 is essential in cancer. Heat map of the essentiality of USP proteins across 804 cancer cell lines and 28 cancer types was derived from Project Achilles data [CRISPR (Avana) Public 21Q1]^25, 26^. For each cancer type, the median CERES score across all associated cell lines was calculated. Genes are essential for cancer cell proliferation if the CERES score is lower than -1.

USP5 has multiple Ub binding modules: an N-terminal zinc finger ubiquitin-binding domain (ZnF-UBD) which recognizes the proximal free C-terminus of a broad subset of polyUb substrates^2, 27–29^, a USP domain that forms the active site, and two ubiquitin-binding associated domains (UBA-1, UBA-2)^30^ (**Figure 2a**). The mechanism of polyUb processing by USP5 is not understood for all polyUb chain variants; site-directed mutagenesis experiments show USP5 processes Met1 and Lys48-linked polyUb chains sequentially from the proximal end of Ub chains^2, 31^. The USP domain is highly conserved making it difficult to develop selective catalytic inhibitors; to date, out of 56 human USPs, selective inhibitors have been reported for USP1, USP7, USP9x, USP14 and USP30^32–38^. The ZnF-UBD domain is present in a limited number of proteins in the human genome: USP3, USP5, USP13, USP16, USP20, USP22, USP33, USP39, USP44, USP45, USP49, USP51, HDAC6, a histone deacetylase protein, and BRAP, a BRCA1-associated protein^39–41^. The role of the USP5 ZnF-UBD is not well understood. Hydrolysis of the artificial substrate, ubiquitin amidomethyl coumarin (Ub-AMC) by USP5 is activated when free monoUb occupies the ZnF-UBD, suggesting that this domain participates in the allosteric modulation of USP5^42^. However, in a separate study, deletion of the ZnF-UBD did not significantly affect hydrolysis of Ub-AMC^30^. It is therefore unclear whether targeting the ZnF-UBD is a valid strategy to antagonize the enzymatic activity of USP5.

**Figure 2.**
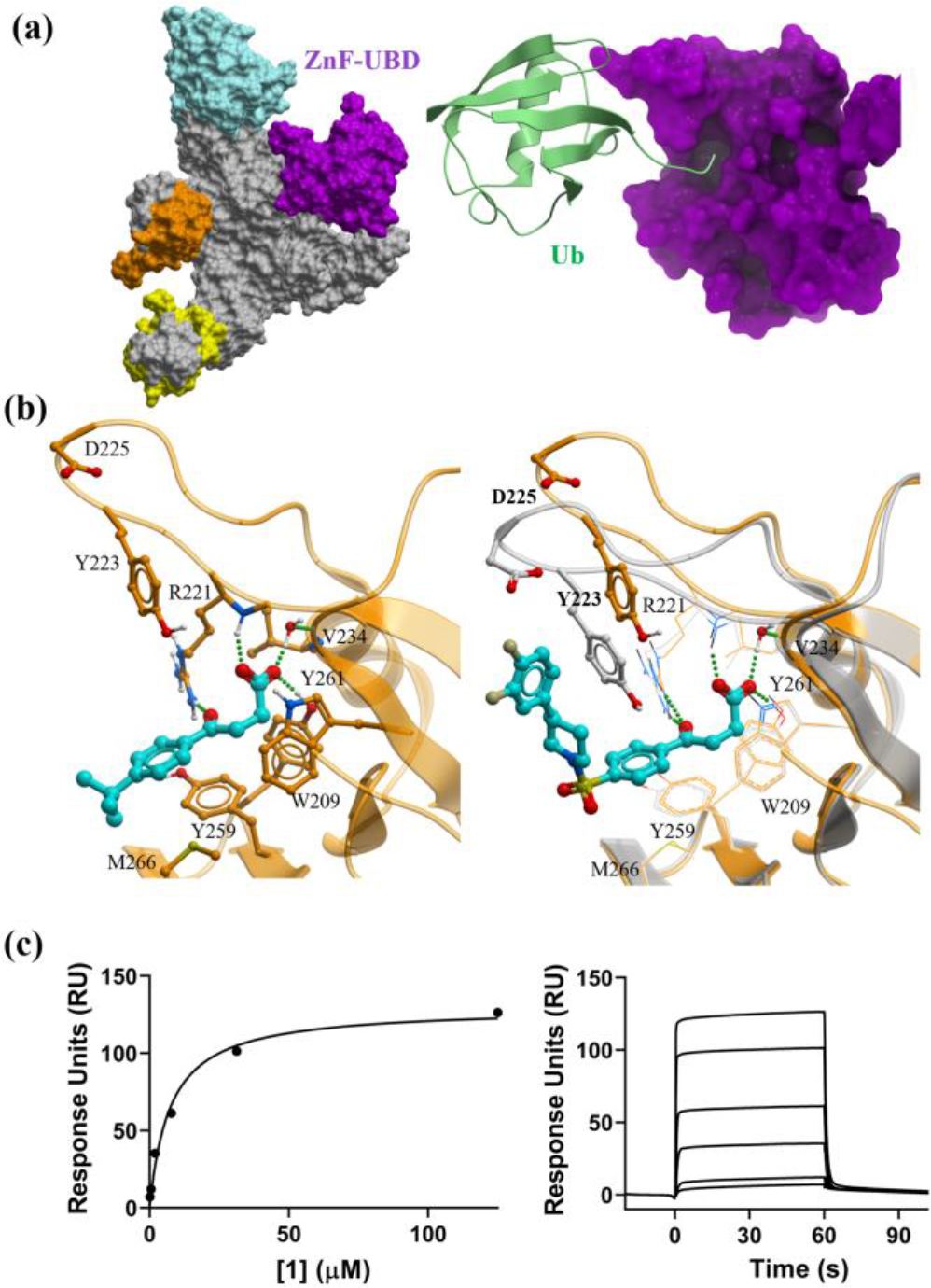
Structural characterization and binding affinity of 1: **(a)** (left) crystal structure of FL USP5 (grey) (PDB: 3IHP), with individual domains highlighted: cryptic N-terminal zinc-finger ubiquitin binding domain (nUBP) (cyan), ZnF-UBD (purple), UBA-1 (yellow), UBA-2 (orange). (right) Co-crystal structure of USP5 ZnF-UBD (purple) in complex with Ub (green) (PDB: 2G45). **(b)** (left) Co-crystal structure of USP5 ZnF-UBD (orange) in complex with a 4-phenyl-4-oxobutanoic acid fragment (PDB: 6DXH)^43^. (right) Co-crystal structure of USP5 ZnF-UBD chain A (grey) in complex with **1** (PDB: 7MS5) superimposed with the crystal structure of USP5 ZnF-UBD (PDB: 6DXH) (orange) highlighting the flexible loop residues Tyr223 and Asp225. The binding pocket residues and ligands are displayed as ball and sticks and the hydrogen bonds as green dashed lines. **(c)** Representative SPR binding curve and sensorgram of USP5 ZnF-UBD and **1** (K_D_= 12 ± 3 µM).

We previously reported fragment-like molecules that bind to the USP5 ZnF-UBD with high micromolar affinity^43^. Here, we describe the progression of one of these compounds into a low micromolar USP5 inhibitor. We show that our ligand can displace Ub from the ZnF-UBD in the context of full-length (FL) USP5, inhibits the catalytic function of USP5 with a Lys48-linked di-ubiquitin (Ub2K48) substrate *in vitro* and is selective over HDAC6 ZnF-UBD. The accompanying crystal structures and structure activity relationship (SAR) provide a robust framework for the development of a chemical probe to interrogate the function of USP5 in cells and *in vivo*.

## RESULTS & DISCUSSION

### Hit Expansion

We focused on exploring the SAR around **UBTR008295a**, a 4-phenyl-4-oxobutanoic acid USP5 ZnF-UBD ligand previously reported (K_D_= 80 µM)^43^. A substructure search of the chemical scaffold from the Enamine REAL database resulted in 285 analogues, which were docked to the USP5 ZnF-UBD crystal structure (PDB: 6DXH)^43^ using Glide^44^ (Schrodinger, NY), with three hydrogen bond constraints at the side chain NH of Arg221, the NH backbone of Arg221 and the side chain OH of Tyr261. Ten compounds were ordered and tested using a fluorescence competition assay, in which displacement of a N-terminal fluorescently labeled Ub substrate from the ZnF-UBD was measured. Nine compounds displaced Ub from the ZnF-UBD at micromolar concentration and were tested using an orthogonal surface plasmon resonance (SPR) assay (**Table S2**). Compound **1,** with an extended sulfonamide group, improved potency 7-fold against the USP5 ZnF-UBD (K_D_=12 µM) compared to **UBTR008295a** (K_D_=80 µM)^43^ (**Table 1**).

**Table 1.**
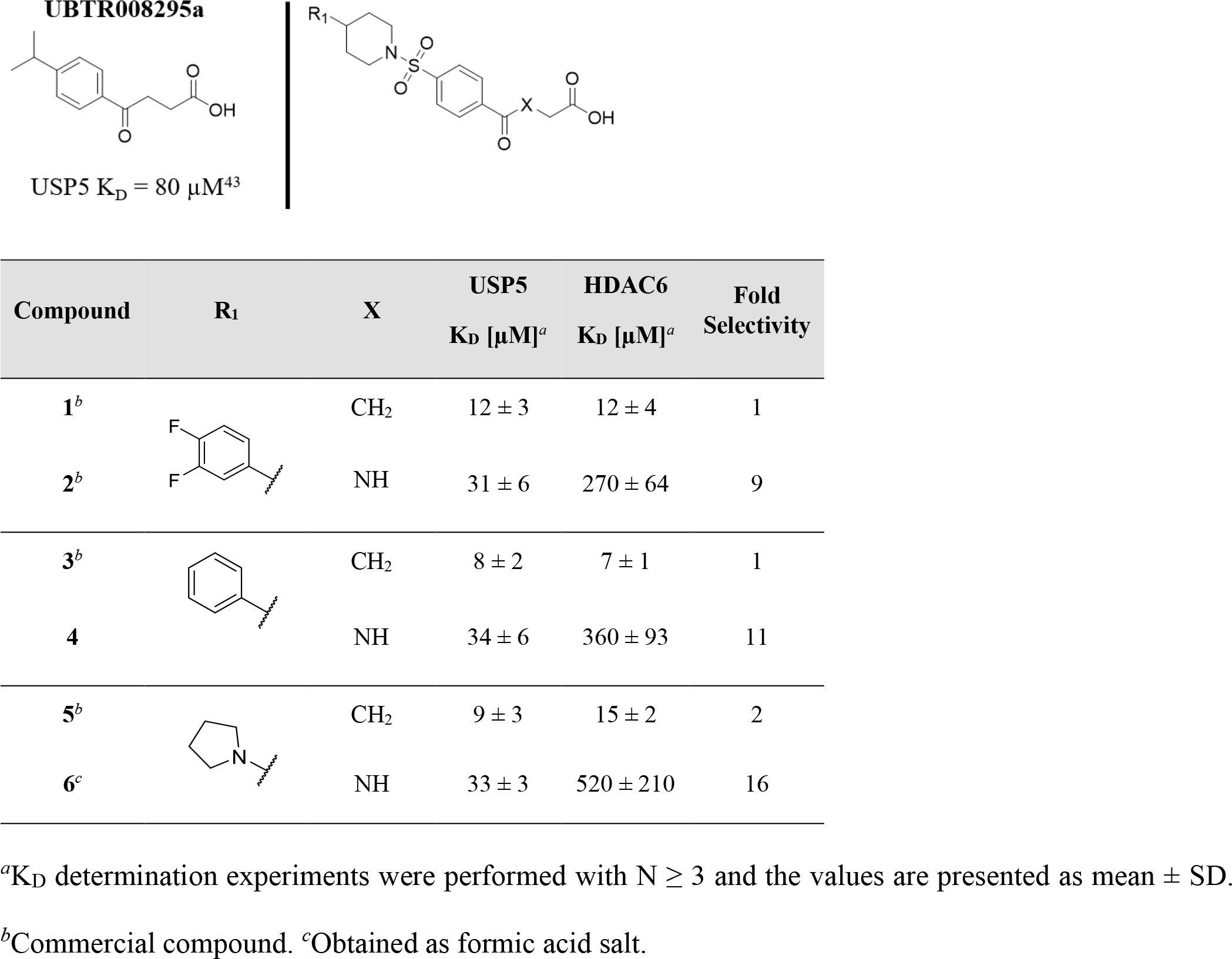
Potency and selectivity of 1-6

| Compound | R<sub>1</sub> | X | USP5<br>$K_D$ [$\mu\text{M}$]<sup>a</sup> | HDAC6<br>$K_D$ [$\mu\text{M}$]<sup>a</sup> | Fold<br>Selectivity |
| --- | --- | --- | --- | --- | --- |
| **1<sup>b</sup>** |  | CH<sub>2</sub> | 12 ± 3 | 12 ± 4 | 1 |
| **2<sup>b</sup>** |  | NH | 31 ± 6 | 270 ± 64 | 9 |
| **3<sup>b</sup>** |  | CH<sub>2</sub> | 8 ± 2 | 7 ± 1 | 1 |
| **4** |  | NH | 34 ± 6 | 360 ± 93 | 11 |
| **5<sup>b</sup>** |  | CH<sub>2</sub> | 9 ± 3 | 15 ± 2 | 2 |
| **6<sup>c</sup>** |  | NH | 33 ± 3 | 520 ± 210 | 16 |

<sup>a</sup> $K_D$ determination experiments were performed with $N \geq 3$ and the values are presented as mean ± SD.
<sup>b</sup>Commercial compound. <sup>c</sup>Obtained as formic acid salt.

**Table 2.** Potency and selectivity of 9-27

| Compound | R<sub>1</sub> | USP5<br>K<sub>D</sub><br>[μM]<sup>a</sup> | HDAC6<br>K<sub>D</sub><br>[μM]<sup>a</sup> | Fold<br>Selectivity | Compound | R<sub>1</sub> | USP5<br>K<sub>D</sub><br>[μM]<sup>a</sup> | HDAC6<br>K<sub>D</sub><br>[μM]<sup>a</sup> | Fold<br>Selectivity |
| --- | --- | --- | --- | --- | --- | --- | --- | --- | --- |
| **4** |  | 34 ± 6 | 360 ± 93 | 11 | **18<sup>d</sup>** |  | 87 ± 23 | >530 | >6 |
| **9** |  | 170 ± 31 | >480 | >3 | **19** |  | 61 ± 26 | >540 | >9 |
| **10** |  | 110 ± 7 | 290 ± 63 | 3 | **20** |  | 130 ± 7 | >480 | >4 |
| **11** |  | 270 ± 27 | NB<sup>b</sup> | - | **21** |  | 220 ± 62 | 440 ± 67 | 2 |
| **12<sup>c</sup>** |  | 130 ± 40 | NB<sup>b</sup> | - | **22<sup>e</sup>** |  | 80 ± 21 | >760 | >10 |
| **13<sup>c</sup>** |  | 94 ± 27 | >640 | >7 | **23** |  | 38 ± 9 | 63 ± 27 | 2 |
| **14** |  | 150 ± 19 | >480 | >3 | **24<sup>e</sup>** |  | 45 ± 8 | NB<sup>b</sup> | - |
| **15** |  | 54 ± 5 | 340 ± 63 | 6 | **25<sup>e</sup>** |  | 34 ± 6 | 370 ± 57 | 11 |
| **16<sup>c</sup>** |  | 90 ± 21 | >570 | >6 | **26<sup>e</sup>** |  | 150 ± 42 | NB<sup>b</sup> | - |
| **17<sup>c</sup>** |  | 92 ± 25 | >570 | >6 | **27<sup>e</sup>** |  | 25 ± 2 | 410 ± 76 | 16 |

<sup>a</sup>K<sub>D</sub> determination experiments were performed with N ≥ 3 and the values are presented as mean ± SD.
<sup>b</sup>K<sub>D</sub> determination experiments were performed with N = 2. <sup>c</sup>Mixture of diastereomers/enantiomers.
<sup>d</sup>Obtained as formic acid salt. NB: no binding (K<sub>D</sub> > 1 mM).

ZnF-UBDs share similar sequence and structural folds and the ZnF-UBD of USP3, USP5, USP16 and HDAC6 bind Ub^31, 40, 45–47^. We have previously shown that the ZnF-UBD of HDAC6 can be targeted by small molecules inhibitors that, as compound **1**, include a carboxylic tail mimicking the C-terminal di-glycine motif of Ub^48, 49^. We therefore decided to use HDAC6 as an anti-target to monitor the selectivity of our compounds. Ultimately, an extended selectively panel should be applied to characterize the specificity of more advanced compounds. We found that **1** bound with equipotency to ZnF-UBDs of USP5 and HDAC6 (USP5 & HDAC6 K_D_=12 µM).

To confirm the binding pose of **1** to USP5 ZnF-UBD, the crystal structure of USP5 ZnF-UBD in complex with **1** was solved to 1.98 Å resolution (PDB: 7MS5). Compound **1** was well resolved in the electron density and modeled for two of the USP5 ZnF-UBD molecules in the crystallographic asymmetric unit.

The electron density of **1** was unambiguous in chain A and was reasonably modeled in chain B. The ligand pose in both molecules differs, with a RMSD of 3.11, due to, the fluorinated ring of **1** stacking on a Phe224 residue from an adjacent molecule in the lattice of the asymmetric unit in chain B (**Figure S1**). As expected, the carboxylate group of **1** recapitulates hydrogen bond interactions observed with the Ub C-terminal glycine carboxylic acid (**Figure 2b**). The backbone amide of Arg221 and side chain OH of Tyr261 are engaged in direct hydrogen bonds with the buried carboxylic acid of the ligand which also makes water-mediated hydrogen bonds with surrounding residues (Arg221, Val234). Additionally, the central aromatic ring is engaged in π-stacking interactions with Tyr259. The angle afforded by the sulfonyl favorably positions the 4-phenylpiperidine group for additional interactions with binding pocket residues. A shift in backbone conformation of a loop that is forming part of the binding pocket (residues 221-230) leads to repositioning of Try223 to optimally accommodate the 4-phenylpiperidine appendage of **1** (**Figure 2b, Figure S2**). In particular, Tyr223 engages in an edge-to-face π-stacking interaction with the terminal phenyl group of **1**, which may account for the improved potency of the ligand.

To identify molecules structurally related to **1**, a similarity search was performed against the Enamine REAL database and SciFinder. 26 compounds were docked to USP5 ZnF-UBD (chain A) in complex with **1** (PDB: 7MS5) with Glide^44^ with the three hydrogen bond constraints described previously. Sixteen compounds were ordered and tested by SPR (**Table S3**). Compound **3**, which lacks fluorine substitutions at positions C3 and C4 of the terminal phenyl, and **5** which has a terminal pyrrolidine had marginally improved potency compared with **1** but were not selective (USP5 K_D_= 8 and 9 µM respectively; HDAC6 K_D_= 7 and 15 µM respectively). Interestingly, the introduction of an amide to the carboxylate chain in compound **2** resulted in a 9-fold gain in selectivity versus HDAC6 (USP5 K_D_=31 µM; HDAC6 K_D_=270 µM). The amide likely stabilizes the co-planarity of the carboxylate tail with the aromatic ring, which may be unfavorable for HDAC6 binding. We synthesized **4** and **6**, the amide analogs of **3** and **5** respectively, to confirm that the amide confers selectivity for USP5 over HDAC6 (**Scheme 1**). Both compounds exhibited comparable affinity for USP5 (K_D_= 34, 33 µM respectively) and selectivity over HDAC6 (11 and 16-fold respectively). Compound **4** was identified as a more favorable scaffold for developing potent and selective ligands and we next focused on further exploring the SAR around **4**.

**Table 3.** Potency and selectivity of 28-38

| Compound | R<sub>1</sub> | USP5<br>K<sub>D</sub><br>[μM]<sup>a</sup> | HDAC6<br>K<sub>D</sub><br>[μM]<sup>a</sup> | Fold<br>Selectivity | Compound | R<sub>1</sub> | USP5<br>K<sub>D</sub><br>[μM]<sup>a</sup> | HDAC6<br>K<sub>D</sub><br>[μM]<sup>a</sup> | Fold<br>Selectivity |
| --- | --- | --- | --- | --- | --- | --- | --- | --- | --- |
| 4 |  | 34 ± 6 | 360 ± 93 | 11 | 33 |  | 79 ± 31 | NB<sup>b</sup> | - |
| 28 |  | 19 ± 5 | 150 ± 34 | 8 | 34 |  | 18 ± 4 | NC | - |
| 29 |  | 40 ± 5 | 130 ± 16 | 3 | 35 |  | 39 ± 6 | 150 ± 9 | 4 |
| 30 |  | 23 ± 4 | 400 ± 48 | 17 | 36 |  | 31 ± 3 | 200 ± 25 | 6 |
| 31 |  | 21 ± 8 | 150 ± 43 | 7 | 37<sup>c</sup> |  | NC | NC | - |
| 32 |  | 22 ± 6 | >160 | >7 | 38<sup>c</sup> |  | NC | NC | - |

<sup>a</sup>K<sub>D</sub> determination experiments were performed with N ≥ 3 and the values are presented as mean ± SD.
<sup>b</sup>K<sub>D</sub> determination experiments were performed with N = 2. <sup>c</sup>Compound with poor solubility in assay conditions. NB: no binding (K<sub>D</sub> > 1 mM). NC: not calculated.

### Structure Activity Relationship

Our initial SAR studies focused on varying substitutions to the carboxylic acid to determine if less acidic moieties could interact with Arg221 and Tyr261 (**Table S4**). Substituting the carboxylic acid to an ester (**S25**) or tetrazole (**S26**, **S27**) resulted in a complete loss of binding against USP5 indicating that the conserved network of direct and water-mediated interactions between the carboxylate and Arg221, Tyr261 and Val234 is essential for binding, as was previously observed with the corresponding binding pocket of HDAC6^49^. The addition of a (S)-methyl or dimethyl at the C2 position of the carboxylic chain and mono-N-methylation of the secondary amide were not tolerated (**S28**, **S30**, **S31**). A (R)-methyl at the C2 position of the carboxylic chain (**S29**) had weaker affinity for USP5 and improved affinity for HDAC6. Lengthening and shortening of the carboxylic chain was also not tolerated (**S32**, **S24**).

**Table 4.** Potency and selectivity of 44-57

| Compound | Core | USP5<br>$K_D$<br>[$\mu M$]<sup>a</sup> | HDAC6<br>$K_D$<br>[$\mu M$]<sup>a</sup> | Fold<br>Selectivity | Compound | Core | USP5<br>$K_D$<br>[$\mu M$]<sup>a</sup> | HDAC6<br>$K_D$<br>[$\mu M$]<sup>a</sup> | Fold<br>Selectivity |
| --- | --- | --- | --- | --- | --- | --- | --- | --- | --- |
| **4** | | $34 \pm 6$ | $360 \pm 93$ | 11 | **51** | | $7 \pm 2$ | $200 \pm 57$ | 29 |
| **44** | | $82 \pm 15$ | $>440$ | $>5$ | **52** | | $22 \pm 1$ | $160 \pm 16$ | 7 |
| **45** | | $42 \pm 5$ | $210 \pm 5$ | 5 | **53** | | $21 \pm 2$ | $70 \pm 23$ | 3 |
| **46** | | $47 \pm 9$ | $200 \pm 21$ | 4 | **54** | | $61 \pm 7$ | $390 \pm 36$ | 6 |
| **47** | | $59 \pm 7$ | $130 \pm 2$ | 2 | **55** | | $67 \pm 14$ | $310 \pm 98$ | 5 |
| **48** | | $9 \pm 2$ | $30 \pm 10$ | 3 | **56** | | $71 \pm 12$ | $>500$ | $>7$ |
| **49** | | $210 \pm 79$ | $150 \pm 65$ | 1 | **57** | | $150 \pm 20$ | NB<sup>b</sup> | - |
| **50** | | NB | $160 \pm 16$ | - | | | | | |

<sup>a</sup> $K_D$ determination experiments were performed with $N \geq 3$ and the values are presented as mean ± SD.
<sup>b</sup> $K_D$ determination experiments were performed with $N = 2$ . NB: no binding ( $K_D > 1$ mM).

Molecules **9-38** were synthesized following a well-stablished synthetic procedure (**Scheme 1**). (4-(chlorosulfonyl)benzoyl)glycine was treated with various commercially available or synthesized 4-substituted-piperidylamines or related analogs under basic conditions to obtain the N-substituted-sulfonamides. Investigation of the SAR around the piperidine (**Table 2, 9-27**), revealed the 6-membered piperidine ring of **4** was preferred over 4, 5, and 7-membered rings (**11-13)**. The phenylpiperidine of compound **4** had at least 3-fold improved affinity for USP5 ZnF-UBD over the bicyclic ring systems of compounds **9** and **10** which had isoindoline and dihydroisoquinoline groups, respectively. Finally, a piperidine was preferred over a piperazine (**14**), 1, 2, 3, 6-tetrahydropyridine (**15**), 2- and 3-methylpiperadine (**16**, **17**), or a bridged-bicyclic ring (**18)**. Substitutions of the piperidine ring (compound **19-25**) showed that a phenyl ring (**4**) or a N-methyl-piperazine (**25**) were preferred at this position. The piperidine substituted with a phenyl (**4**) is preferred over other groups for efficient positioning of the terminal phenyl for π-stacking interactions with Tyr223, as seen with the complex structure of **1** (PDB: 7MS5). A spiro group (**27**) was found to be slightly more potent and selective than **4**, but chemical tractability considerations led us to further explore substitutions on the phenyl ring of **4**.

Substitutions at the terminal phenyl ring of **4** with methyl or halogen groups in ortho improved affinity by about 2-fold (**28, 30-32**), while a methoxy group decreased potency and selectivity (**29**) (**Table 3**). Meta-substitution with a methyl (**34**) markedly improved affinity (K_D_= 18 µM) while other groups at the same position were not favorable (**33, 35-38**).

To examine the role of the central aromatic core we synthesized and surveyed different substituted aromatic and heteroaromatic scaffolds (**Table 4**) following a previously described methodology (**Scheme 2**), with slight modifications, as shown in **Scheme 3**. For compounds, **44-57**, 4-phenylpiperidine was reacted with the sulfonyl chloride of the core, to form the corresponding N-substituted sulfonamide. The condensation reaction between the carboxylic acid and *tert*-butyl glycinate was achieved through application of a coupling agent, HATU. The final step involved an ester deprotection under acid conditions to obtain the desired molecule. Compound **51** required a saponification step before the carboxamide formation.

Comparing **4** vs. **44** indicated that transversal branching afforded by -1,4 substitution was favored over the -1,3 trajectory. Addition of a methyl or halogen in the C3 of the phenyl ring (**45-47**) improved neither potency nor selectivity; however, a fluorine in C2 resulted in a four-fold increase in potency **48**: K_D_= 9 ± 2 µM, while bulkier groups such as chlorine or methyl (**49**, **50**) were not tolerated. We solved the crystal structure of **48** (PDB: 7MS6) in complex with USP5 ZnF-UBD to a resolution of 1.55 Å with unambiguous electron density for the bound ligand (**Figure S3**). The fluorine group of **48** makes hydrophobic interactions with Trp209, which may account for the 4-fold increase in potency for USP5 ZnF-UBD in comparison to **4** (**Figure 3**). However, while the added fluorine increases affinity for USP5, it also results in a four-fold decrease in selectivity against HDAC6.

**Figure 3.**
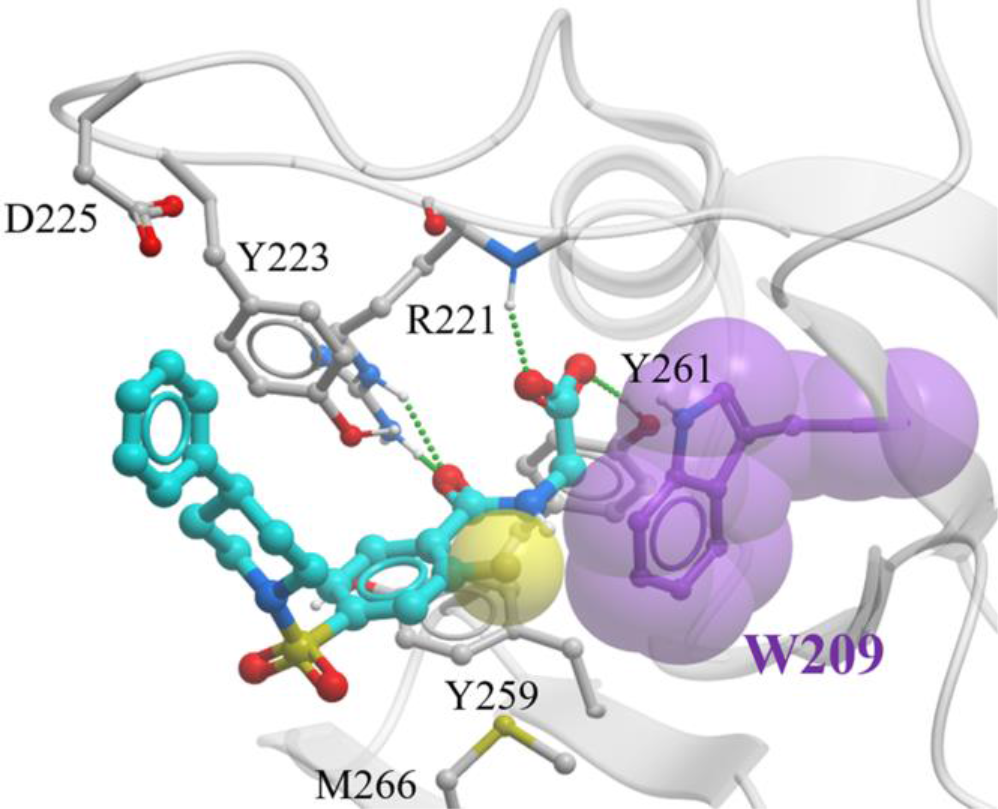
Co-crystal structure of USP5 ZnF-UBD in complex with 48 (PDB: 7MS6) The Trp209 residue (purple) in the binding pocket of USP5 and the fluorine (yellow) of **48** are represented as space-filling models. The binding pocket residues and ligands are displayed as ball and sticks and the hydrogen bonds as green dashed lines.

The effect of replacing the core phenyl ring with other ring systems was also explored (**Table 4**, compounds **51-57**). Importantly, introducing a pyridine ring (compound **51**) improved potency for USP5 five-fold (K_D_= 7 ± 2 µM) and selectivity over HDAC6 (K_D_= 200 ± 57 µM). Core hopping with 5-membered heterocycles led to no significant improvement in potency or selectivity. (**52-57**).

To build upon the improvement afforded by the fluoro-phenyl of **48** and the pyridine of **51**, four derivatives of these compounds were synthesized and tested (Compounds **61-64**; **Scheme 3**; **Table 5**). While no apparent benefit was observed in replacing the terminal phenyl ring of **48**, we were pleased to observe further improvement in potency and selectivity when chlorinating this ring in **51** (compound **64**: USP5 K_D_= 2.8 µM: HDAC6 K_D_=110 µM). We solved the crystal structure of **64** in complex with USP5 ZnF-UBD to 1.45 Å resolution (PDB: 7MS7). The USP5-**64** complex is modeled with high confidence in the context of chain A; however, the electron density of **64** was not fully resolved in chain B and has ambiguous density for ligand positioning due to the adjacent molecule in the lattice partially blocking the binding site for the crystallographic asymmetric unit which is likely reducing occupancy of this ligand pocket (**Figure S4**). **64** makes similar interactions with binding pockets residues as **1** (PDB: 7MS5) and **48** (PDB: 7MS6), including hydrogen bond interactions with the backbone and side chains of Arg221, side chain of Tyr261, and π-stacking interaction with Tyr259 and Tyr223 (**Figure 4**). The pyridine nitrogen of **64** can engage in an intramolecular hydrogen bond with the amide nitrogen of the aliphatic tail, which reinforces π-stacking interactions with Tyr259, Tyr223 and Trp209 through electronic effects and contributes to the gain in potency. We observe significant variation in the backbone and side-chain conformation of residues 223-225, at the extremity of the loop lining the ligand binding pocket, confirming our previously reported observation that this loop can rearrange structurally to accommodate different ligands^43^ (**Figure S5**).

**Figure 4.**
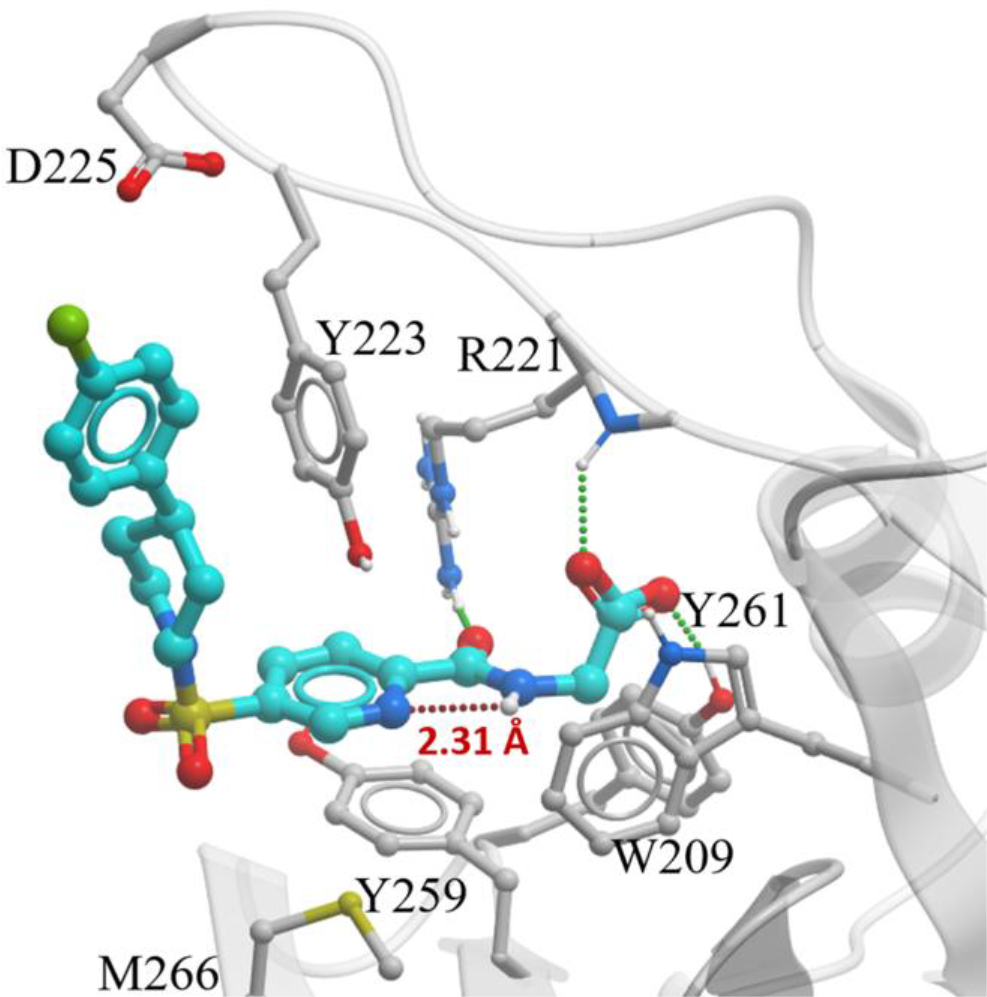
Co-crystal structure of USP5 ZnF-UBD in complex with 64 (PDB: 7MS7). Binding pocket residues and ligands are displayed as ball and sticks. Intermolecular and intramolecular hydrogen bonds are shown as green and red dashed lines respectively.

**Table 5.**
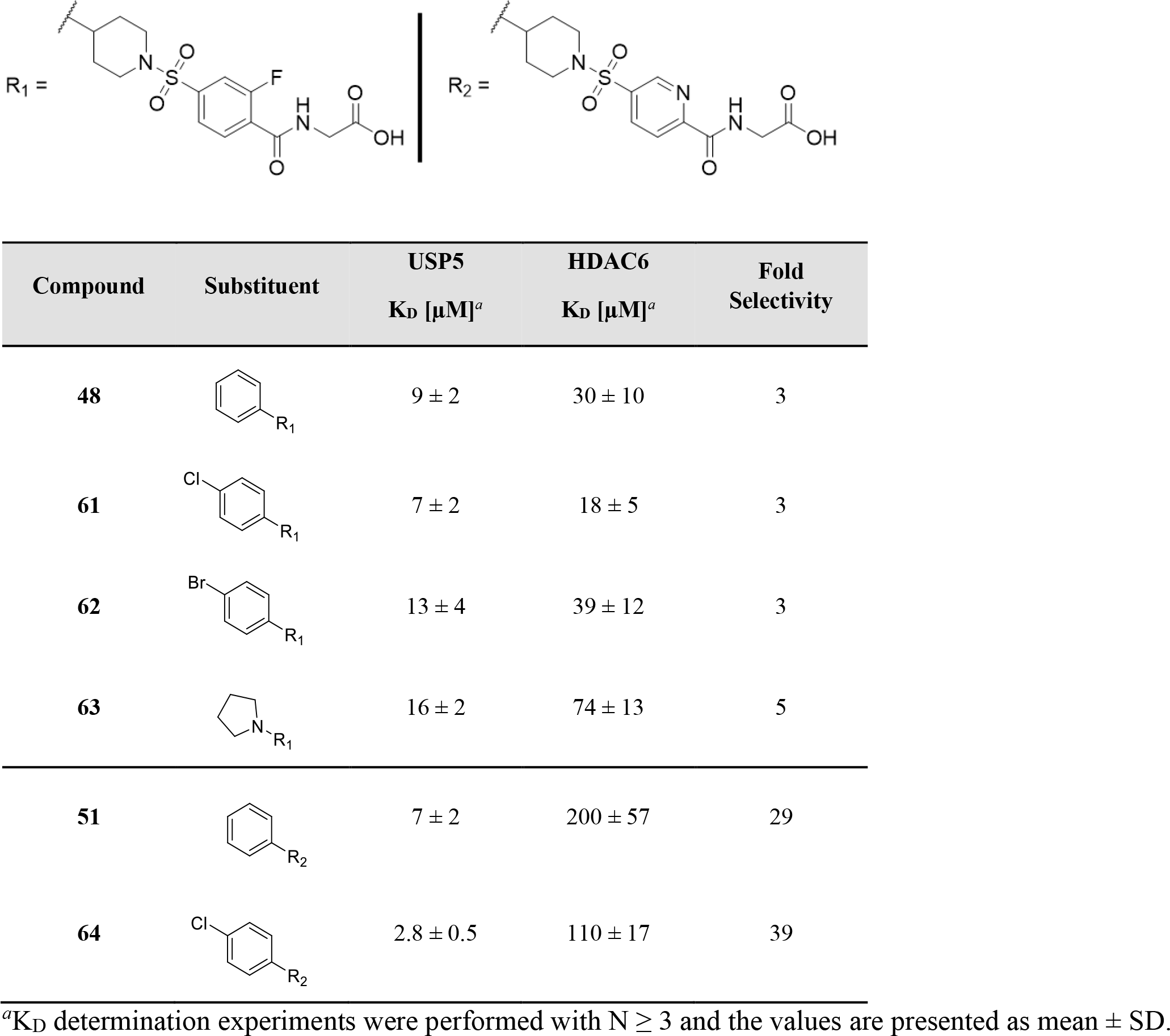
Potency and selectivity of 48, 51 and 61-64

| Compound | Substituent | USP5<br>K<sub>D</sub> [μM]<sup>a</sup> | HDAC6<br>K<sub>D</sub> [μM]<sup>a</sup> | Fold<br>Selectivity |
| --- | --- | --- | --- | --- |
| **48** |  | 9 ± 2 | 30 ± 10 | 3 |
| **61** |  | 7 ± 2 | 18 ± 5 | 3 |
| **62** |  | 13 ± 4 | 39 ± 12 | 3 |
| **63** |  | 16 ± 2 | 74 ± 13 | 5 |
| **51** |  | 7 ± 2 | 200 ± 57 | 29 |
| **64** |  | 2.8 ± 0.5 | 110 ± 17 | 39 |

<sup>a</sup>K<sub>D</sub> determination experiments were performed with N ≥ 3 and the values are presented as mean ± SD.

### Characterization of Compound 64

We used SPR to show that lead compound **64** binds to the full-length (FL) USP5 with a K_D_ of 8 ± 2 µM, 2.5-fold weaker in binding affinity compared to the isolated ZnF-UBD (K_D_ = 2.8 ± 0.5 µM) (**Figure 5ab**). **64** shows no detectable binding to a FL R221A USP5 mutant (**Figure S6**), indicating that **64** is binding only at the ZnF-UBD in the context of the FL enzyme. *In vitro* displacement of Ub was tested using a fluorescence competition assay measuring the displacement of a N-terminal FITC-tagged Ub from both the ZnF-UBD and the FL USP5 (**Figure 5c**). We found that compound **64** decreased the interaction of Ub from the isolated ZnF-UBD and the FL USP5 in a dose-dependent manner with an IC_50_ of 7 ± 2 µM and 46 ± 15 µM, respectively. **64** displaces Ub approximately seven-fold less efficiently from FL USP5 than from the isolated ZnF-UBD; suggesting that in the context of the FL protein, the Ub molecule occupying the ZnF-UBD is also making stabilizing interactions with other domains of USP5; this is in agreement with the five-fold difference in binding affinity of Ub to the ZnF-UBD (K_D_= 1.4 µM) versus FL USP5 (K_D_=0.3 µM)^43^ (**Figure S7**). Importantly, we observed dose-dependent inhibition of USP5 catalytic activity by **64** in an internally quenched fluorophore (IQF) assay with a native K48-linked di-ubiquitin (Ub2K48) substrate, with an IC_50_ of 26 ± 9 µM *in vitro* (**Figure 5d**). Taken together, these data suggest that targeting the ZnF-UBD is a valid strategy for the selective inhibition of USP5, a putative cancer target. The potency of the compounds presented in this work needs to be further improved before inhibitors can be tested in cellular systems or animal models. To evaluate the development potential of this chemical series and support further optimization efforts, we verified that our most advanced compound was stable in mouse and human liver microsome stability assays (>90% compound remaining after 60 min, **Table S5**).

**Figure 5.**
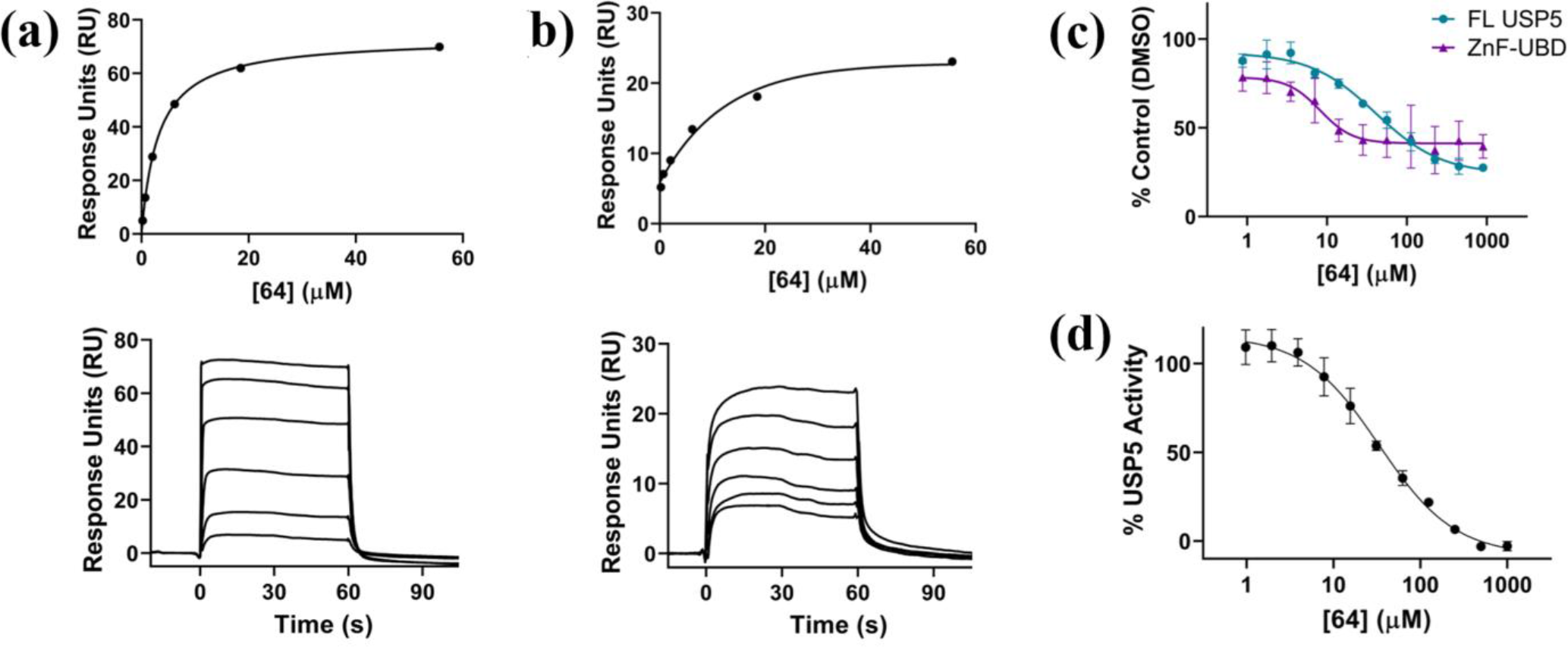
Characterization of 64: (a) Representative SPR binding curve and sensorgram for USP5 ZnF-UBD. A K_D_ of 2.8 ± 0.5 µM was obtained from the average of seven independent measurements (b) SPR binding curve and sensorgram for FL USP5. A K_D_ of 8 ± 2 µM was obtained from the average of three independent measurements (c) Fluorescence competition assay with a N-terminal FITC-labeled Ub and USP5 ZnF-UBD and FL USP5. An IC50 of 7 ± 2 µM and 46 ± 15 µM was obtained from the average of three independent measurements for the ZnF-UBD and FL USP5, respectively. Fluorescence signal is normalized to control (no compound, DMSO only) (d) Inhibition of USP5 catalytic activity on a Ub2K48 substrate. An IC50 of 26 ± 9 µM was obtained from the average of three independent measurements. Fluorescence signal is normalized to control (no compound, DMSO only).

## CONCLUSION

In summary, we optimized a preliminary chemical scaffold through structural analysis, docking and SAR studies to identify a small molecule antagonist targeting the USP5 ZnF-UBD with low micromolar affinity that is selective over a structurally similar ZnF-UBD in HDAC6. We show that compound **64** occupies the USP5 ZnF-UBD, can displace Ub in the context of FL USP5 and inhibits USP5 deubiquitinase activity.

The ZnF-UBD has been hypothesized to recognize and position substrates or allosterically modulate USP5 catalytic activity^30, 31^. Our antagonists inhibit USP5-mediated cleavage of a Ub2K48 substrate showing that when the ZnF-UBD is blocked, Ub2K48 cannot be properly positioned for catalytic cleavage. USP5 was reported to cleave a diverse array of differently branched polyubiquitin substrates, and it remains to be seen whether the inhibitory activity of ZnF-UBD ligands is substrate-specific. The chemical inhibitors and accompanying crystal structures reported here will enable further optimization into a potent and selective chemical probe to investigate the cellular function of USP5.

## EXPERIMENTAL SECTION

### Hit Expansion & Docking

Substructure search was run against ∼1.2 billion compounds from the Enamine REAL database (Enamine Ltd., 2019) and SciFinder. Ligprep (Schrodinger, New York) was used to prepare the ligands using default settings. The X-ray structures of USP5 ZnF-UBD (PDB ID: 6DXH^43^, 7MS5) were prepared with PrepWizard^50^ (Schrodinger, New York) for the assignment of bond order, assessment of correct protonation states and optimization of hydrogen-bond assignment at pH 7.3, and restrained minimization using the OPLS3e force field. Receptor grids were calculated at the centroid of the co-crystallized ligand for each respective complex. The library was docked to the USP5 ZnF-UBD structure for each respective complex using Glide SP^44^ (Schrodinger, New York) with hydrogen-bond constraints with the NH of Arg221 backbone, NH of Arg221 side chain and OH of Tyr261. Docked compounds were clustered in ICM^51^ and compounds were selected to be purchased based on docking pose, score, visual inspection, and cost.

### Cloning, Protein Expression and Purification

cDNAs encoding Ub^1–76^, USP5^171–290^, USP5^1–835^, USP5^1–835 R221A^, HDAC6^1109–1215^ were cloned as previously described^43, 48, 49^. Proteins were purified as before^43, 48, 49^; briefly, all proteins were expressed in *Escherichia coli* and purified by metal affinity chromatography, tag cleavage, gel filtration and ion exchange chromatography. The final concentration of purified proteins was 2-45 mg/mL by UV absorbance at 280 nm. Protein identity was confirmed by mass spectrometry and purity was assessed by SDS-PAGE, showing all samples to be >90% pure.

### Fluorescence Competition Assay

All experiments were performed in 384-well black flat-bottomed streptavidin plates (Greiner). 20 µL of 1 µM biotinylated-USP5^171–290^, USP5^1–835^ prepared with buffer (20 mM HEPES pH 7.4, 150 mM NaCl, 1 mM TCEP, 0.005% Tween-20 (v/v), 1% DMSO (v/v)) was incubated in wells for 1 hour at 4⁰C. Wells were washed with 3x50 µL buffer to remove excess unbound protein. 20 µL of two-fold dilution series of ligand and 0.2 µM N-terminally FITC-labeled Ub (Boston Biochem) prepared in buffer were incubated in wells for 1 hour at 4⁰C. Wells were washed with 7x50 µL buffer to remove excess unbound FITC-UBQ. Fluorescence was measured using a Biotek Synergy H1 microplate reader (Biotek) at emission and excitation of 528 nm and 485 nm, respectively. The data were analyzed with GraphPad Prism 8.4.2.

### Surface Plasmon Resonance Assay

Studies were performed with a Biacore T200 (GE Health Sciences) at 20⁰C. Approximately 3000-6000 response units (RU) of biotinylated USP5^171-290^, HDAC6^1109–1215^, USP5^1–835^, USP5^1–835 R221A^ were captured to flow cells of a streptavidin-conjugated SA chip per manufacturer’s protocol, and an empty flow cell used for reference subtraction. Serial dilutions were prepared in 20 mM HEPES pH 7.4, 150 mM NaCl, 1 mM TCEP, 0.005% Tween-20 (v/v), 1% DMSO (v/v). K_D_ determination experiments were performed using multi-cycle kinetics with 60-120 s contact time, and 60-120 s dissociation time at a flow rate of 30 µL/min. K_D_ values were calculated using steady state affinity fitting with the Biacore T200 evaluation software (GE Health Sciences).

### Crystallization

12 mg/mL of tag-free USP5^171–29^ and 1.1% (v/v) of 200 mM DMSO-solubilized stock of compounds were prepared in 50 mM Tris pH 8, 150 mM NaCl, 1 mM TCEP. Sparse-matrix crystallization experiments yielded crystals in different conditions for each USP5 ZnF-UBD-compound complex. **1** was co-crystallized with 1:1 protein/compound: mother liquor (1 M sodium/potassium phosphate pH 6.9). **48** was co-crystallized with 1:1 protein/compound: mother liquor (2 M ammonium sulfate, 0.2 M sodium acetate, 0.1 M HEPES pH 7.5, 5% MPD). **64** was co-crystallized with 1:1 protein/compound: mother liquor (20% PEG3350, 0.2 M magnesium acetate). Co-crystals were soaked in a cryoprotectant consisting of mother liquor supplemented with 25% ethylene glycol (v/v) for **1** and **48** and 20% glycerol (v/v) for **64** prior to mounting and cryo-cooling.

### Data Collection, Structure Determination and Refinement

X-ray diffraction data for USP5^171–290^ co-crystals with **1**, **48** were collected at 100 K on RIGAKU FR-E X-ray generator with a RIGAKU Saturn A200 CCD detector. Diffraction data for **64** was collected at APS beamline 24-ID-E. Images were processed with HKL3000^52^ or Xia2^53^ and scaled with AIMLESS^54^. Graphics program COOT^55^ was used for model building and visualization, REFMAC^56^ for restrained refinement and MOLPROBITY^57^ for analysis of model geometry.

### Internally Quenched Fluorophore Assay

Experiments were performed in a total volume of 60 µL in 384-well black polypropylene microplates (Greiner). Fluorescence was measured using a Biotek Synergy H1 microplate reader (Biotek) at excitation and emission wavelengths of 540 and 580 nm, respectively. Ligands were prepared in 20 mM Tris pH 7.5, 125 mM NaCl, 5 mM DTT, 0.01% TritonX-100 (v/v), 1% DMSO (v/v) for a two-fold dilution titration series. 125 nM USP5^1–835^ was added and the plate was incubated at room temperature for 1 hour before the addition of 200 nM IQF Ub2K48 (LifeSensors, DU4802). Following a 1-minute centrifugation at 250 g, fluorescence readings were immediately taken for ten minutes. The data were analyzed with GraphPad Prism 8.4.2.

### Chemistry

All reagents were purchased from commercial vendors and used without further purification. Volatiles were removed under reduced pressure by rotary evaporation or by using the V-10 solvent evaporator system by Biotage^®^. Very high boiling point (6000 rpm, 0 mbar, 56 °C), mixed volatile (7000 rpm, 30 mbar, 36 °C) and volatile (6000 rpm, 30 mbar, 36 °C) methods were used to evaporate solvents. The yields given refer to chromatographically purified and spectroscopically pure compounds. Compounds were purified using a Biotage Isolera One system by normal phase chromatography using Biotage^®^ SNAP KP- Sil or Sfär Silica D columns (Part No.: FSKO-1107/FSRD-0445) or by reverse-phase chromatography using Biotage^®^ SNAP KP-C18-HS or Sfär C18 D (Part No.: FSLO-1118/FSUD-040). If additional purification was required, compounds were purified by solid phase extraction (SPE) using Biotage Isolute Flash SCX-2 cation exchange cartridges (Part No.: 532-0050-C and 456-0200-D). Products were washed with 2 cartridge volumes of MeOH and eluted with a solution of MeOH and NH_4_OH (9:1 v/v). Final compounds were dried using the Labconco^TM^ Benchtop FreeZone^TM^ Freeze-Dry System (4.5 L Model). ^1^H and proton-decoupled ^19^F NMRs were recorded on a Bruker Avance-III 500 MHz spectrometer at ambient temperature. An Agilent D500 was used for ^13^C NMRs data collection. Residual protons of CDCl_3_, DMSO-*d*_6_ and CD_3_OD solvents were used as internal references. Spectral data are reported as follows: chemical shift (δ in ppm), multiplicity (br = broad, s = singlet, d = doublet, dd = doublet of doublets, m = multiplet), coupling constants (*J* in Hz) and proton integration. Compound purity was determined by UV absorbance at 254 nm during tandem liquid chromatography/mass spectrometry (LCMS) using a Waters Acquity separations module. All compounds have ≥95% purity, with exceptions for compounds **5** (90%), and **S22** (93%) as determined using this method. Low resolution mass spectrometry (LRMS) was conducted in positive/negative ion mode using a Waters Acquity SQD mass spectrometer (electrospray ionization source) fitted with a PDA detector. Mobile phase A consisted of 0.1% formic acid in water, while mobile phase B consisted of 0.1% formic acid in acetonitrile. One of three types of columns were used: Method 1: Acquity UPLC CSH C18 (2.1 x 50 mm, 130 Å, 1.7 µm. Part No. 186005296), Method 2: Acquity UPLC BEH C8 (2.1 x 50 mm, 130 Å, 1.7 µm. Part No. 186002877) or Method 3: Acquity UPLC HSS T3 (2.1 x 50 mm, 100 Å, 1.8 µm. Part No. 186003538). All were used with column temperature maintained at 25 °C. High resolution mass spectrometry (HRMS) was conducted using a Waters Synapt G2-S quadrupole-time-of-flight (QTOF) hybrid mass spectrometer system coupled with an Acquity ultra-performance liquid chromatography (UPLC) system (**Table S6**). Chromatographic separations were carried out on an Acquity UPLC CSH C18 (2.1 x 50 mm, 130 Å, 1.7 µm. Part No. 186005296, Method 1), Acquity UPLC BEH C8 (2.1 x 50 mm, 130 Å, 1.7 µm. Part No. 186002877) or Acquity UPLC HSS T3 (2.1 x 50 mm, 100 Å, 1.8 µm. Part No. 186003538). The mobile phase was 0.1% formic acid in water (solvent A) and 0.1% formic acid in acetonitrile (solvent B). Leucine Enkephalin was used as lock mass. MassLynx 4.1 was used for data analysis.

**Scheme 1.**
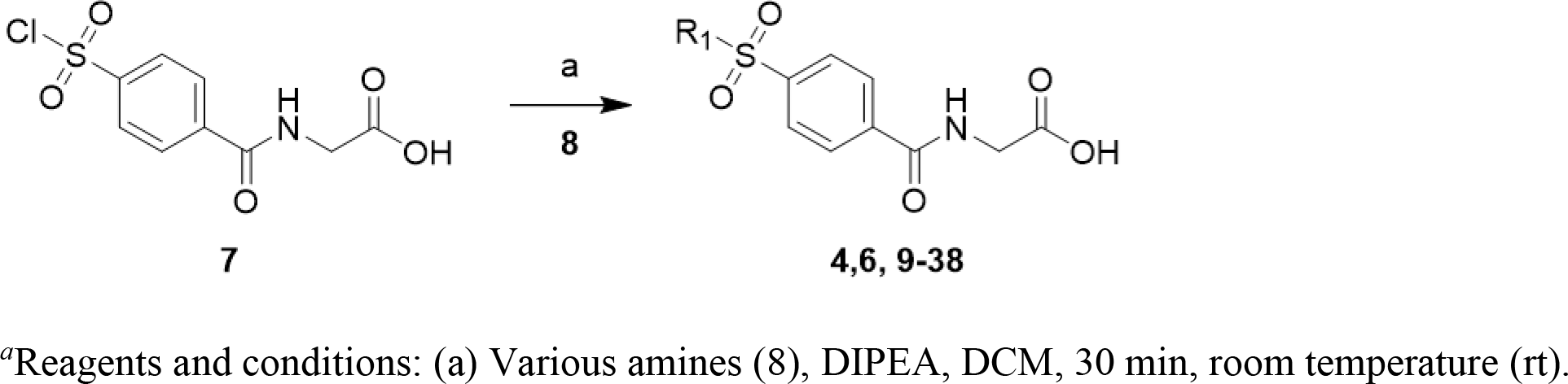
Synthesis of Compounds 4, 6 and 9-38*^a^*.

### (4-((4-phenylpiperidin-1-yl)sulfonyl)benzoyl)glycine (4)

A solution of 2-{[4-(chlorosulfonyl)phenyl]formamido}acetic acid (**7**, 54 mg, 0.194 mmol), 4-phenyl-piperidine (**8**, 47 mg, 0.292 mmol) and DIPEA (68 µL, 0.389 mmol) in DCM (2 mL) was stirred for 30 min at room temperature (rt), after which time LC analysis indicated total consumption of the starting material and only one major peak. The volatiles were removed by evaporation to yield a yellow solid to which DCM (10 mL) and aqueous HCl 10% (10 mL) were added. Upon combination of organic and aqueous solutions, an insoluble precipitate appears, which floats between the two layers. The organic phase alongside the insoluble precipitate was collected and washed with (2x 10% HCl). The crude material was purified by reverse-phase chromatography (2-95% ACN (0.1% formic acid) in water (0.1% formic acid)). After evaporation of the desired fractions the product was suspended in water and freeze dried for 2 days. White powder; yield 61% (48 mg, 0.119 mmol). ^1^H NMR (500 MHz, DMSO) δ 9.12 (t, *J* = 5.8 Hz, 1H), 8.12 (d, *J* = 8.4 Hz, 2H), 7.89 (d, *J* = 8.4 Hz, 2H), 7.30 – 7.24 (m, 2H), 7.21 – 7.15 (m, 3H), 3.96 (d, *J* = 5.9 Hz, 2H), 3.80 (d, *J* = 11.7 Hz, 2H), 2.37 – 2.31 (m, 2H), 1.81 (d, *J* = 11.2 Hz, 2H), 1.66 (qd, *J* = 12.7, 3.9 Hz, 2H). LC: Method 1, t_R_ = 1.68 min. MS (ESI): *m/z* = 401.56 [M - 1]^-^; HRMS (ESI) for C20H23N2O5S [M + H]^+^: *m/z* = calcd, 403.1328; found, 403.1326.

### (4-((4-(pyrrolidin-1-yl)piperidin-1-yl)sulfonyl)benzoyl)glycine (6)

A solution of 2-{[4-(chlorosulfonyl)phenyl]formamido}acetic acid (52 mg, 0.188 mmol), 4-(1-pyrrolidinyl)piperidine (44 mg, 0.282 mmol)) and DIPEA (66 µL, 0.377 mmol) in DCM (2 mL) was stirred for 30 min at rt, after which time LC analysis indicated total consumption of the starting material and only one major peak. The volatiles were removed by evaporation to yield a yellow solid. The crude material was purified by reverse-phase chromatography (2-95% ACN (0.1% formic acid) in water (0.1% formic acid)). After evaporation of the desired fractions the product was suspended in water and freeze dried for 2 days. White powder; yield 39% (29 mg, 0.073 mmol). Obtained as a formic acid salt. ^1^H NMR (500 MHz, DMSO) δ 9.01 (t, *J* = 5.7 Hz, 1H), 8.08 (d, *J* = 8.5 Hz, 2H), 7.85 (d, *J* = 8.5 Hz, 2H), 3.90 (d, *J* = 5.8 Hz, 2H), 3.49 (d, *J* = 12.0 Hz, 4H), 2.46 (s, 4H), 2.05 (t, *J* = 9.6 Hz, 1H), 1.85 (d, *J* = 10.3 Hz, 2H), 1.63 (s, 4H), 1.47 – 1.39 (m, 2H). LC: Method 1, t_R_ = 0.94 min. MS (ESI): *m/z* = 394.56 [M - 1]^-^; HRMS (ESI) for C_18_H_26_N_3_O_5_S [M + H]^+^: *m/z* = calcd, 396.1593; found, 396.1584.

### (4-(isoindolin-2-ylsulfonyl)benzoyl)glycine (9)

Compound **9** was prepared as described for **4**, using 2,3-dihydro-1*H*-isoindole hydrochloride (56 mg, 0.360 mmol, 2 equiv) as amine. Off-white powder; yield 54% (35 mg, 0.096 mmol). ^1^H NMR (500 MHz, DMSO) δ 12.63 (s, 1H), 9.05 (t, *J* = 5.9 Hz, 1H), 8.04 (d, *J* = 8.6 Hz, 2H), 7.98 (d, *J* = 8.6 Hz, 2H), 7.24 (d, *J* = 1.0 Hz, 4H), 4.61 (s, 4H), 3.92 (d, *J* = 5.9 Hz, 2H). LC: Method 1, t_R_ = 1.47 min. MS (ESI): *m/z* = 359.27 [M - H]^-^; HRMS (ESI) for C_17_H_17_N_2_O_5_S [M+ H]^+^: *m/z* = calcd, 361.0858; found, 361.0860.

### (4-((3,4-dihydroisoquinolin-2(1H)-yl)sulfonyl)benzoyl)glycine (10)

Compound **10** was prepared as described for **4**, using 1,2,3,4-tetrahydroisoquinoline (52 mg, 0.388 mmol, 2 equiv) as amine. White powder; yield 63% (44 mg, 0.117 mmol).^1^H NMR (500 MHz, DMSO) δ 12.66 (s, 1H), 9.08 (t, *J* = 5.9 Hz, 1H), 8.07 (d, *J* = 8.6 Hz, 2H), 7.94 (d, *J* = 8.6 Hz, 2H), 7.14 (dd, *J* = 5.7, 3.7 Hz, 3H), 7.11 (dt, *J* = 7.9, 3.9 Hz, 1H), 4.24 (s, 2H), 3.94 (d, *J* = 5.9 Hz, 2H), 3.34 (t, *J* = 6.0 Hz, 2H), 2.85 (t, *J* = 5.9 Hz, 2H); LC: Method 1, t_R_ = 1.54 min. MS (ESI): *m/z* = 373.26 [M - H]^-^; HRMS (ESI) for C_18_H_19_N_2_O_5_S [M + H]^+^: *m/z* = calcd, 375.1015; found, 375.1021.

### (4-((3-phenylazetidin-1-yl)sulfonyl)benzoyl)glycine (11)

Compound **11** was prepared as described for **4**, using **7** (100 mg, 0.360 mmol) and 3-phenylazetidine, HCl (92 mg, 0.540 mmol, 2 equiv) as amine. White powder; yield 36% (48 mg, 0.129 mmol). ^1^H NMR (500 MHz, DMSO) δ 9.18 (t, *J* = 5.8 Hz, 1H), 8.19 (d, *J* = 8.4 Hz, 2H), 8.01 (d, *J* = 8.4 Hz, 2H), 7.24 – 7.15 (m, 3H), 6.89 – 6.84 (m, 2H), 4.16 (dd, *J* = 10.6, 5.0 Hz, 2H), 4.00 – 3.97 (m, 2H), 3.75 – 3.65 (m, 3H); LC: Method 1, t_R_= 1.63 min. MS (ESI): *m/z*=375.48 [M + H]^+^; HRMS (ESI) for C_18_H_19_N_2_O_5_S [M + H]^+^: *m/z* = calcd, 375.1015; found, 375.1008.

### (4-((3-phenylpyrrolidin-1-yl)sulfonyl)benzoyl)glycine (12)

Compound **12** was prepared as described for **4**, using **7** (100 mg, 0.360 mmol) and 3-phenylpyrrolidine (0.105 mL, 0.720 mmol, 2 equiv) as amine. White powder; yield 56% (79 mg, 0.202 mmol). ^1^H NMR (500 MHz, DMSO) δ 9.10 (t, *J* = 5.7 Hz, 1H), 8.10 (d, *J* = 8.4 Hz, 2H), 7.97 (d, *J* = 8.4 Hz, 2H), 7.26 (t, *J* = 7.3 Hz, 2H), 7.22 – 7.17 (m, 1H), 7.14 (d, *J* = 7.2 Hz, 2H), 3.95 (d, *J* = 5.7 Hz, 2H), 3.72 (dd, *J* = 9.6, 7.4 Hz, 1H), 3.46 (ddd, *J* = 10.4, 8.4, 3.0 Hz, 1H), 3.30 – 3.24 (m, 2H), 3.17 (dt, *J* = 16.6, 8.2 Hz, 1H), 3.09 (t, *J* = 9.5 Hz, 1H), 2.15 (dtd, *J* = 9.7, 6.8, 3.0 Hz, 1H), 1.78 (td, *J* = 18.5, 9.1 Hz, 1H); LC: Method 1, t_R_= 1.70 min. MS (ESI): *m/z*= 389.27 [M + H]^+^; HRMS (ESI) for C_19_H_21_N_2_O_5_S [M + H]^+^: *m/z* = calcd, 389.1171; found, 389.1171.

### (4-((4-phenylazepan-1-yl)sulfonyl)benzoyl)glycine (13)

Compound **13** was prepared as described for **4**, using 4-phenylazepane, HCl (57 mg, 0.270 mmol) as amine. Off-white powder; yield 46% (35 mg, 0.083 mmol).^1^H NMR (500 MHz, DMSO) δ 9.08 (t, *J* = 5.9 Hz, 1H), 8.08 (d, *J* = 8.6 Hz, 2H), 7.93 (d, *J* = 8.5 Hz, 2H), 7.26 (t, *J* = 7.5 Hz, 2H), 7.18 – 7.10 (m, 3H), 3.95 (d, *J* = 5.9 Hz, 2H), 3.55 – 3.50 (m, 2H), 3.22 – 3.17 (m, 2H), 2.67 – 2.61 (m, 1H), 1.94 – 1.87 (m, 2H), 1.79 – 1.59 (m, 4H); LC: Method 1, t_R_ = 1.75 min. MS (ESI): *m/z* = 415.40 [M - H]^-^; HRMS (ESI) for C_21_H_25_N_2_O_5_S [M + H]^+^: *m/z* = calcd, 417.1484; found, 417.1484.

### (4-((4-phenylpiperazin-1-yl)sulfonyl)benzoyl)glycine (14)

Compound **14** was prepared as described for **4**, using 1-phenylpiperazine (63 mg, 0.388 mmol) as amine. White powder; yield 74% (55 mg, 0.137 mmol); ^1^H NMR (500 MHz, DMSO) δ 9.09 (t, *J* = 5.8 Hz, 1H), 8.12 – 8.08 (m, 2H), 7.92 – 7.87 (m, 2H), 7.22 – 7.17 (m, 2H), 6.90 (dd, *J* = 8.7, 0.8 Hz, 2H), 6.80 (t, *J* = 7.3 Hz, 1H), 3.94 (d, *J* = 5.9 Hz, 2H), 3.22 – 3.18 (m, 4H), 3.08 – 3.03 (m, 4H); LC: Method 1, t_R_ = 1.56 min. MS (ESI): *m/z* = 402.23 [M - H]^-^; HRMS (ESI) for C_19_H_22_N_3_O_5_S [M + H]^+^: *m/z* = calcd, 404.1280; found, 404.1274.

### (4-((4-phenyl-3,6-dihydropyridin-1(2H)-yl)sulfonyl)benzoyl)glycine (15)

Compound **15** was prepared as described for **4**, using **7** (100 mg, 0.360 mmol) and 4-phenyl-1,2,3,6-tetrahydropyridine, HCl (106 mg, 0.540 mmol) as amine. White powder; yield 35% (50 mg, 0.125 mmol); ^1^H NMR (500 MHz, DMSO) δ 9.10 (t, *J* = 5.9 Hz, 1H), 8.09 (d, *J* = 8.5 Hz, 2H), 7.93 (d, *J* = 8.5 Hz, 2H), 7.36 (d, *J* = 7.1 Hz, 2H), 7.32 (t, *J* = 7.6 Hz, 2H), 7.27 – 7.23 (m, 1H), 6.09 (t, *J* = 3.4 Hz, 1H), 3.95 (d, *J* = 4.4 Hz, 1H), 3.94 (s, 1H), 3.72 (d, *J* = 2.8 Hz, 2H), 3.27 (t, *J* = 5.7 Hz, 2H), 2.54 (d, *J* = 1.5 Hz, 2H); LC: Method 1, t_R_= 1.78 minzz. MS (ESI): *m/z*= 401.5 [M + H]^+^; HRMS (ESI) for C_20_H_21_N_2_O_5_S [M + H]^+^: *m/z* = calcd, 401.1171; found, 401.1166.

### (4-((3-methyl-4-phenylpiperidin-1-yl)sulfonyl)benzoyl)glycine (16)

Compound **16** was prepared as described for **4**, using 3-methyl-4-phenylpiperidine, mixture of diastereomers (47 mg, 0.270 mmol) as amine. Off-white powder; yield 69% (55 mg, 0.124 mmol); ^1^H NMR (500 MHz, DMSO) δ 9.09 (q, *J* = 6.1 Hz, 1H), 8.06 (d, *J* = 8.6 Hz, 2H), 7.86 (d, *J* = 8.6 Hz, 2H), 7.27 – 7.16 (m, 3H), 7.11 – 7.05 (m, 2H), 3.95 (d, *J* = 5.9 Hz, 2H), 3.50 (dd, *J* = 10.0, 7.4 Hz, 1H), 3.20 – 3.16 (m, 1H), 2.85 (dd, *J* = 10.0, 8.4 Hz, 1H), 2.76 (dd, *J* = 9.9, 8.6 Hz, 1H), 2.67 (dd, *J* = 13.7, 5.4 Hz, 1H), 1.90 – 1.81 (m, 1H), 1.74 – 1.66 (m, 1H), 0.82 (d, *J* = 6.6 Hz, 3H) major rotamer reported. LC: Method 1, t_R_ = 1.74 min. MS (ESI): *m/z* = 415.40 [M - H]^-^; HRMS (ESI) for C_21_H_25_N_2_O_5_S [M + H]^+^: *m/z* = calcd, 417.1484; found, 417.1486.

### (4-((2-methyl-4-phenylpiperidin-1-yl)sulfonyl)benzoyl)glycine (17)

Compound **17** was prepared as described for **4**, using 2-methyl-4-phenylpiperidine, mixture of diastereomers (68.0 mg, 0.388 mmol) as amine. Light yellow powder; yield 37% (29 mg, 0.069 mmol); ^1^H NMR (500 MHz, DMSO) δ 9.10 (t, *J* = 5.7 Hz, 1H), 8.08 (d, *J* = 8.6 Hz, 2H), 7.98 (d, *J* = 8.6 Hz, 2H), 7.28 – 7.23 (m, 2H), 7.18 – 7.14 (m, 1H), 7.09 – 7.05 (m, 2H), 4.35 – 4.27 (m, 1H), 3.95 (d, *J* = 5.9 Hz, 2H), 3.82 (dq, *J* = 13.7, 2.1 Hz, 1H), 3.19 – 3.13 (m, 1H), 2.95 – 2.84 (m, 1H), 1.69 (dd, *J* = 13.0, 2.6 Hz, 1H), 1.57 (dd, *J* = 8.9, 4.1 Hz, 2H), 1.40 – 1.31 (m, 1H), 1.12 (d, *J* = 6.9 Hz, 3H); LC: Method 1, t_R_ = 1.70 min. MS (ESI): *m/z* = 415.40 [M - H]^-^; HRMS (ESI) for C_21_H_25_N_2_O_5_S [M + H]^+^: *m/z* = calcd, 417.1484; found, 417.1488.

### *Rac*-(4-(((1*R*,3*R*,5*S*)-3-phenyl-8-azabicyclo[3.2.1]octan-8-yl)sulfonyl)benzoyl)glycine (18)

Compound **18** was prepared as described for **4**, using **7** (100 mg, 0.360 mmol) and *rac*-(1*R*,3*R*,5*S*)-3-phenyl-8-azabicyclo[3.2.1]octane, HCl (120 mg, 0.540 mmol) as amine. Yellow powder; yield 15% (24 mg, 0.055 mmol); ^1^H NMR (500 MHz, DMSO) δ 9.06 (t, *J* = 5.8 Hz, 1H), 8.03 (d, *J* = 8.5 Hz, 2H), 7.99 (d, *J* = 8.5 Hz, 2H), 7.29 – 7.25 (m, 4H), 7.19 – 7.14 (m, 1H), 4.29 (s, 2H), 3.93 (d, *J* = 5.6 Hz, 2H), 2.83 (p, *J* = 9.3 Hz, 1H), 2.40 – 2.34 (m, 2H), 1.61 (dd, *J* = 13.1, 10.0 Hz, 2H), 1.49 (d, *J* = 7.4 Hz, 2H), 1.35 (dd, *J* = 8.4, 4.2 Hz, 2H); LC: Method 1, t_R_= 1.83 min. MS (ESI): *m/z*= 429.32 [M + H]^+^; HRMS (ESI) for C_22_H_25_N_2_O_5_S [M + H]^+^: *m/z* = calcd, 429.1484; found, 429.1487.

### (4-((4-phenoxypiperidin-1-yl)sulfonyl)benzoyl)glycine (19)

Compound **19** was prepared as described for **4**, using 4-phenoxy-piperidine (49 mg, 0.277 mmol) as amine. White powder; yield 79% (61 mg, 0.146 mmol); ^1^H NMR (500 MHz, DMSO) δ 9.12 (t, *J* = 5.8 Hz, 1H), 8.11 (d, *J* = 8.6 Hz, 2H), 7.89 (d, *J* = 8.6 Hz, 2H), 7.22 (dd, *J* = 8.5, 7.5 Hz, 2H), 6.90 – 6.85 (m, 3H), 4.47 – 4.41 (m, 1H), 3.96 (d, *J* = 5.9 Hz, 2H), 2.90 – 2.82 (m, 2H), 2.54 (dd, *J* = 3.8, 1.9 Hz, 2H), 2.02 – 1.93 (m, 2H), 1.70 – 1.61 (m, 2H); LC: Method 1, t_R_ = 1.67 min. MS (ESI): *m/z* = 417.28 [M - H]^-^; HRMS (ESI) for C_20_H_23_N_2_O_6_S [M + H]^+^: *m/z* = calcd, 419.1277; found, 419.1278.

### (4-((4-(2-oxoindolin-1-yl)piperidin-1-yl)sulfonyl)benzoyl)glycine (20)

Compound **20** was prepared as described for **4**, using 1-piperidin-4-yl-1,3-dihydro-2*H*-indol-2-one (58.4 mg, 0.270 mmol) as amine. Off-white powder; yield 70% (59 mg, 0.126 mmol); ^1^H NMR (500 MHz, DMSO) δ 12.69 (s, 1H), 9.14 (t, *J* = 5.9 Hz, 1H), 8.17 – 8.13 (m, 2H), 7.96 – 7.92 (m, 2H), 7.22 (dd, *J* = 7.3, 0.7 Hz, 1H), 7.15 (td, *J* = 7.8, 0.7 Hz, 1H), 6.95 (td, *J* = 7.8, 0.7 Hz, 1H), 6.80 (d, *J* = 8.0 Hz, 1H), 4.10 (tt, *J* = 12.0, 3.9 Hz, 1H), 3.97 (d, *J* = 5.9 Hz, 2H), 3.86 (d, *J* = 12.2 Hz, 2H), 3.50 (s, 2H), 2.56 (td, *J* = 12.5, 1.8 Hz, 2H), 2.32 (qd, *J* = 12.9, 4.3 Hz, 2H), 1.65 (d, *J* = 10.1 Hz, 2H); LC: Method 1, t_R_ = 1.47 min. MS (ESI): *m/z* = 456.41 [M - H]^-^; HRMS (ESI) for C_22_H_24_N_3_O_6_S [M + H]^+^: *m/z* = calcd, 458.1386; found, 458.1384.

### (4-((3*H*-spiro[isobenzofuran-1,4’-piperidin]-1’-yl)sulfonyl)benzoyl)glycine (21)

Compound **21** was prepared as described for **4**, using 3*H*-spiro[2-benzofuran-1,4’-piperidine] (86 mg, 0.382 mmol) as amine.

White powder; yield 53% (43 mg, 0.101 mmol);^1^H NMR (500 MHz, DMSO) δ 9.13 (t, *J* = 5.8 Hz, 1H), 8.11 (d, *J* = 8.4 Hz, 2H), 7.90 (d, *J* = 8.4 Hz, 2H), 7.29 – 7.23 (m, 4H), 4.87 (s, 2H), 3.96 (d, *J* = 5.8 Hz, 2H), 3.68 (d, *J* = 11.1 Hz, 2H), 2.02 (td, *J* = 13.3, 4.6 Hz, 2H), 1.67 (d, *J* = 12.6 Hz, 2H); LC: Method 1, t_R_ = 1.61 min. MS (ESI): *m/z* = 429.47 [M - H]^-^; HRMS (ESI) for C_21_H_23_N_2_O_6_S [M + H]^+^: *m/z* = calcd, 431.1277; found, 431.1264.

### (4-((4-(dimethylamino)piperidin-1-yl)sulfonyl)benzoyl)glycine (22)

Compound **22** was prepared as described for **6**, using 4-(dimethylamino)piperidine (36 mg, 0.277 mmol) as amine. Off-white powder; yield 47% (36 mg, 0.082 mmol); obtained as a formic acid salt; ^1^H NMR (500 MHz, DMSO) δ 9.09 (t, *J* = 5.9 Hz, 1H), 8.09 (d, *J* = 8.6 Hz, 2H), 7.86 (d, *J* = 8.6 Hz, 2H), 3.95 (d, *J* = 5.9 Hz, 2H), 3.67 (d, *J* = 12.0 Hz, 2H), 2.52 (d, *J* = 1.8 Hz, 2H), 2.33 (s, 6H), 1.85 (d, *J* = 11.5 Hz, 2H), 1.50 (qd, *J* = 12.3, 3.9 Hz, 2H); LC: Method 1, t_R_ = 0.92 min. MS (ESI): *m/z* = 368.30 [M - H]^-^; HRMS (ESI) for C_16_H_24_N_3_O_5_S [M + H]^+^: *m/z* = calcd, 370.1437; found, 370.1434.

### (4-((4-cyclopentylpiperidin-1-yl)sulfonyl)benzoyl)glycine (23)

Compound **23** was prepared as described for **4**, 4-cyclopentylpiperidine hydrochloride (50 mg, 0.264 mmol) as amine. White powder; yield 53% (37 mg, 0.093 mmol); ^1^H NMR (500 MHz, DMSO) δ 9.07 (t, *J* = 5.8 Hz, 1H), 8.08 (d, *J* = 8.5 Hz, 2H), 7.83 (d, *J* = 8.4 Hz, 2H), 3.94 (d, *J* = 5.7 Hz, 2H), 3.65 (d, *J* = 11.7 Hz, 2H), 2.18 (td, *J* = 11.7, 1.8 Hz, 2H), 1.71 (d, *J* = 11.5 Hz, 2H), 1.68 – 1.61 (m, 2H), 1.56 – 1.47 (m, 2H), 1.47 – 1.38 (m, 3H), 1.15 (qd, *J* = 12.5, 3.9 Hz, 2H), 1.05 – 0.93 (m, 3H); LC: Method 1, t_R_= 1.97 min. MS (ESI): *m/z*= 395.54 [M + H]^+^; HRMS ESI) for C_19_H_27_N_2_O_5_S [M + H]^+^: *m/z* = calcd, 395.1641; found, 395.1638.

### (4-((4-morpholinopiperidin-1-yl)sulfonyl)benzoyl)glycine (24)

Compound **24** was prepared as described for **6**, using 4-(piperidin-4-yl)-morpholine (46 mg, 0.270 mmol) as amine. Off-white powder; yield 62% (51 mg, 0.112 mmol); obtained as a formic acid salt; ^1^H NMR (500 MHz, DMSO) δ 9.10 (t, *J* = 5.9 Hz, 1H), 8.08 (d, *J* = 8.5 Hz, 2H), 7.85 (d, *J* = 8.5 Hz, 2H), 3.95 (d, *J* = 5.9 Hz, 2H), 3.66 (d, *J* = 12.0 Hz, 2H), 3.52 – 3.50 (m, 4H), 2.40 – 2.37 (m, 4H), 2.29 (td, *J* = 11.9, 9.9 Hz, 2H), 2.14 (tt, *J* = 10.9 Hz, 1H), 1.79 (d, *J* = 11.2 Hz, 2H), 1.41 (qd, *J* = 12.4, 3.9 Hz, 2H); LC: Method 1, t_R_ = 0.92 min. MS (ESI): *m/z* = 410.36 [M - H]^-^; HRMS (ESI) for C_18_H_26_N_3_O_6_S [M + H]^+^: *m/z* = calcd, 412.1542; found, 412.1550.

### (4-((4-(4-methylpiperazin-1-yl)piperidin-1-yl)sulfonyl)benzoyl)glycine (25)

Compound **25** was prepared as described for **6**, using 1-methyl-4-(piperidin-4-yl)piperazine (49.5 mg, 0.270 mmol) as amine. Off-white powder; yield 59% (55 mg, 0.106 mmol); obtained as a 2X formic acid salt; ^1^H NMR (500 MHz, DMSO) δ 9.03 (t, *J* = 5.8 Hz, 1H), 8.08 (d, *J* = 8.6 Hz, 2H), 7.84 (d, *J* = 8.5 Hz, 2H), 3.92 (d, *J* = 5.8 Hz, 2H), 3.65 (d, *J* = 11.9 Hz, 4H), 2.41 (s, 4H), 2.34 (s, 2H), 2.28 (td, *J* = 11.9, 10.0 Hz, 2H), 2.16 (s, 3H), 1.75 (d, *J* = 11.0 Hz, 2H), 1.41 (qd, *J* = 12.4, 3.9 Hz, 2H); LC: Method 1, t_R_ = 0.88 min. MS (ESI): *m/z* = 423.38 [M - H]^-^; HRMS (ESI) for C_19_H_29_N_4_O_5_S [M + H]^+^: *m/z* = calcd, 425.1859; found, 425.1852.

### (4-(N-(1-methylpiperidin-4-yl)sulfamoyl)benzoyl)glycine (26)

Compound **26** was prepared as described for **6**, using 4-amino-1-methylpiperidine (32 mg, 0.277 mmol) as amine. White powder; yield 22% (16 mg, 0.040 mmol); obtained as a formic acid salt; ^1^H NMR (500 MHz, DMSO) δ 9.06 (t, *J* = 5.9 Hz, 1H), 8.04 (d, *J* = 8.6 Hz, 2H), 8.00 (d, *J* = 6.6 Hz, 1H), 7.92 (d, *J* = 8.6 Hz, 2H), 3.95 (d, *J* = 5.9 Hz, 2H), 3.01 (s, 2H), 2.54 (dd, *J* = 3.8, 2.0 Hz, 2H), 1.67 (d, *J* = 11.0 Hz, 2H), 1.50 (d, *J* = 10.4 Hz, 2H); LC: Method 1, t_R_ = 0.88 min. MS (ESI): *m/z* = 354.23 [M - H]^-^; HRMS (ESI) for C_15_H_22_N_3_O_5_S [M + H]^+^: *m/z* = calcd, 356.1280; found, 356.1282.

### (4-((9-methyl-3,9-diazaspiro[5.5]undecan-3-yl)sulfonyl)benzoyl)glycine (27)

Compound **27** was prepared as described for **6**, using 3-methyl-3,9-diazaspiro[5.5]undecane (46 mg, 0.270 mmol) as amine. Off-white powder; yield 67% (55 mg, 0.120 mmol); obtained as a formic acid salt; ^1^H NMR (500 MHz, DMSO) δ 8.93 (t, *J* = 5.6 Hz, 1H), 8.07 (d, *J* = 8.5 Hz, 2H), 7.84 (d, *J* = 8.5 Hz, 2H), 3.87 (d, *J* = 5.7 Hz, 2H), 2.94 – 2.90 (m, 4H), 2.37 (s, 4H), 2.22 (s, 3H), 1.49 – 1.44 (m, 4H), 1.34 – 1.30 (m, 4H); LC: Method 1, t_R_ = 1.00 min. MS (ESI): *m/z* = 408.25 [M - H]^-^; HRMS (ESI) for C_19_H_28_N_3_O_5_S [M + H]^+^: *m/z* = calcd, 410.1750; found, 410.1748.

### (4-((4-(*p*-tolyl)piperidin-1-yl)sulfonyl)benzoyl)glycine (28)

Compound **28** was prepared as described for **4**, using 4-(4-methylphenyl)piperidine, HCl (57 mg, 0.270 mmol) as amine. Off-white powder; yield 75% (60 mg, 0.135 mmol); ^1^H NMR (500 MHz, DMSO) δ 9.10 (t, *J* = 5.8 Hz, 1H), 8.11 (d, *J* = 8.6 Hz, 2H), 7.88 (d, *J* = 8.5 Hz, 2H), 7.09 – 7.03 (m, 4H), 3.96 (d, *J* = 5.9 Hz, 2H), 3.78 (d, *J* = 11.7 Hz, 2H), 2.43 (tt, *J* = 11.9, 3.5 Hz, 1H), 2.32 (td, *J* = 12.0, 2.2 Hz, 2H), 2.23 (s, 3H), 1.78 (d, *J* = 11.0 Hz, 2H), 1.62 (qd, *J* = 12.7, 4.0 Hz, 2H); LC: Method 1, t_R_ = 1.79 min. MS (ESI): *m/z* = 415.40 [M - H]^-^; HRMS (ESI) for C_21_H_25_N_2_O_5_S [M + H]^+^: *m/z* = calcd, 417.1484; found, 417.1475.

### (4-((4-(4-methoxyphenyl)piperidin-1-yl)sulfonyl)benzoyl)glycine (29)

Compound **29** was prepared as described for **4**, using 4-(4-methoxyphenyl)piperidine (52 mg, 0.270 mmol) as amine. Off-white powder; yield 75% (59 mg, 0.135 mmol); ^1^H NMR (500 MHz, DMSO) δ 9.11 (t, *J* = 5.9 Hz, 1H), 8.11 (d, *J* = 8.6 Hz, 2H), 7.88 (d, *J* = 8.6 Hz, 2H), 7.08 (d, *J* = 8.7 Hz, 2H), 6.83 (d, *J* = 8.8 Hz, 2H), 3.96 (d, *J* = 5.9 Hz, 2H), 3.78 (d, *J* = 11.7 Hz, 2H), 3.70 (s, 3H), 2.42 (tt, *J* = 11.9, 3.4 Hz, 1H), 2.33 (td, *J* = 11.8, 1.9 Hz, 2H), 1.77 (d, *J* = 11.0 Hz, 2H), 1.61 (qd, *J* = 12.7, 4.0 Hz, 2H); LC: Method 1, t_R_ = 1.67 min. MS (ESI): *m/z* = 431.43 [M - H]^-^; HRMS (ESI) for C_21_H_25_N_2_O_6_S [M + H]^+^: *m/z* = calcd, 433.1433; found, 433.1430.

### (4-((4-(4-fluorophenyl)piperidin-1-yl)sulfonyl)benzoyl)glycine (30)

Compound **30** was prepared as described for **4**, using 4-(4-fluorophenyl)-piperidine (32 mg, 0.180 mmol) as amine. Off-white powder; yield 77% (59 mg, 0.138 mmol); ^1^H NMR (500 MHz, DMSO) δ 9.12 (t, *J* = 5.9 Hz, 1H), 8.11 (d, *J* = 8.6 Hz, 2H), 7.89 (d, *J* = 8.6 Hz, 2H), 7.24 – 7.20 (m, 2H), 7.11 – 7.06 (m, 2H), 3.96 (d, *J* = 5.9 Hz, 2H), 3.79 (d, *J* = 11.7 Hz, 2H), 2.33 (td, *J* = 12.0, 2.2 Hz, 2H), 1.79 (d, *J* = 11.0 Hz, 2H), 1.64 (qd, *J* = 12.7, 3.9 Hz, 2H); ^19^F NMR (471 MHz, DMSO) δ -116.85 (s); LC: Method 1, t_R_ = 1.72 min. MS (ESI): *m/z* = 419.39 [M - H]^-^; HRMS (ESI) for C_20_H_22_FN_2_O_5_S [M + H]^+^: *m/z* = calcd, 421.1233; found, 421.1226.

### (4-((4-(4-chlorophenyl)piperidin-1-yl)sulfonyl)benzoyl)glycine (31)

Compound **31** was prepared as described for **4**, using 4-(4-chlorophenyl)piperidine, HCl (63 mg, 0.270 mmol) as amine. Off-white powder; yield 71% (58 mg, 0.128 mmol); ^1^H NMR (500 MHz, DMSO) δ 9.10 (t, *J* = 5.9 Hz, 1H), 8.11 (d, *J* = 8.6 Hz, 2H), 7.89 (d, *J* = 8.6 Hz, 2H), 7.32 (d, *J* = 8.5 Hz, 2H), 7.21 (d, *J* = 8.5 Hz, 2H), 3.96 (d, *J* = 5.9 Hz, 2H), 3.79 (d, *J* = 11.7 Hz, 2H), 2.33 (td, *J* = 12.0, 2.1 Hz, 2H), 1.79 (d, *J* = 11.1 Hz, 2H), 1.64 (qd, *J* = 12.7, 3.9 Hz, 2H); LC: Method 1, t_R_ = 1.82 min. MS (ESI): *m/z* = 435.42 [M - H]^-^; HRMS (ESI) for C_20_H_22_ClN_2_O_5_S [M + H]^+^: *m/z* = calcd, 437.0938; found, 437.0928.

### (4-((4-(4-bromophenyl)piperidin-1-yl)sulfonyl)benzoyl)glycine (32)

Compound **32** was prepared as described for **4**, using 4-(4-bromophenyl)piperidine (93 mg, 0.389 mmol) as amine. White powder; yield 56% (56 mg, 0.108 mmol); ^1^H NMR (500 MHz, CD_3_CN) δ 8.01 (d, *J* = 8.5 Hz, 2H), 7.87 (d, *J* = 8.5 Hz, 2H), 7.52 (s, 1H), 7.44 (d, *J* = 8.5 Hz, 2H), 7.12 (d, *J* = 8.4 Hz, 2H), 4.07 (d, *J* = 5.9 Hz, 2H), 3.86 (dd, *J* = 9.7, 2.1 Hz, 2H), 2.47 (tt, *J* = 12.2, 3.6 Hz, 1H), 2.38 (td, *J* = 12.1, 2.5 Hz, 2H), 1.84 (d, *J* = 11.8 Hz, 2H), 1.70 (qd, *J* = 12.7, 4.0 Hz, 2H); LC: Method 1, t_R_ = 1.82 min. MS (ESI): *m/z* = 479.36 [M - 1]^-^, 481.46 [M - 1]^-^ +2; HRMS (ESI) for C_20_H_22_BrN_2_O_5_S [M + H]^+^: *m/z* = calcd, 481.0433; found, 481.0434.

### (4-((4-(3-(trifluoromethyl)phenyl)piperidin-1-yl)sulfonyl)benzoyl)glycine (33)

Compound **33** was prepared as described for **4**, using 4-(3-trifluoromethylphenyl)piperidine, HCl (103 mg, 0.389 mmol) as amine. White powder; yield 76% (66 mg, 0.141 mmol); ^1^H NMR (500 MHz, DMSO) δ 8.53 (s, 1H), 7.56 (d, *J* = 8.4 Hz, 2H), 7.34 (d, *J* = 8.4 Hz, 2H), 7.01 – 6.94 (m, 4H), 3.38 (d, *J* = 5.8 Hz, 2H), 3.25 (d, *J* = 11.6 Hz, 2H), 2.12 – 2.04 (m, 1H), 1.77 (td, *J* = 10.3, 2.0 Hz, 2H), 1.28 (d, *J* = 10.9 Hz, 2H), 1.17 (qd, *J* = 12.6, 3.8 Hz, 2H); ^19^F NMR (471 MHz, CD_3_CN) δ -63.04 (s); LC: Method 1, t_R_ = 1.81 min. MS (ESI): *m/z* = 469.35 [M - H]^-^; HRMS (ESI) for C_21_H_22_F_3_N_2_O_5_S [M + H]^+^: *m/z* = calcd, 471.1202; found, 471.1192.

### (4-((4-(*m*-tolyl)piperidin-1-yl)sulfonyl)benzoyl)glycine (34)

Compound **34** was prepared as described for **4**, using 4-(3-methylphenyl)piperidine, HCl (57 mg, 0.270 mmol) as amine. Off-white powder; yield 42% (33 mg, 0.076mmol); ^1^H NMR (500 MHz, DMSO) δ 9.11 (t, *J* = 5.9 Hz, 1H), 8.11 (d, *J* = 8.5 Hz, 2H), 7.89 (d, *J* = 8.5 Hz, 2H), 7.15 (t, *J* = 7.5 Hz, 1H), 6.98 (d, *J* = 8.3 Hz, 1H), 6.96 (d, *J* = 9.1 Hz, 2H), 3.96 (d, *J* = 5.9 Hz, 2H), 3.79 (d, *J* = 11.7 Hz, 2H), 2.43 (tt, *J* = 12.1, 3.4 Hz, 1H), 2.33 (td, *J* = 12.0, 2.2 Hz, 2H), 2.25 (s, 3H), 1.78 (d, *J* = 11.0 Hz, 2H), 1.64 (qd, *J* = 12.7, 3.9 Hz, 2H); LC: Method 1, t_R_ = 1.79 min. MS (ESI): *m/z* = 415.33 [M - H]^-^; HRMS (ESI) for C_21_H_25_N_2_O_5_S [M + H]^+^: *m/z* = calcd, 417.1484; found, 417.1480.

### (4-((4-(3-methoxyphenyl)piperidin-1-yl)sulfonyl)benzoyl)glycine (35)

Compound **35** was prepared as described for **4**, using 4-(3-methoxyphenyl)piperidine (52 mg, 0.270 mmol) as amine. Off-white powder; yield 80% (63 mg, 0.145 mmol); ^1^H NMR (500 MHz, DMSO) δ 9.11 (t, *J* = 5.9 Hz, 1H), 8.11 (d, *J* = 8.5 Hz, 2H), 7.89 (d, *J* = 8.5 Hz, 2H), 7.18 (t, *J* = 7.8 Hz, 1H), 6.74 (dd, *J* = 7.9, 2.0 Hz, 2H), 6.72 (d, *J* = 1.6 Hz, 1H), 3.96 (d, *J* = 5.9 Hz, 2H), 3.79 (d, *J* = 11.6 Hz, 2H), 3.70 (s, 3H), 2.46 (tt, *J* = 12.1, 3.5 Hz, 1H), 2.34 (td, *J* = 11.6, 1.8 Hz, 2H), 1.79 (d, *J* = 11.0 Hz, 2H), 1.66 (qd, *J* = 12.6, 3.9 Hz, 2H); LC: Method 1, t_R_ = 1.68 min. MS (ESI): *m/z* = 431.28 [M - H]^-^; HRMS (ESI) for C_21_H_25_N_2_O_6_S [M + H]^+^: *m/z* = calcd, 433.1433; found, 433.1444.

### (4-((4-(3-fluorophenyl)piperidin-1-yl)sulfonyl)benzoyl)glycine (36)

Compound **36** was prepared as described for **4**, using 4-(3-fluorophenyl)piperidine (48 mg, 0.270 mmol) as amine. Off-white powder; yield 74% (56 mg, 0.134 mmol); ^1^H NMR (500 MHz, DMSO) δ 9.12 (t, *J* = 5.9 Hz, 1H), 8.11 (d, *J* = 8.6 Hz, 2H), 7.89 (d, *J* = 8.6 Hz, 2H), 7.31 (td, *J* = 8.1, 6.4 Hz, 1H), 7.06 – 6.97 (m, 3H), 3.96 (d, *J* = 5.9 Hz, 2H), 3.79 (d, *J* = 11.6 Hz, 2H), 2.32 (td, *J* = 11.9, 2.1 Hz, 2H), 1.82 (d, *J* = 11.1 Hz, 2H), 1.67 (qd, *J* = 12.6, 3.9 Hz, 2H); ^19^F NMR (471 MHz, DMSO) δ -113.27 (s); LC: Method 1, t_R_ = 1.72 min. MS (ESI): *m/z* = 419.39 [M - H]^-^; HRMS (ESI) for C_20_H_22_FN_2_O_5_S [M + H]^+^: *m/z* = calcd, 421.1233; found, 421.1227.

### (4-((4-(3-chlorophenyl)piperidin-1-yl)sulfonyl)benzoyl)glycine (37)

Compound **37** was prepared as described for **4**, using 4-(3-chlorophenyl)piperidine, HCl (63 mg, 0.270 mmol) as amine. Off-white powder; yield 65% (51 mg, 0.116 mmol); ^1^H NMR (500 MHz, DMSO) δ 9.10 (t, *J* = 5.8 Hz, 1H), 8.11 (d, *J* = 8.6 Hz, 2H), 7.89 (d, *J* = 8.5 Hz, 2H), 7.31 (dd, *J* = 8.5, 7.7 Hz, 1H), 7.25 – 7.23 (m, 2H), 7.16 (d, *J* = 7.7 Hz, 1H), 3.96 (d, *J* = 5.9 Hz, 2H), 3.79 (d, *J* = 11.7 Hz, 2H), 2.31 (td, *J* = 11.9, 2.0 Hz, 2H), 1.81 (d, *J* = 11.1 Hz, 2H), 1.67 (qd, *J* = 12.6, 3.9 Hz, 2H); LC: Method 1, t_R_ = 1.80 min. MS (ESI): *m/z* = 435.27 [M- H]^-^; HRMS (ESI) for C_20_H_22_ClN_2_O_5_S [M + H]^+^: *m/z* = calcd, 437.0938; found, 437.0937.

### (4-((4-(3-bromophenyl)piperidin-1-yl)sulfonyl)benzoyl)glycine (38)

Compound **38** was prepared as described for **4**, using 4-(3-bromophenyl)piperidine, HCl (75 mg, 0.270 mmol) as amine. Off-white powder; yield 55% (48 mg, 0.099 mmol); ^1^H NMR (500 MHz, DMSO) δ 9.10 (t, *J* = 5.9 Hz, 1H), 8.11 (d, *J* = 8.6 Hz, 2H), 7.89 (d, *J* = 8.5 Hz, 2H), 7.40 – 7.36 (m, 2H), 7.26 – 7.22 (m, 1H), 7.20 (dt, *J* = 7.8, 1.2 Hz, 1H), 3.96 (d, *J* = 5.9 Hz, 2H), 3.79 (d, *J* = 11.6 Hz, 2H), 2.31 (td, *J* = 11.8, 2.0 Hz, 2H), 1.81 (d, *J* = 11.1 Hz, 2H), 1.66 (qd, *J* = 12.6, 3.9 Hz, 2H); LC: Method 1, t_R_ = 1.83 min. MS (ESI): *m/z* = 479.28 [M- 1]^-^, 481.09 [M - 1]^-^ +2; HRMS (ESI) for C_20_H_22_BrN_2_O_5_S [M + H]^+^: *m/z* = calcd, 481.0433; found, 481.0436.

**Scheme 2.**
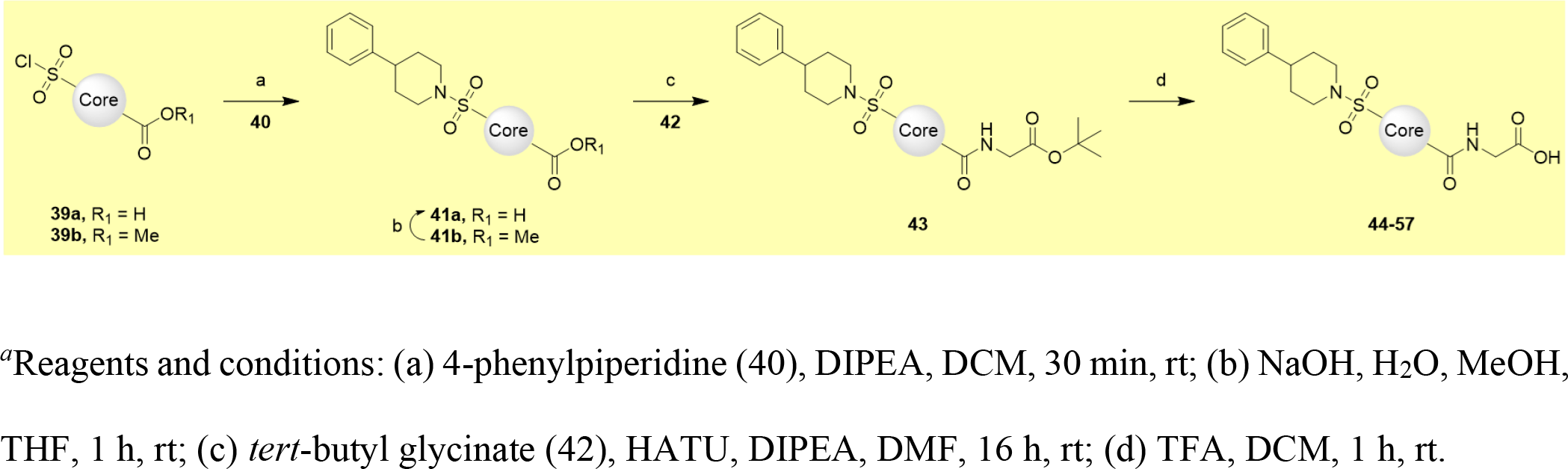
Synthesis of Compounds 44-57*^a^*.

### (3-((4-phenylpiperidin-1-yl)sulfonyl)benzoyl)glycine (44)

**Step 1,** A solution of 3-(chlorosulfonyl)benzoic acid (100 mg, 0.453 mmol), 4-phenyl-piperidine (**8**, 110 mg, 0.680 mmol) and DIPEA (0.158 mL, 0.906 mmol) in DCM (4.5 mL) was stirred for 30 min at rt, after which time LC analysis indicated total consumption of the starting material and only one major peak. The volatiles were removed by evaporation to yield a white solid to which DCM (10 mL) and aqueous HCl 10% (10 mL) were added. Upon combination of organic and aqueous solutions, an insoluble precipitate appears, which floats between the two layers. The organic phase alongside the insoluble precipitate was collected and washed with (2x 10% HCl). The crude product was suspended in water and freeze dried for 2 days to obtain **3-((4-phenylpiperidin-1-yl)sulfonyl)benzoic acid (41),** that was used in the next step without further purification. White powder; yield 71% (130 mg, 0.324 mmol). LC: Method 1, t_R_ = 1.84 min. MS (ESI): *m/z* = 344.37 [M - 1]^-^. **Step 2,** To a solution of 3-((4-phenylpiperidin-1-yl)sulfonyl)benzoic acid (50 mg, 0.145 mmol), DIPEA (0.101 mL, 0.579 mmol) and HATU (83 mg, 0.217 mmol) in *N,N*- Dimethylformamide (DMF) (2 mL) was added *tert*-butyl 2-aminoacetate, HCl (**42**, 36.4 mg, 0.217 mmol). The orange solution was stirred at rt for 24 h under N2 atmosphere, after which time water (1 mL) was added to the suspension. The volatiles were removed by evaporation to yield a white solid to which DCM (10 mL) and aqueous HCl 10% (10 mL) were added. The organic phase was collected and washed with (2x 10% HCl). The remaining DCM was evaporated to obtain a yellow solid. The protecting group was removed by dissolving the sample in DCM (2 to 3 mL) and adding TFA (0.750 mL), the solution was stirred at rt for 1 h. The volatiles were evaporated. The crude material was purified by reverse-phase chromatography (2-95% ACN (0.1% formic acid) in water (0.1% formic acid)). After evaporation of the desired fractions the product was suspended in water and freeze dried for 2 days. Off-white powder; yield 82% (50 mg, 0.118 mmol); ^1^H NMR (500 MHz, DMSO) δ 12.68 (s, 1H), 9.20 (t, *J* = 5.8 Hz, 1H), 8.25 (t, *J* = 1.6 Hz, 1H), 8.23 – 8.21 (m, 1H), 7.95 (ddd, *J* = 7.8, 1.6, 1.1 Hz, 1H), 7.80 (t, *J* = 7.8 Hz, 1H), 7.27 (dd, *J* = 8.0, 7.1 Hz, 2H), 7.19 (d, *J* = 7.3 Hz, 3H), 3.97 (d, *J* = 5.9 Hz, 2H), 3.81 (d, *J* = 11.6 Hz, 2H), 2.34 (td, *J* = 12.0, 2.0 Hz, 2H), 1.82 (d, *J* = 11.1 Hz, 2H), 1.67 (qd, *J* = 12.7, 3.9 Hz, 2H); LC: Method 1, t_R_ = 1.67 min. MS (ESI): *m/z* = 401.41 [M - H]^-^; HRMS (ESI) for C_20_H_23_N_2_O_5_S [M + H]^+^: *m/z* = calcd, 403.1328; found, 403.1317.

### (3-fluoro-4-((4-phenylpiperidin-1-yl)sulfonyl)benzoyl)glycine (45)

Compound **45** was prepared as described for **44**, using 4-(chlorosulfonyl)-3-fluorobenzoic acid (100 mg, 0.419 mmol) as a core. Off-white powder; yield 73% (44 mg, 0.100 mmol); ^1^H NMR (500 MHz, DMSO) δ 12.71 (s, 1H), 9.18 (t, *J* = 5.9 Hz, 1H), 7.97 – 7.90 (m, 3H), 7.31 – 7.26 (m, 2H), 7.21 – 7.16 (m, 3H), 3.97 (d, *J* = 5.9 Hz, 2H), 3.83 (d, *J* = 11.9 Hz, 2H), 2.64 (t, *J* = 12.2 Hz, 2H), 2.58 (tt, *J* = 12.0, 3.3 Hz, 1H), 1.84 (d, *J* = 11.2 Hz, 2H), 1.65 (qd, *J* = 12.7, 4.0 Hz, 2H); ^19^F NMR (471 MHz, DMSO) δ -107.65; LC: Method 1, t_R_ = 1.72 min. MS (ESI): *m/z* = 419.39 [M - H]^-^; HRMS (ESI) for C_20_H_22_FN_2_O_5_S [M + H]^+^: *m/z* = calcd, 421.1233; found, 421.1231.

### (3-chloro-4-((4-phenylpiperidin-1-yl)sulfonyl)benzoyl)glycine (46)

Compound **46** was prepared as described for **44**, using 3-chloro-4-(chlorosulfonyl)benzoic acid (100 mg, 0.392 mmol) as a core. Off-white powder; yield 92% (53 mg, 0.121 mmol); ^1^H NMR (500 MHz, DMSO) δ 12.72 (s, 1H), 9.21 (t, *J* = 5.9 Hz, 1H), 8.14 (d, *J* = 1.7 Hz, 1H), 8.12 (d, *J* = 8.2 Hz, 1H), 8.01 (dd, *J* = 8.2, 1.7 Hz, 1H), 7.29 (dd, *J* = 8.0, 7.1 Hz, 2H), 7.20 (d, *J* = 7.2 Hz, 3H), 3.96 (d, *J* = 5.9 Hz, 2H), 3.86 (d, *J* = 12.4 Hz, 2H), 2.84 (td, *J* = 12.4, 2.1 Hz, 2H), 2.65 (tt, *J* = 6.0, 3.5 Hz, 1H), 1.83 (d, *J* = 11.0 Hz, 2H), 1.61 (qd, *J* = 12.7, 4.1 Hz, 2H); LC: Method 1, t_R_ = 1.76 min. MS (ESI): *m/z* = 435.42 [M - H]^-^; HRMS (ESI) for C_20_H_22_ClN_2_O_5_S [M + H]^+^: *m/z* = calcd, 437.0938; found, 437.0925.

### (3-methyl-4-((4-phenylpiperidin-1-yl)sulfonyl)benzoyl)glycine (47)

Compound **47** was prepared as described for **44**, using 4-(chlorosulfonyl)-3-methylbenzoic acid (100 mg, 0.426 mmol) as a core. Off-white powder; yield 79% (48 mg, 0.110 mmol); ^1^H NMR (500 MHz, DMSO) δ 12.66 (s, 1H), 9.03 (t, *J* = 5.7 Hz, 1H), 7.94 – 7.91 (m, 2H), 7.87 (dd, *J* = 8.3, 1.4 Hz, 1H), 7.31 – 7.26 (m, 2H), 7.22 – 7.18 (m, 3H), 3.94 (d, *J* = 5.9 Hz, 2H), 3.76 (d, *J* = 12.0 Hz, 2H), 2.70 (td, *J* = 12.2, 2.1 Hz, 2H), 2.65 (s, 3H), 2.60 (tt, *J* = 12.4, 3.6 Hz, 1H), 1.83 (d, *J* = 11.0 Hz, 2H), 1.62 (qd, *J* = 12.6, 4.0 Hz, 2H); LC: Method 1, t_R_ = 1.7 min. MS (ESI): *m/z* = 415.40 [M - H]^-^; HRMS (ESI) for C_21_H_25_N_2_O_5_S [M + H]^+^: *m/z* = calcd, 417.1484; found, 417.1482.

### (2-fluoro-4-((4-phenylpiperidin-1-yl)sulfonyl)benzoyl)glycine (48)

Compound **48** was prepared as described for **44**, using 4-(chlorosulfonyl)-2-fluorobenzoic acid (100 mg, 0.419 mmol) as a core. Off-white powder; yield 84% (50 mg, 0.116 mmol); ^1^H NMR (500 MHz, DMSO) δ 13.14 (s, 1H), 9.27 (dd, *J* = 5.8, 4.3 Hz, 1H), 8.32 (t, *J* = 7.3 Hz, 1H), 8.16 – 8.11 (m, 2H), 7.70 (dd, *J* = 8.0, 7.1 Hz, 2H), 7.61 (d, *J* = 7.2 Hz, 3H), 4.39 (d, *J* = 5.9 Hz, 2H), 4.23 (d, *J* = 11.6 Hz, 2H), 2.97 – 2.94 (m, 1H), 2.83 (td, *J* = 12.0, 2.1 Hz, 2H), 2.25 (d, *J* = 11.0 Hz, 2H), 2.10 (qd, *J* = 12.7, 4.0 Hz, 2H); ^13^C NMR (126 MHz, DMSO) δ 171.14, 163.31, 159.33 (d, ^1^*J* C-F = 254.7 Hz), 145.53, 139.47 (d, ^3^*J* C-F = 6.8 Hz), 131.95, 128.84, 127.98 (d, ^2^*J* C-F = 15.0 Hz), 127.12, 126.74, 124.07 (d, ^3^*J* C-F = 3.5 Hz), 115.98 (d, ^2^*J* C-F = 25.6 Hz), 46.96, 41.77, 40.83, 32.40; ^19^F NMR (471 MHz, DMSO) δ -110.90 (s); LC: Method 1, t_R_ = 1.73 min. MS (ESI): *m/z* = 419.54 [M - H]^-^; HRMS (ESI) for C_20_H_22_FN_2_O_5_S [M + H]^+^: *m/z* = calcd, 421.1233; found, 421.1241.

### (2-chloro-4-((4-phenylpiperidin-1-yl)sulfonyl)benzoyl)glycine (49)

Compound **49** was prepared as described for **44**, using 2-chloro-4-(chlorosulfonyl)benzoic acid (100 mg, 0.392 mmol) as a core. Off-white powder; yield 94% (55 mg, 0.124 mmol); ^1^H NMR (500 MHz, DMSO) δ 12.73 (s, 1H), 8.98 (t, *J* = 5.4 Hz, 1H), 7.84 (d, *J* = 1.6 Hz, 1H), 7.82 (dd, *J* = 7.9, 1.7 Hz, 1H), 7.71 (d, *J* = 7.9 Hz, 1H), 7.28 (dd, *J* = 10.0, 5.0 Hz, 2H), 7.20 (dd, *J* = 8.0, 1.7 Hz, 3H), 3.95 (d, *J* = 5.9 Hz, 2H), 3.81 (d, *J* = 11.6 Hz, 2H), 2.39 (td, *J* = 11.6, 1.7 Hz, 2H), 1.84 (d, *J* = 11.2 Hz, 2H), 1.68 (qd, *J* = 12.6, 3.9 Hz, 2H); LC: Method 1, t_R_ = 1.74 min. MS (ESI): *m/z* = 435.34 [M - H]^-^; HRMS (ESI) for C_20_H_22_ClN_2_O_5_S [M + H]^+^: *m/z* = calcd, 437.0938; found, 437.0934.

### (2-methyl-4-((4-phenylpiperidin-1-yl)sulfonyl)benzoyl)glycine (50)

Compound **50** was prepared as described for **44**, using 4-(chlorosulfonyl)-2-methylbenzoic acid (100 mg, 0.426 mmol) as a core. Off-white powder; yield 100% (65 mg, 0.139 mmol); ^1^H NMR (500 MHz, DMSO) δ 12.67 (s, 1H), 8.80 (t, *J* = 6.0 Hz, 1H), 7.65 (dd, *J* = 7.9, 1.2 Hz, 2H), 7.59 – 7.57 (m, 1H), 7.30 – 7.26 (m, 2H), 7.19 (dd, *J* = 8.0, 1.8 Hz, 3H), 3.93 (d, *J* = 6.0 Hz, 2H), 3.79 (d, *J* = 11.6 Hz, 2H), 2.46 (s, 3H), 2.33 (td, *J* = 11.9, 2.1 Hz, 2H), 1.82 (d, *J* = 11.1 Hz, 2H), 1.68 (qd, *J* = 12.7, 4.0 Hz, 2H); LC: Method 1, t_R_ = 1.66 min. MS (ESI): *m/z* = 415.40 [M - H]^-^; HRMS (ESI) for C_21_H_25_N_2_O_5_S [M + H]^+^: *m/z* = calcd, 417.1484; found, 417.1472.

### (5-((4-phenylpiperidin-1-yl)sulfonyl)picolinoyl)glycine (51)

**Step 1,** A solution of methyl 5-(chlorosulfonyl)pyridine-2-carboxylate (50 mg, 0.212 mmol), 4-phenyl-piperidine (**8**, 51 mg, 0.318 mmol) and DIPEA (0.074 mL, 0.424 mmol) in DCM (2 mL) was stirred for 30 min at rt, after which time LC analysis indicated total consumption of the starting material and only one major peak. The volatiles were removed by evaporation to yield a white solid to which DCM (10 mL) and aqueous HCl 10% (10 mL) were added. Upon combination of organic and aqueous solutions, an insoluble precipitate appears, which floats between the two layers. The organic phase alongside the insoluble precipitate was collected and washed with (2x 10% HCl). Saponification of the crude product was carried out by stiring the solution with NaOH (1 pellet) in THF, water, MeOH. The volatiles were removed by evaporation and the aqueous solution was neutralized using formic acid. The product was purified by reverse-phase chromatography (2-95% ACN (0.1% formic acid) in water (0.1% formic acid)), the fractions containing the product were freeze dried for 2 days to obtain **5-((4-phenylpiperidin-1-yl)sulfonyl)picolinic acid.** White powder; yield 34% (25 mg, 0.072 mmol). LC: Method 1, t_R_ = 1.73 min. MS (ESI): *m/z* = 345.27 [M - 1]^-^. **Step 2,** To a solution of 5-((4-phenylpiperidin-1-yl)sulfonyl)picolinic acid (25 mg, 0.072 mmol), DIPEA (0.050 mL, 0.289 mmol) and HATU (41.2 mg, 0.108 mmol) in DMF (1 mL) was added *tert*-butyl 2-aminoacetate, HCl (**42**, 18.2 mg, 0.108 mmol). The orange solution was stirred at rt for 24 h under N2 atmosphere, after which time water (1 mL) was added to the suspension. The volatiles were removed by evaporation to yield a white solid to which DCM (10 mL) and aqueous HCl 10% (10 mL) were added. The organic phase was colected and washed with (2x 10% HCl). The remaining DCM was evaporated to obtain a yellow solid. The *tert*-butyloxycarbonyl protecting group was removed by dissolving the sample in DCM (2 to 3 mL) and adding TFA (∼0.750 mL), the solution was stirred at rt for 1 h. The volatiles were evaporated. The crude material was purified by reverse-phase chromatography (2-95% ACN (0.1% formic acid) in water (0.1% formic acid). After evaporation of the desired fractions the product was suspended in water and freeze dried for 2 days. Off-white powder; yield 85% (25 mg, 0.061 mmol); ^1^H NMR (500 MHz, DMSO) δ 12.74 (s, 1H), 9.18 (t, *J* = 6.0 Hz, 1H), 9.00 (dd, *J* = 2.2, 0.7 Hz, 1H), 8.40 (dd, *J* = 8.2, 2.3 Hz, 1H), 8.29 (dd, *J* = 8.2, 0.7 Hz, 1H), 7.30 – 7.25 (m, 2H), 7.21 – 7.16 (m, 3H), 4.00 (d, *J* = 6.0 Hz, 2H), 3.84 (d, *J* = 11.6 Hz, 2H), 2.54 (tt, *J* = 12.3, 3.6 Hz, 1H), 2.45 (td, *J* = 12.0, 2.2 Hz, 2H), 1.83 (d, *J* = 11.0 Hz, 2H), 1.68 (qd, *J* = 12.6, 3.9 Hz, 2H); ^13^C NMR (126 MHz, DMSO) δ 170.79, 162.84, 152.60, 146.82, 145.07, 137.53, 134.35, 128.40, 126.65, 126.30, 122.69, 46.31, 41.25, 40.22, 31.94; LC: Method 1, t_R_ = 1.79 min. MS (ESI): *m/z* = 402.23 [M - H]^-^; HRMS (ESI) for C_19_H_22_N_3_O_5_S [M + H]^+^: *m/z* = calcd, 404.1280; found, 404.1274.

### (5-((4-phenylpiperidin-1-yl)sulfonyl)thiophene-2-carbonyl)glycine (52)

Compound **52** was prepared as described for **44**, using 5-(chlorosulfonyl)thiophene-2-carboxylic acid (100 mg, 0.441 mmol) as a core.

Off-white powder; yield 69% (43 mg, 0.099 mmol);^1^H NMR (500 MHz, DMSO) δ 12.75 (s, 1H), 9.23 (t, *J* = 5.9 Hz, 1H), 7.92 (d, *J* = 4.0 Hz, 1H), 7.69 (d, *J* = 4.0 Hz, 1H), 7.30 – 7.27 (m, 2H), 7.20 (dd, *J* = 9.2, 4.3 Hz, 3H), 3.94 (d, *J* = 5.9 Hz, 2H), 3.77 (d, *J* = 11.6 Hz, 2H), 2.58 (tt, *J* = 12.3, 3.5 Hz, 1H), 1.86 (d, *J* = 11.1 Hz, 2H), 1.71 (qd, *J* = 12.7, 3.9 Hz, 2H); LC: Method 1, t_R_ = 1.71 min, MS (ESI): *m/z* = 407.28 [M- H]^-^; HRMS (ESI) for C_18_H_21_N_2_O_5_S_2_ [M + H]^+^: *m/z* = calcd, 409.0892; found, 409.0886.

### (3-methyl-5-((4-phenylpiperidin-1-yl)sulfonyl)thiophene-2-carbonyl)glycine (53)

Compound **53** was prepared as described for **44**, using 5-(chlorosulfonyl)-3-methylthiophene-2-carboxylic acid (50 mg, 0.208 mmol) as a core. Off-white powder; yield 71% (43 mg, 0.097 mmol); ^1^H NMR (500 MHz, DMSO) δ 12.69 (s, 1H), 8.63 (t, *J* = 5.9 Hz, 1H), 8.38 (s, 1H), 7.31 – 7.27 (m, 2H), 7.23 – 7.17 (m, 3H), 3.90 (d, *J* = 5.9 Hz, 2H), 3.77 (d, *J* = 11.8 Hz, 2H), 2.66 – 2.59 (m, 3H), 2.56 (s, 3H), 1.85 (d, *J* = 11.2 Hz, 2H), 1.65 (qd, *J* = 12.7, 4.0 Hz, 2H); LC: Method 1, t_R_ = 1.77 min. MS (ESI): *m/z* = 421.20 [M - H]^-^; HRMS (ESI) for C_19_H_23_N_2_O_5_S_2_ [M + H]^+^: *m/z* = calcd, 423.1048; found, 423.1048.

### (4-((4-phenylpiperidin-1-yl)sulfonyl)thiophene-2-carbonyl)glycine (54)

Compound **54** was prepared as described for **44**, using 4-(chlorosulfonyl)thiophene-2-carboxylic acid (100 mg, 0.441 mmol) as a core. Off-white powder; yield 71% (43 mg, 0.101 mmol); ^1^H NMR (500 MHz, DMSO) δ 9.19 (t, *J* = 5.9 Hz, 1H), 8.45 (d, *J* = 1.4 Hz, 1H), 8.10 (d, *J* = 1.4 Hz, 1H), 7.30 – 7.27 (m, 2H), 7.23 – 7.18 (m, 3H), 3.93 (d, *J* = 5.9 Hz, 2H), 3.76 (d, *J* = 11.5 Hz, 2H), 2.57 (tt, *J* = 12.3, 3.7 Hz, 1H), 2.45 (td, *J* = 12.1, 2.2 Hz, 2H), 1.85 (d, *J* = 11.3 Hz, 2H), 1.70 (qd, *J* = 12.7, 3.9 Hz, 2H); LC: Method 1, t_R_ = 1.67 min. MS (ESI): *m/z* = 407.43 [M - H]^-^; HRMS (ESI) for C_18_H_21_N_2_O_5_S_2_ [M + H]^+^: *m/z* = calcd, 409.0892; found, 409.0880.

### (5-((4-phenylpiperidin-1-yl)sulfonyl)thiophene-3-carbonyl)glycine (55)

Compound **55** was prepared as described for **44**, using 5-(chlorosulfonyl)thiophene-3-carboxylic acid (100 mg, 0.441 mmol) as a core. Off-white powder; yield 59% (37 mg, 0.084 mmol); ^1^H NMR (500 MHz, DMSO) δ 12.68 (s, 1H), 8.97 (t, *J* = 5.9 Hz, 1H), 8.58 (t, *J* = 1.7 Hz, 1H), 8.04 (d, *J* = 1.6 Hz, 1H), 7.31 – 7.27 (m, 2H), 7.24 – 7.18 (m, 3H), 3.92 (d, *J* = 5.9 Hz, 2H), 3.77 (d, *J* = 11.2 Hz, 2H), 2.57 (tt, *J* = 12.5, 3.6 Hz, 1H), 2.47 (td, *J* = 12.1, 2.4 Hz, 2H), 1.87 (d, *J* = 11.2 Hz, 2H), 1.72 (qd, *J* = 12.7, 3.9 Hz, 2H); LC: Method 1, t_R_ = 1.69 min. MS (ESI): *m/z* = 407.35 [M - H]^-^; HRMS (ESI) for C_18_H_21_N_2_O_5_S_2_ [M + H]^+^: *m/z* = calcd, 409.0892; found, 409.0883.

### (5-((4-phenylpiperidin-1-yl)sulfonyl)furan-2-carbonyl)glycine (56)

Compound **56** was prepared as described for **44**, using 5-(chlorosulfonyl)furan-2-carboxylic acid (100 mg, 0.475 mmol) as a core. Off-white powder; yield 68% (40 mg, 0.101 mmol); ^1^H NMR (500 MHz, DMSO) δ 12.75 (s, 1H), 9.00 (t, *J* = 5.9 Hz, 1H), 7.36 (d, *J* = 3.7 Hz, 1H), 7.33 (d, *J* = 3.7 Hz, 1H), 7.30 – 7.26 (m, 2H), 7.19 (d, *J* = 7.3 Hz, 3H), 3.92 (d, *J* = 6.0 Hz, 2H), 3.83 (d, *J* = 12.3 Hz, 2H), 2.78 (td, *J* = 12.3, 2.2 Hz, 2H), 2.63 (tt, *J* = 12.1, 3.3 Hz, 1H), 1.82 (d, *J* = 11.0 Hz, 2H), 1.62 (qd, *J* = 12.8, 4.1 Hz, 2H); LC: Method 1, t_R_ = 1.66 min. MS (ESI): *m/z* = 391.25 [M - H]^-^; HRMS (ESI) for C_18_H_21_N_2_O_6_S [M + H]^+^: *m/z* = calcd, 393.1120; found, 393.1121.

### (1-methyl-4-((4-phenylpiperidin-1-yl)sulfonyl)-1H-pyrrole-2-carbonyl)glycine (57)

Compound **57** was prepared as described for **44**, using 4-(chlorosulfonyl)-1-methyl-1*H*-pyrrole-2-carboxylic acid (50 mg, 0.224 mmol) as a core. Off-white powder; yield 58% (32 mg, 0.080 mmol); ^1^H NMR (500 MHz, DMSO) δ 12.61 (s, 1H), 8.69 (t, *J* = 6.0 Hz, 1H), 7.59 (d, *J* = 1.8 Hz, 1H), 7.31 – 7.26 (m, 2H), 7.23 (t, *J* = 4.2 Hz, 2H), 7.21 – 7.17 (m, 1H), 7.16 (d, *J* = 1.9 Hz, 1H), 3.90 (s, 3H), 3.85 (d, *J* = 6.0 Hz, 2H), 3.66 (d, *J* = 11.4 Hz, 2H), 2.53 (tt, *J* = 12.3, 3.4 Hz, 1H), 2.32 (td, *J* = 11.9, 2.1 Hz, 2H), 1.84 (d, *J* = 10.9 Hz, 2H), 1.71 (qd, *J* = 12.6, 3.9 Hz, 2H); LC: Method 1, t_R_ = 1.66 min. MS (ESI): *m/z* = 404.19 [M - H]^-^; HRMS (ESI) for C_19_H_24_N_3_O_5_S [M + H]^+^: *m/z* = calcd, 406.1437; found, 406.1432.

**Scheme 3.**
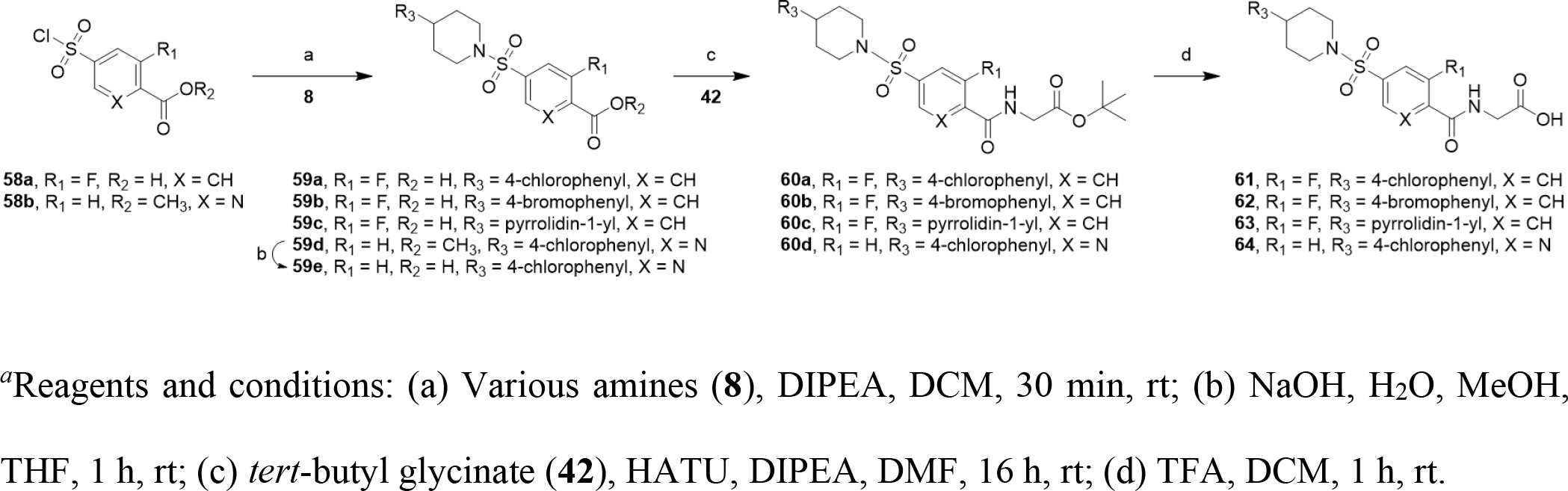
Synthesis of Compounds 61-64*^a^*.

### (4-((4-(4-chlorophenyl)piperidin-1-yl)sulfonyl)-2-fluorobenzoyl)glycine (61)

Compound **61** was prepared as described for **44**, using 4-(chlorosulfonyl)-2-fluorobenzoic acid (50 mg, 0.210 mmol) as a core. Off-white powder; yield 78% (69 mg, 0.151 mmol); ^1^H NMR (500 MHz, DMSO) δ 12.71 (s, 1H), 8.85 (dd, *J* = 5.8, 4.3 Hz, 1H), 7.89 (dd, *J* = 7.9, 6.9 Hz, 1H), 7.71 (ddd, *J* = 9.6, 8.8, 1.6 Hz, 2H), 7.33 (d, *J* = 8.5 Hz, 2H), 7.23 (d, *J* = 8.5 Hz, 2H), 3.97 (d, *J* = 5.9 Hz, 2H), 3.80 (d, *J* = 11.6 Hz, 2H), 2.55 (tt, *J* = 12.1, 3.3 Hz, 1H), 2.39 (td, *J* = 11.9, 2.0 Hz, 2H), 1.81 (d, *J* = 11.3 Hz, 2H), 1.66 (qd, *J* = 12.7, 3.9 Hz, 2H); ^13^C NMR (126 MHz, DMSO) δ 170.68, 162.84, 158.88 (d, ^1^*J*C-F = 254.6 Hz), 144.07, 138.96 (d, ^3^*J*C- F = 6.8 Hz), 131.49 (d, ^3^*J*C-F = 3.0 Hz), 130.80, 128.62, 128.31, 127.54 (d, ^2^*J*C-F = 14.9 Hz), 123.63 (d, ^4^*J*C-F = 3.4 Hz), 115.53 (d, ^2^*J*C-F = 25.6 Hz), 46.41, 41.32, 31.78; ^19^F NMR (471 MHz, DMSO) δ -110.88 (s); LC: Method 1, t_R_ = 1.85 min. MS (ESI): *m/z* = 453.25 [M - H]^-^; HRMS (ESI) for C_20_H_21_ClFN_2_O_5_S [M +H]^+^: *m/z* = calcd, 455.0844; found, 455.0844.

### (4-((4-(4-bromophenyl)piperidin-1-yl)sulfonyl)-2-fluorobenzoyl)glycine (62)

Compound **62** was prepared as described for **44**, using 4-(chlorosulfonyl)-2-fluorobenzoic acid (50 mg, 0.210 mmol) as a core. Off-white powder; yield 76% (66 mg, 0.131 mmol); ^1^H NMR (500 MHz, DMSO) δ 8.85 (td, *J* = 5.6, 1.4 Hz, 1H), 7.90 (t, *J* = 7.3 Hz, 1H), 7.74 – 7.68 (m, 2H), 7.46 (d, *J* = 8.4 Hz, 2H), 7.17 (d, *J* = 8.5 Hz, 2H), 3.96 (d, *J* = 5.9 Hz, 2H), 3.80 (d, *J* = 11.7 Hz, 2H), 2.43 – 2.37 (m, 2H), 1.81 (d, *J* = 11.3 Hz, 2H), 1.65 (qd, *J* = 12.7, 3.9 Hz, 2H); ^13^C NMR (126 MHz, DMSO) δ 170.69, 162.86, 158.88 (d, ^1^*J*C-F = 254.6 Hz), 144.49, 138.97 (d, ^3^*J* C-F = 6.9 Hz), 131.50 (d, ^3^*J* C-F = 2.9 Hz), 131.23, 129.03, 127.54 (d, ^2^*J* C-F = 14.9 Hz), 123.63 (d, ^4^*J* C-F = 3.5 Hz), 119.26, 115.53 (d, ^2^*J* C-F = 25.3 Hz), 46.40, 41.32, 39.78, 31.72; ^19^F NMR (471 MHz, DMSO) δ -110.88 (s); LC: Method 1, t_R_ = 1.91 min. MS (ESI): *m/z* = 497.27 [M - H]^-^, 499.30 [M - H]^-^ +2; HRMS (ESI) for C_20_H_21_BrFN_2_O_5_S [M + H]^+^: *m/z* = calcd, 499.0339; found, 499.0333.

### (2-fluoro-4-((4-(pyrrolidin-1-yl)piperidin-1-yl)sulfonyl)benzoyl)glycine (63)

Compound **63** was prepared as described for **44**, using 4-(chlorosulfonyl)-2-fluorobenzoic acid (50 mg, 0.210 mmol) as a core and 4-(1-pyrrolidinyl)piperidine (49 mg, 0.314 mmol) as amine. Off-white powder; yield 80% (66 mg, 0.143 mmol); obtained as a formic acid salt; ^1^H NMR (500 MHz, DMSO) δ 9.61 (s, 1H), 8.84 (dd, *J* = 5.8, 4.1 Hz, 1H), 7.93 – 7.88 (m, 1H), 7.75 (dd, *J* = 9.5, 1.5 Hz, 1H), 7.70 (dd, *J* = 8.0, 1.6 Hz, 1H), 3.96 (d, *J* = 5.9 Hz, 2H), 3.78 (d, *J* = 12.4 Hz, 2H), 3.49 (d, *J* = 4.9 Hz, 2H), 3.17 – 3.08 (m, 1H), 3.04 – 2.95 (m, 2H), 2.40 (t, *J* = 11.4 Hz, 2H), 2.10 (d, *J* = 11.4 Hz, 2H), 2.02 – 1.92 (m, 2H), 1.86 – 1.76 (m, 2H), 1.59 (qd, *J* = 12.3, 4.1 Hz, 2H); ^13^C NMR (126 MHz, DMSO) δ 170.67, 162.78, 158.92 (d, ^1^*J*C-F = 255.0 Hz), 138.97 (d, ^3^*J*C-F = 6.9 Hz), 131.61 (d, ^3^*J*C-F = 3.0 Hz), 127.65 (d, ^2^*J*C-F = 15.0 Hz), 123.59 (d, ^4^*J*C-F = 3.8 Hz), 115.55 (d, ^2^*J*C-F = 25.2 Hz), 59.66, 51.00, 44.36, 41.30, 27.41, 22.48; ^19^F NMR (471 MHz, DMSO) δ -110.70 (s); LC: Method 1, t_R_ = 0.99 min. MS (ESI): *m/z* = 414.35 [M - H]^-^; HRMS (ESI) for C_18_H_25_FN_3_O_5_S [M + H]^+^: *m/z* = calcd, 414.1499; found, 414.1501.

### (5-((4-(4-chlorophenyl)piperidin-1-yl)sulfonyl)picolinoyl)glycine (64)

Compound **64** was prepared as described for **51**, using methyl 5-(chlorosulfonyl)pyridine-2-carboxylate (50 mg, 0.212 mmol) as a core and 4-(4-chlorophenyl)piperidine hydrochloride (74 mg, 0.318 mmol) as amine. Off-white powder; yield 97% (30 mg, 0.066 mmol); ^1^H NMR (500 MHz, DMSO) δ 12.73 (s, 1H), 9.19 (t, *J* = 6.1 Hz, 1H), 9.00 (dd, *J* = 2.2, 0.7 Hz, 1H), 8.39 (dd, *J* = 8.2, 2.3 Hz, 1H), 8.29 (dd, *J* = 8.2, 0.7 Hz, 1H), 7.33 (d, *J* = 8.5 Hz, 2H), 7.22 (d, *J* = 8.5 Hz, 2H), 4.01 (d, *J* = 6.1 Hz, 2H), 3.83 (d, *J* = 11.6 Hz, 2H), 2.57 (tt, *J* = 12.1, 3.4 Hz, 1H), 2.44 (td, *J* = 11.8, 2.0 Hz, 2H), 1.82 (d, *J* = 11.2 Hz, 2H), 1.66 (qd, *J* = 12.7, 4.0 Hz, 2H); ^13^C NMR (126 MHz, DMSO) δ 170.79, 162.88, 152.59, 146.81, 144.05, 137.53, 134.29, 130.81, 128.60, 128.32, 122.70, 46.21, 41.17, 39.58, 31.77; LC: Method 1, t_R_ = 1.83 min. MS (ESI): *m/z* = 436.09 [M - H]^-^; HRMS (ESI) for C_19_H_21_ClN_3_O_5_S [M + H]^+^: *m/z* = calcd, 438.0890; found, 438.0882.

## ASSOCIATED CONTENT

### Supporting Information

- USP5-**1** coordinates (CIF)
- USP5-**1** sf (CIF)
- USP5-**48** coordinates (CIF)
- USP5-**48** sf (CIF)
- USP5-**64** coordinates (CIF)
- USP5-**64** sf (CIF)
- Supporting tables and figures, X-ray crystallographic statistics, methods, characterization data, ^1^H NMR, ^13^C NMR, ^19^F NMR, HLPC purity analysis for the compounds (PDF)
- Molecular formula strings (CSV)
- 3D PDB file used for docking studies (PDB)
  - Hit Expansion 1 6DXH
  - Hit Expansion 2 7MS5

## Accession Codes

7MS5, 7MS6, 7MS7

Authors will release atomic coordinates and experimental data upon article publication.

## AUTHOR INFORMATION

### Author Contributions

The manuscript was written by M.K.M, and M.S. and revised by R. J. H. All authors have given approval to the final version of the manuscript. M.K.M. completed hit expansion and docking, expressed and purified proteins, designed and optimized biophysical assays and tested compounds, and performed co-crystallization screens. A.D. solved the X-ray structure with compound **1**. R. J. H. solved the X-ray crystal structures with **48** and **64**. Analogs were synthesized and characterized by C.A.Z. and H.G.A. T.K. and A.A. performed liver microsomal stability studies and generated compound HRMS data. R.A., C.H.A, R.J.H. and M.S. advised throughout the project.

### Notes

The authors declare no competing financial interest.

## ACKNOWLEDGMENT

The Structural Genomics Consortium is a registered charity (no: 1097737) that receives funds from AbbVie, Bayer AG, Boehringer Ingelheim, Genentech, Genome Canada through Ontario Genomics Institute [OGI-196], the EU and EFPIA through the Innovative Medicines Initiative 2 Joint Undertaking [EUbOPEN grant 875510], Janssen, Merck KGaA (aka EMD in Canada and US), Pfizer, Takeda and the Wellcome Trust [106169/ZZ14/Z].We thank Albina Bolotokova for compound management, Darcy Burns and Jack Sheng for ^13^C NMR data collection, and Amir Jafari and Rick Briltz for their help in the procurement of the starting materials.

## ABBREVIATIONS

BRAP: BRCA-1 associated protein
DCM: dichloromethane
DMF: dimethylformamide
DMSO: dimethylsulfoxide
DIPEA: *N,N*-diisopropylethylamine
DUB: deubiquitinase
FL: full-length
HATU: hexafluorophosphate azabenzotriazole tetramethyl uranium
HDAC6: histone deacetylase 6
HRMS: high resolution mass spectrometry
IQF: internally quenched fluorophore
K_D_: dissociation constant
LCMS: liquid chromatography mass spectrometry
LRMS: low resolution mass spectrometry
QTOF: quadrupole-time-of-flight
SAR: structure activity relationship
SD: standard deviation
SPE: solid phase extraction
SPR: surface plasmon resonance
TFA: trifluoroacetic acid
Ub: ubiquitin
Ub2K48: Lys48-linked di-ubiquitin
UBA: ubiquitin-binding associated domain
Ub-AMC: ubiquitin amidomethyl coumarin
nUBP: N-terminal ubiquitin-binding domain
USP: ubiquitin specific protease
UPLC: ultra-performance liquid chromatography
WT: wild-type
ZnF-UBD: zinc finger ubiquitin-binding domain

