## Supporting Information for "Structure Activity Relationship of USP5 Allosteric Inhibitors"

*Corresponding Author

**Contents**

X-ray crystallographic statistics, supporting tables, figures & methods, S3-S13
Characterization data, $^{1}H$ NMR, $^{13}C$ NMR, $^{19}F$ NMR, LCMS: S14-S66

**Table S1**. Data collection and refinement statistics for USP5 co-crystal structures

| **PDB code** | **7MS5** | **7MS6** | **7MS7** |
|---|---|---|---|
| Compound | **1** | **48** | **64** |
| Space group | C 121 | P 43 | C 121 |
| a,b,c [Å] | 61.70, 84.49, 59.64 | 53.58, 53.58, 54.10 | 61.56, 84.87, 59.54 |
| α,β,γ [°] | 90.00, 98.27, 90.00 | 90.00, 90.00, 90.00 | 90.00, 98.86, 90.00 |
| Resolution limits [Å] | 29.53-1.98 | 38.10-1.55 | 40.42-1.45 |
| Rmerge | 0.10 | 0.04 | 0.05 |
| I/sigma | 2.38 | 12.90 | 2.54 |
| Completeness [%] | 95.3 | 99.3 | 99.4 |
| Multiplicity | 4.1 | 7.8 | 5.7 |
| No. Reflections used/free | 21165/1018 | 22304/1075 | 53141/2653 |
| Rwork/Rfree | 18.2/21.6 | 14.8/16.4 | 16.9/18.1 |
| No. atoms | | | |
| Protein | 1796 | 898 | 1918 |
| Inhibitor | 60 | 29 | 29 |
| Water | 215 | 137 | 261 |
| Others | 7 | 10 | 10 |
| rmsd bonds [Å]/ angles [°] | 0.0100/1.405 | 0.0153/1.978 | 0.0158/1.959 |
| B-factor [$Å^2$] | 22.0 | 14.0 | 22.0 |
| Molprobity Ramachandran favored/outliers [%] | 98/0 | 99/0 | 98/0 |

**Table S2**: Hit Expansion #1 Displacement Assay and SPR data

| Compound | Compound Structure | %Displacement USP5[a] | USP5 $K_D$ (μM)[b] | HDAC6 $K_D$ (μM)[b] |
|---|---|---|---|---|
| **1** | 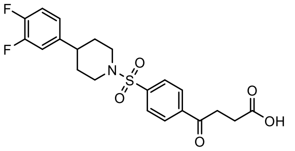 | 73 | 12 ± 3 | 12 ± 4 |
| **S1** | 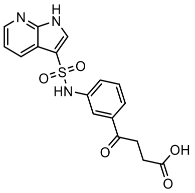 | 40 | 98 ± 25 | 190 ± 67 |
| **S2** | 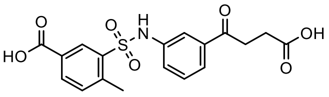 | 36 | 300 ± 55 | 300 ± 73 |
| **S3** | 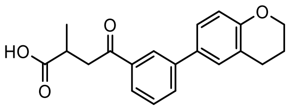 | 31 | 580 ± 160 | 150 ± 46 |
| **S4** | 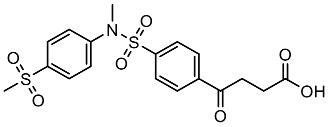 | 61 | 37 ± 5 | 13 ± 2 |
| **S5** | 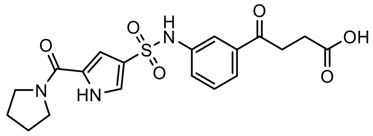 | 50 | 84 ± 12 | 210 ± 45 |
| **S6** | 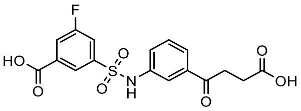 | 3 | N/A | |
| **S7** | 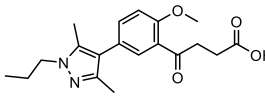 | 32 | 190 ± 16 | >770 |
| **S8** | 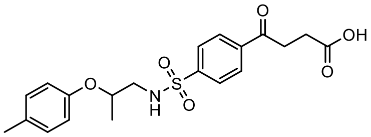 | 37 | 144 ± 24 | 64 ± 6 |
| **S9** | 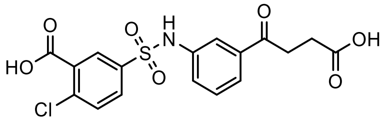 | 26 | 340 ± 150 | 300 ± 65 |

[a]Compounds tested at 100 μM (N=1); fluorescence normalized to control (DMSO). [b]$K_D$ determination experiments were performed with N ≥ 3 and the values are presented as mean ± SD. N/A= not tested.

**Table S3**: Hit expansion #2 SPR Data

| Compound | Compound Structure | USP5 $K_D$ (μM)[a] | HDAC6 $K_D$ (μM)[a] | Compound | Compound Structure | USP5 $K_D$ (μM)[a] | HDAC6 $K_D$ (μM)[a] |
|---|---|---|---|---|---|---|---|
| **2** | 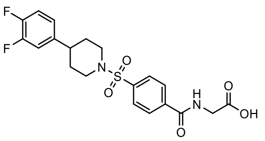 | 31 ± 6 | 270 ± 64 | **S15** | 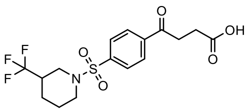 | 115 ± 18 | 44 ± 4 |
| **3** | 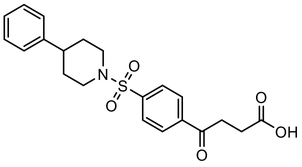 | 8 ± 2 | 7 ± 1 | **S16** | 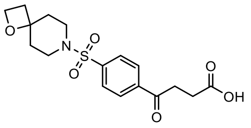 | 41 ± 9 | 6 ± 1 |
| **5** | 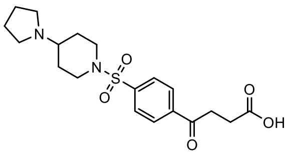 | 9 ± 3 | 15 ± 2 | **S17** | 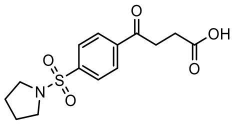 | 60 ± 8 | 22 ± 1 |
| **S10** | 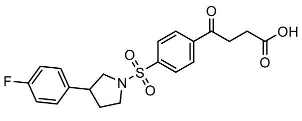 | 28 ± 2 | 25 ± 7 | **S18** | 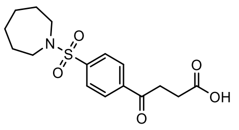 | 61 ± 8 | 26 ± 8 |
| **S11** | 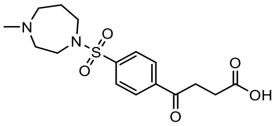 | 34 ± 4 | 33 ± 5 | **S19** | 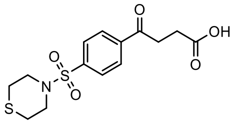 | 77 ± 10 | 31 ± 7 |
| **S12** | 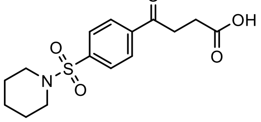 | 57 ± 7 | 28 ± 7 | **S20** | 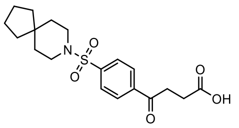 | 50 ± 6 | 24 ± 6 |
| **S13** | 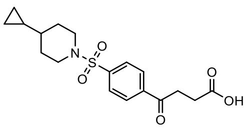 | 31 ± 5 | 25 ± 3 | **S21** | 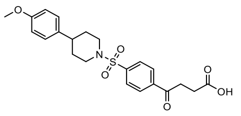 | 21 ± 5 | 19 ± 2 |
| **S14** | 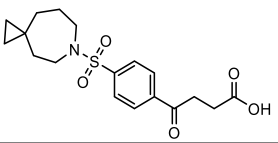 | 53 ± 5 | 27 ± 7 | **S22** | 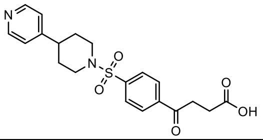 | 47 ± 8 | 51 ± 7 |

[a]$K_D$ determination experiments were performed with N ≥ 3 and the values are presented as mean ± SD.

### Materials & Methods: Chemistry

**Scheme S1. Synthesis of Compounds S24-S32*[a]***

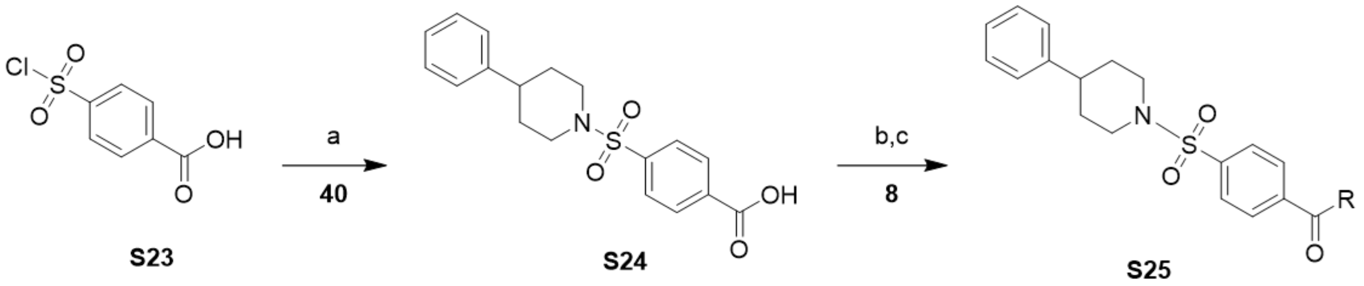

[a]Reagents and conditions: (a) 4-phenylpiperidine (**40**), DIPEA, DCM, 30 min, rt; (b) Various amines (**8**), HATU, DIPEA, DMF, 16 h, rt; (c) Only for **S28-S32**, TFA, DCM, 1 h, rt.

**4-((4-phenylpiperidin-1-yl)sulfonyl)benzoic acid (S24).** A solution of 4-(chlorosulfonyl)benzoic acid (2 g, 9.20 mmol), 4-phenyl-piperidine (2.2 g, 13.80 mmol) in DCM (92 mL) was stirred for 10 min at rt. The reaction mixture was diluted with 10 mL of DCM and two acidic washes (10 mL, 10% HCl each) were used to remove the excess of 4-phenyl-piperidine. The organic phase was collected, and the solvent was removed under vacuum. The resulting product was suspended in water/ACN and freeze-dried for 2 days to afford a white powder; yield 77% (2.5 g, 7.10 mmol). $^1$H NMR (500 MHz, DMSO) δ 13.52 (s, 1H), 8.18 (d, J = 8.3 Hz, 2H), 7.89 (d, J = 8.3 Hz, 2H), 7.30 – 7.25 (m, 2H), 7.20 – 7.16 (m, 3H), 3.79 (d, J = 11.6 Hz, 2H), 2.36 (t, J = 11.1 Hz, 2H), 1.81 (d, J = 11.9 Hz, 2H), 1.66 (qd, J = 12.6, 3.8 Hz, 2H); LC: Method 1, $t_R$ = 1.85 min, purity ($UV^{254}$) = 100%. MS (ESI): *m/z* = 344.22 $[M - 1]^-$; HRMS (ESI) for $C_{18}H_{20}NO_4S$ $[M + H]^+$: *m/z* = calcd, 346.1113; found, 346.1115.

**Methyl(4-((4-phenylpiperidin-1-yl)sulfonyl)benzoyl)glycinate (S25).** To a solution of 4-((4-phenylpiperidin-1-yl)sulfonyl)benzoic acid (50 mg, 0.145 mmol), N,N-Diisopropylethylamine (0.1 mL, 0.579 mmol) and HATU (83 mg, 0.217 mmol) in N,N-Dimethylformamide (DMF) (2 mL) was added methyl-2-aminoacetate HCl (27 mg, 0.217 mmol). The reaction mixture was stirred for 24 hours at rt. under nitrogen atmosphere. The solvents were evaporated under reduced pressure. DCM (10 mL) and aqueous HCl 10% (10 mL) were added and an insoluble precipitate was formed between the two layers. The organic phase was collected and washed two times with 10 mL of aqueous HCl 10%. The solvent was removed. Purification was performed via reverse-phase chromatography (95-5% to 0-100%, water (0.1% formic acid); ACN (0.1% formic acid)). The desired fractions were collected and evaporated under vacuum. Later, the resulting product was suspended in water/ACN and freeze-dried for 2 days to afford the final product as a white powder; yield 37% (23 mg, 0.054 mmol). $^1$H NMR (500 MHz, DMSO) δ 9.24 (t, J = 5.7 Hz, 1H), 8.11 (d, J = 8.3 Hz, 2H), 7.90 (d, J = 8.3 Hz, 2H), 7.27 (t, J = 7.6 Hz, 2H), 7.18 (d, J = 8.0 Hz, 3H), 4.06 (d, J = 5.8 Hz, 2H), 3.80 (d, J = 11.6 Hz, 2H), 3.67 (s, 3H), 2.34 (d, J = 12.1 Hz, 2H), 1.81 (d, J = 11.7 Hz, 2H), 1.66 (qd, J = 12.8, 3.9 Hz, 2H); LC: Method 1, $t_R$ = 1.83 min, purity ($UV^{254}$) = 100%. MS (ESI): *m/z* = 417.58 $[M + H]^+$; HRMS (ESI) for $C_{21}H_{25}N_2O_5S$ $[M + H]^+$: *m/z* = calcd, 417.1484; found, 417.1481.

**N-((1H-tetrazol-5-yl)methyl)-4-((4-phenylpiperidin-1-yl)sulfonyl)benzamide (S26).** Compound **S26** was prepared as described for **S25**, using (1H-1,2,3,4-tetrazol-5-yl)methylamine (22 mg, 0.217 mmol) as amine. White powder; yield 68% (42 mg, 0.098 mmol). $^1$H NMR (500 MHz, DMSO) δ 9.52 (t, J = 5.6 Hz, 1H), 8.14 (d, J = 8.6 Hz, 2H), 7.91 (d, J = 8.5 Hz, 2H), 7.30 – 7.25 (m, 2H), 7.20 – 7.16 (m, 3H), 4.80 (d, J = 5.6 Hz, 2H), 3.81 (d, J = 11.7 Hz, 2H), 2.49 – 2.44 (m, 1H), 2.34 (td, J = 11.8, 2.0 Hz, 2H), 1.81

(d, J = 11.0 Hz, 2H), 1.67 (qd, J = 12.7, 4.0 Hz, 2H); LC: Method 1, $t_R$ = 1.66 min, purity ($UV^{254}$) = 100%. MS (ESI): *m/z* = 425.41 $[M - H]^-$; HRMS (ESI) for $C_{20}H_{23}N_6O_3S$ $[M + H]^+$: *m/z* = calcd, 427.1552; found, 427.1551.

**4-((4-phenylpiperidin-1-yl)sulfonyl)-N-(1H-tetrazol-5-yl)benzamide (S27).** Compound **S27** was prepared as described for **S25**, using 1H-tetrazol-5-amine (18 mg, 0.217 mmol) as amine. White powder; yield 38% (24 mg, 0.055 mmol). $^1$H NMR (500 MHz, DMSO) δ 8.21 (d, J = 8.4 Hz, 2H), 7.96 (d, J = 8.3 Hz, 2H), 7.27 (t, J = 7.5 Hz, 2H), 7.21 – 7.15 (m, 3H), 3.80 (d, J = 11.7 Hz, 2H), 2.38 (t, J = 11.1 Hz, 2H), 1.81 (d, J = 12.1 Hz, 2H), 1.66 (qd, J = 12.6, 3.8 Hz, 2H); LC: Method 1, $t_R$ = 2.18 min, purity ($UV^{254}$) = 95%. MS (ESI): *m/z* = cannot be observed $[M + H]^+$; HRMS (ESI) for $C_{19}H_{21}N_6O_3S$ $[M + H]^+$: *m/z* = calcd, 413.1396; found, N/A. HRMS of the major fragment (loss of tetrazole group) (ESI) for $C_{18}H_{19}N_2O_3S$ $[M + H]^+$: m/z = calcd, 343.1111; found, 343.1117.

**(4-((4-phenylpiperidin-1-yl)sulfonyl)benzoyl)-L-alanine (S28).** To a solution of 4-((4-phenylpiperidin-1-yl)sulfonyl)benzoic acid (50 mg, 0.145 mmol), N,N-Diisopropylethylamine (0.1 mL, 0.579 mmol) and HATU (83 mg, 0.217 mmol) in N,N-Dimethylformamide (DMF) (2 mL) was added L-alanine t-butyl ester hydrochloride (40 mg, 0.217 mmol). The reaction mixture was stirred for 24 hours at rt. under nitrogen atmosphere. The solvents were evaporated under vacuum. DCM (10 mL) and aqueous HCl 10% (10 mL) were added and an insoluble precipitate was formed between the two layers. The organic phase was collected and washed two times with 10 mL of aqueous HCl 10%. The solvent was evaporated. Afterwards, the resulting crude was redissolved in DCM (1 mL) and TFA (0.750 mL) was added. The reaction mixture was stirred for 1 h. The volatiles were evaporated. Purification was performed via reverse-phase chromatography (95-5% to 0-100%, water (0.1% formic acid); ACN (0.1% formic acid)). The desired fractions were collected and evaporated under vacuum. Later, the resulting product was suspended in water/ACN and freeze-dried for 2 days to afford the final product as a white powder; yield 81% (49mg, 0.118 mmol). $^1$H NMR (500 MHz, DMSO) δ 12.63 (s, 1H), 8.95 (d, J = 7.2 Hz, 1H), 8.13 (d, J = 8.6 Hz, 2H), 7.88 (d, J = 8.5 Hz, 2H), 7.30 – 7.24 (m, 2H), 7.18 (ddd, J = 7.1, 3.3, 2.4 Hz, 3H), 4.45 (p, J = 7.3 Hz, 1H), 3.80 (d, J = 11.6 Hz, 2H), 2.34 (dd, J = 12.0, 10.1 Hz, 2H), 1.81 (d, J = 11.0 Hz, 2H), 1.66 (qd, J = 12.6, 3.9 Hz, 2H), 1.42 (d, J = 7.4 Hz, 3H); LC: Method 1, $t_R$ = 1.72 min, purity ($UV^{254}$) = 100%. MS (ESI): *m/z* = 415.33 $[M - H]^-$; HRMS (ESI) for $C_{21}H_{25}N_2O_5S$ $[M + H]^+$: *m/z* = calcd, 417.1484; found, 417.1479.

**(4-((4-phenylpiperidin-1-yl)sulfonyl)benzoyl)-D-alanine (S29).** Compound **S29** was prepared as described for **S28**, using tert-butyl (2R)-2-aminopropanoate hydrochloride (39 mg, 0.217 mmol) as amine. White powder; yield 54% (34 mg, 0.078 mmol). $^1$H NMR (500 MHz, DMSO) δ 12.62 (s, 1H), 8.95 (d, J = 7.2 Hz, 1H), 8.13 (d, J = 8.5 Hz, 2H), 7.88 (d, J = 8.5 Hz, 2H), 7.29 – 7.25 (m, 2H), 7.20 – 7.15 (m, 3H), 4.44 (p, J = 7.3 Hz, 1H), 3.80 (d, J = 11.6 Hz, 2H), 2.37 – 2.29 (m, 2H), 1.81 (d, J = 11.2 Hz, 2H), 1.66 (qd, J = 12.7, 3.9 Hz, 2H), 1.41 (d, J = 7.4 Hz, 3H); LC: Method 1, $t_R$ = 1.72 min, purity ($UV^{254}$) = 95%. MS (ESI): *m/z* = 415.48 $[M - H]^-$; HRMS (ESI) for $C_{21}H_{25}N_2O_5S$ $[M + H]^+$: *m/z* = calcd, 417.1484; found, 417.1488.

**2-methyl-2-(4-((4-phenylpiperidin-1-yl)sulfonyl)benzamido)propanoic acid (S30).** Compound **S30** was prepared as described for **S28**, using tert-butyl 2-amino-2-methylpropanoate hydrochloride (57 mg, 0.290 mmol) as amine. White powder; yield 64% (40mg, 0.093 mmol). $^1$H NMR (500 MHz, DMSO) δ 12.27 (s, 1H), 8.74 (s, 1H), 8.09 (d, J = 8.6 Hz, 2H), 7.86 (d, J = 8.5 Hz, 2H), 7.29 – 7.24 (m, 2H), 7.20 – 7.16 (m, 3H), 3.79 (d, J = 11.6 Hz, 2H), 2.47 (tt, J = 12.2, 3.5 Hz, 1H), 2.32 (td, J = 11.9, 2.1 Hz, 2H), 1.81 (d, J = 11.0 Hz, 2H), 1.67 (qd, J = 12.7, 4.0 Hz, 2H), 1.48 (s, 6H); LC: Method 1, $t_R$ = 1.77 min, purity ($UV^{254}$) = 100%. MS (ESI): *m/z* = 429.40 $[M - H]^-$; HRMS (ESI) for $C_{22}H_{27}N_2O_5S$ $[M + H]^+$: *m/z* = calcd, 431.1641; found, 431.1639.

**N-methyl-N-(4-((4-phenylpiperidin-1-yl)sulfonyl)benzoyl)glycine (S31).** Compound **S31** was prepared as described for **S28**, using tert-butyl 2-(methylamino)acetate (32 mg, 0.217 mmol) as amine. White powder; yield 88% (53 mg, 0.127 mmol). $^{1}$H NMR (500 MHz, DMSO) δ 7.86 (d, J = 8.3 Hz, 2H), 7.68 (d, J = 8.3 Hz, 2H), 7.27 (t, J = 7.6 Hz, 2H), 7.20 – 7.16 (m, 3H), 4.18 (s, 2H), 3.80 (d, J = 8.9 Hz, 2H), 2.94 (s, 3H), 2.37 (q, J = 12.3 Hz, 2H), 1.81 (d, J = 11.1 Hz, 2H), 1.71 – 1.61 (m, 2H); LC: Method 1, $t_R$ = 1.69 min, purity ($UV^{254}$) = 100%. MS (ESI): *m/z* = 415.40 $[M - H]^-$; HRMS (ESI) for $C_{21}H_{25}N_2O_5S$ $[M + H]^+$: *m/z* = calcd, 417.1484; found, 417.1490.

**3-(4-((4-phenylpiperidin-1-yl)sulfonyl)benzamido)propanoic acid (S32).** Compound **S32** was prepared as described for **S28**, using beta-alanine t-butyl ester hydrochloride (39 mg, 0.217 mmol) as amine. White powder; yield 32% (20 mg, 0.047 mmol). $^{1}$H NMR (500 MHz, DMSO) δ 8.81 (t, J = 5.3 Hz, 1H), 8.07 (d, J = 8.3 Hz, 2H), 7.86 (d, J = 8.3 Hz, 2H), 7.27 (t, J = 7.5 Hz, 2H), 7.21 – 7.15 (m, 3H), 3.79 (d, J = 11.6 Hz, 2H), 3.49 (dt, J = 12.6, 6.7 Hz, 2H), 2.54 (t, J = 7.0 Hz, 2H), 2.33 (t, J = 12.1 Hz, 2H), 1.81 (d, J = 11.9 Hz, 2H), 1.66 (qd, J = 12.7, 3.8 Hz, 2H); LC: Method 1, $t_R$ = 1.68 min, purity ($UV^{254}$) = 100%. MS (ESI): *m/z* = 415.48 $[M - 1]^-$; HRMS (ESI) for $C_{21}H_{25}N_2O_5S$ $[M + H]^+$: *m/z* = calcd, 417.1484; found, 417.1485.

**Table S4. Potency and selectivity of S24-32**

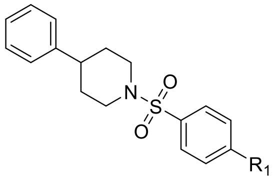


| Compound | $R_1$ | USP5 $K_D$ [μM][a] | HDAC6 $K_D$ [μM][a] | Fold Selectivity | Compound | $R_1$ | USP5 $K_D$ [μM][a] | HDAC6 $K_D$ [μM][a] | Fold Selectivity |
|---|---|---|---|---|---|---|---|---|---|
| **S25** | 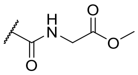 | NB[b] | NB[b] | - | **S30** | 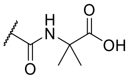 | NB[b] | NB[b] | - |
| **S26** | 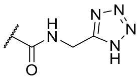 | NB[b] | 180 ± 23 | - | **S31** | 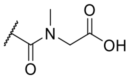 | NB[b] | NB[b] | - |
| **S27** | 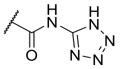 | NB[b] | NB[b] | - | **S32** | 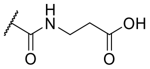 | NB[b] | NB[b] | - |
| **S28** | 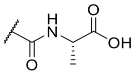 | NB[b] | NB[b] | - | **S24** | 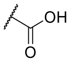 | NB[b] | NB[b] | - |
| **S29** | 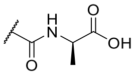 | 310 ± 121 | 70 ± 31 | - | | | | | |

[a]$K_D$ determination experiments were performed with N ≥ 3 and the values are presented as mean ± SD. [b]$K_D$ determination experiments were performed with N = 2. NB: no binding ($K_D$ > 1 mM).

### Materials and Methods: Liver microsomal metabolic stability

Stock solutions of test compounds in DMSO (1 mM) were initially diluted to a concentration of 40.0 µM using 0.1 M potassium phosphate buffer (pH 7.4). Test compounds were then added to reaction wells at a final concentration of 1 µM which was assumed to be well below Km values to ensure linear reaction conditions (i.e. avoid saturation). The final DMSO concentration was kept constant at 0.1%. Each compound was tested in duplicate for both time points (0 and 60 minutes). CD-1 mouse (male) or pooled human liver microsomes (Corning Gentest) were added to the reaction wells at a final concentration of 0.5 mg/mL (protein). The final volume for each reaction was 100 µL, which included the NADPH-Regeneration Solution (NRS) mix (Corning Gentest). This NRS mix was comprised of glucose 6-phosphate dehydrogenase, $NADP^+$, $MgCl_2$, and glucose 6-phosphate. Reactions were carried out at 37°C in an orbital shaker at 175 rpm. Upon completion of the 60-minute time point, reactions were terminated by the addition of 2-volumes (200 µL) of acetonitrile containing 0.5% formic acid and an internal standard. Samples were then centrifuged at 4,000 rpm for 10 minutes to remove debris and precipitated proteins. Approximately 150 µL of supernatant was subsequently transferred to a new 96 well microplate for LC/MS analysis.
Narrow-window mass extraction LC-MS analysis was performed for all samples in this study using a Waters Xevo quadrupole time-of-flight (QTof) mass spectrometer to determine relative peak areas of test compounds. The percent remaining values were calculated using the following equations:

$$\% \, remaining = \frac{A}{A_0} \times 100$$

Where
A is area response after incubation
$A_0$ is area response at initial time point

**Table S5**. Stability of **64** in Mouse and Human Liver Microsomes (MLM, HLM)

| Compound | Compound Structure | Phase | % Remaining Mouse | Classification Mouse | % Remaining Human | Classification Human |
|---|---|---|---|---|---|---|
| **64** | 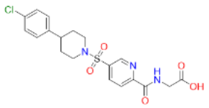 | I | 105.3 ± 5.0 | Stable | 92.6 ± 3.8 | Stable |
| Stable control in MLM & HLM-antipyrine | 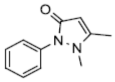 | I | 85.2 ± 4.9 | Stable | 95.2 ± 7.0 | Stable |
| Moderate stability control in HLM-dextromethorphan | 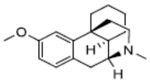 | I | 1.1 ± 0.2 | Unstable | 50.7 ± 3.6 | Moderate |
| Unstable control in MLM-propranolol | 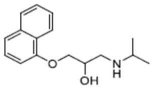 | I | 2.7 ± 0.3 | Unstable | N/A | N/A |
| Unstable control in HLM- verapamil | 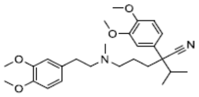 | I | N/A | N/A | 1.0 ± 0.0 | Unstable |

**Table S6:** HRMS Experimental Conditions

<table>
<tr><td>Mass analyzer</td><td>Waters Synapt G2-S Q-TOF mass spectrometer</td></tr>
<tr><td colspan="2">Sample probe</td></tr>
<tr><td>Ionization mode</td><td>ESI+</td></tr>
<tr><td>Capillary voltage</td><td>1.0 kV</td></tr>
<tr><td>Cone voltage</td><td>25 V</td></tr>
<tr><td>Source temperature</td><td>150 °C</td></tr>
<tr><td>Desolvation temperature</td><td>500 °C</td></tr>
<tr><td>Cone gas flow</td><td>150 L $h^{-1}$</td></tr>
<tr><td>desolvation gas flow</td><td>500 L $h^{-1}$</td></tr>
<tr><td>Scan time</td><td>0.3 s</td></tr>
<tr><td colspan="2">Reference probe</td></tr>
<tr><td>Capillary voltage</td><td>3.0 kV</td></tr>
<tr><td>Scan time</td><td>0.3 s</td></tr>
<tr><td>collision energy</td><td>19 V</td></tr>
<tr><td>Reference substance (ions)</td><td>Leucine-Enkephalin (m/z 221.0926 and 556.2771)</td></tr>
<tr><td>Liquid chromatography</td><td>Waters ACQUITY UPLC I-Class system</td></tr>
<tr><td>Column</td><td>Waters ACQUITY UPLC HSS T3 column (2.1 × 50 mm, 1.8 μm at 40 °C)</td></tr>
<tr><td>Mobile phase A</td><td>0.1% formic acid in water</td></tr>
<tr><td>Mobile phase B</td><td>0.1% formic acid in acetonitrile</td></tr>
<tr><td>Gradient</td><td><table>
<tr><td>Time/min</td><td>A%</td><td>Flow (μL/min)</td></tr>
<tr><td>0.0</td><td>95</td><td>400</td></tr>
<tr><td>2.0</td><td>5</td><td>400</td></tr>
<tr><td>2.5</td><td>5</td><td>400</td></tr>
<tr><td>3.0</td><td>95</td><td>400</td></tr>
<tr><td>4.0</td><td>95</td><td>400</td></tr>
</table></td></tr>
</table>

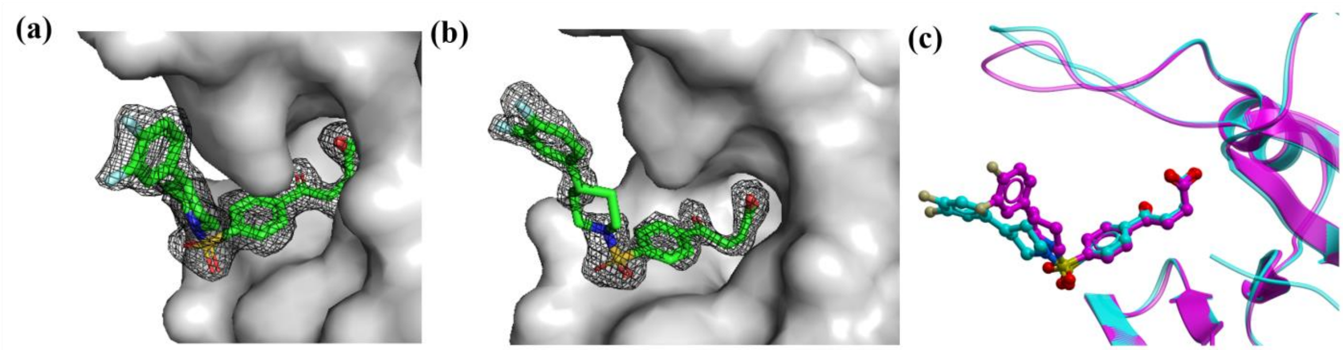


**Figure S1**. Ligand pose of **1 (PDB: 7MS5)**: (a) Omit map (σ2) of **1** in USP5 ZnF-UBD chain A (b) Omit map (σ2) of **1** in USP5 ZnF-UBD chain B (c) superimposed chain A (magenta) and B (cyan) (RMSD of ligand= 3.11)

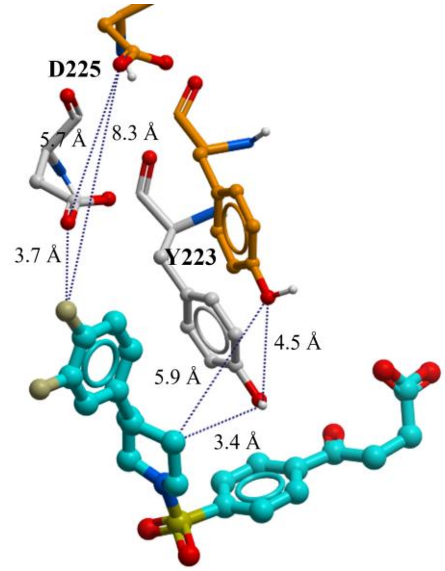


**Figure S2**. Co-crystal structure of USP5 ZnF-UBD in complex with **1 (PDB: 7MS5)** superimposed with the crystal structure of USP5 ZnF-UBD (PDB: 6DXH) (orange) highlighting the flexible loop residues Tyr223 and Asp225, and distances of residue movement. The binding pocket residues and ligands are displayed as ball and sticks.

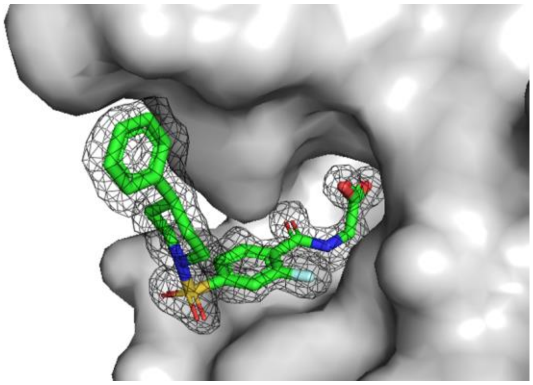

**Figure S3**. Omit map (σ2) of USP5 ZnF-UBD and compound **48 (PDB: 7MS6)**

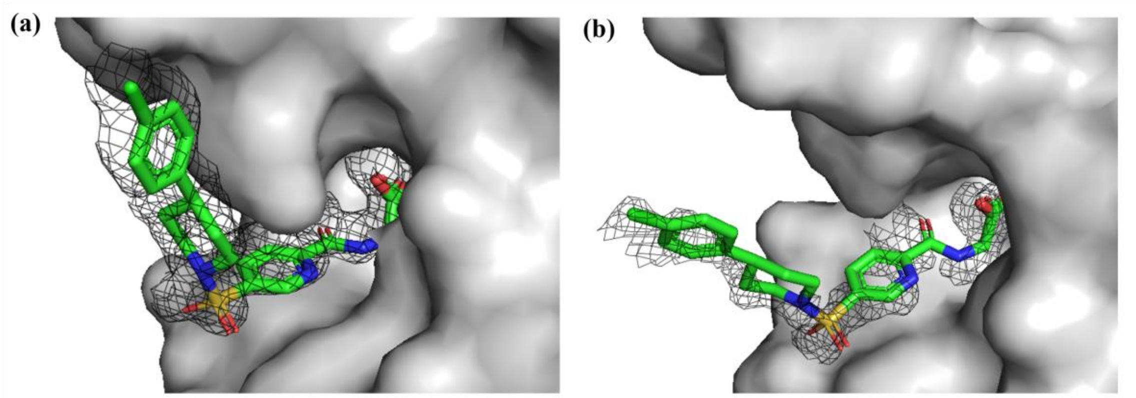


**Figure S4.** Ligand pose of **64 (PDB: 7MS7**): (a) Omit map (σ2) of **64** in USP5 ZnF-UBD chain A (b) Omit map (σ2) of **64** in USP5 ZnF-UBD chain B

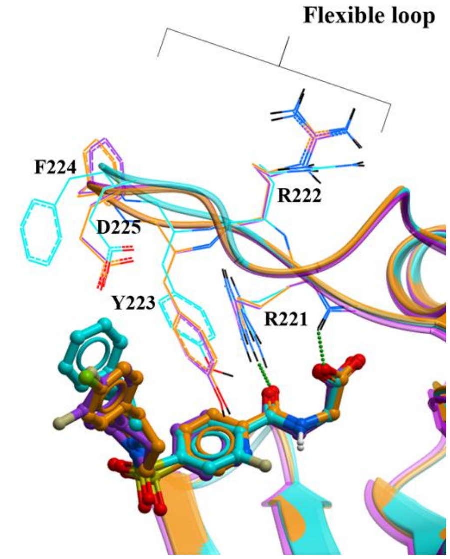


**Figure S5. Flexible loop residues of USP5 ZnF-UBD:** Superimposed structures of USP5 ZnF-UBD in complex with **1** (purple) (**PDB: 7MS5)**, **48** (cyan) (**PDB: 7MS6)** and **64** (orange) (**PDB: 7MS7)** showing flexibility of residues 222-225.

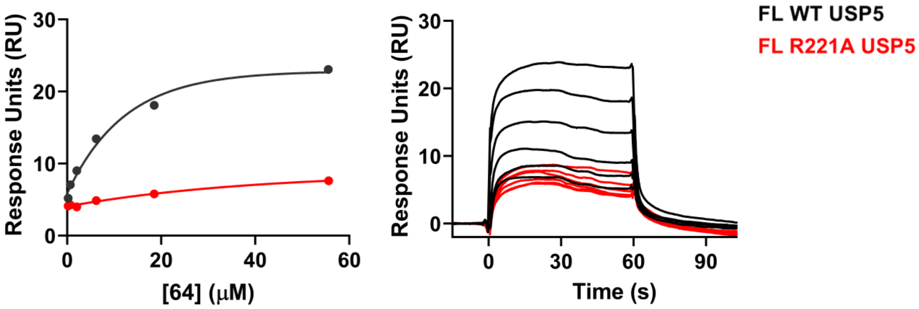


**Figure S6**. **Binding of 64 to FL USP5**: Representative SPR binding curve and sensorgram for FL WT USP5 versus FL R221A USP5. A $K_D$ of 8 ± 2 μM was obtained from the average of three independent measurements for FL WT USP5. No binding was observed from three independent measurements for FL R221A USP5.

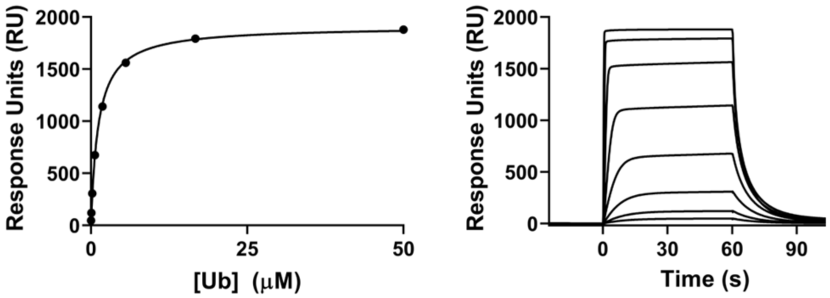


**Figure S7**. **Ub binding to USP5 ZnF-UBD**: Representative SPR binding curve and sensorgram for USP5 ZnF-UBD and Ub. A $K_D$ of 1.4 ± 0.2 μM was obtained from the average of six independent measurements.

**[1]**

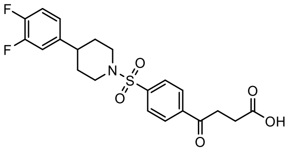

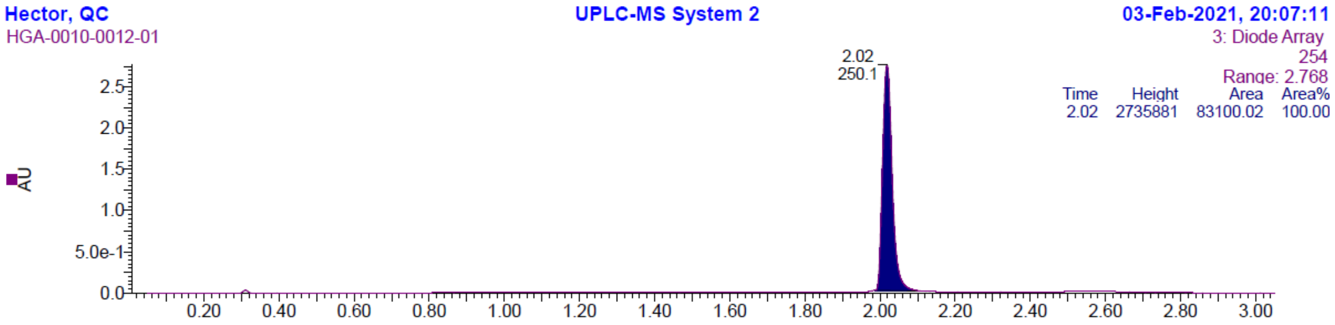


LC: Method 1, $t_R$= 2.02 min. MS (ESI): *m/z*= 438.19 $[M + H]^+$

**[2]**

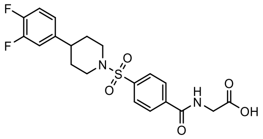

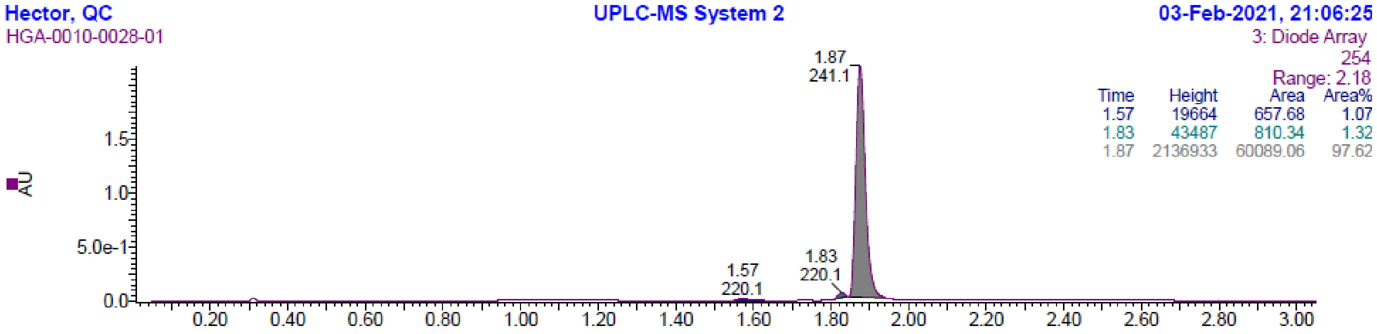


LC: Method 1, $t_R$ = 1.87 min. MS (ESI): m/z = 437.25 $[M - H]^-$

**[3]**

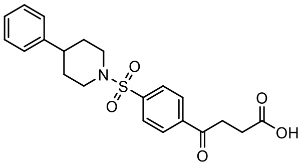

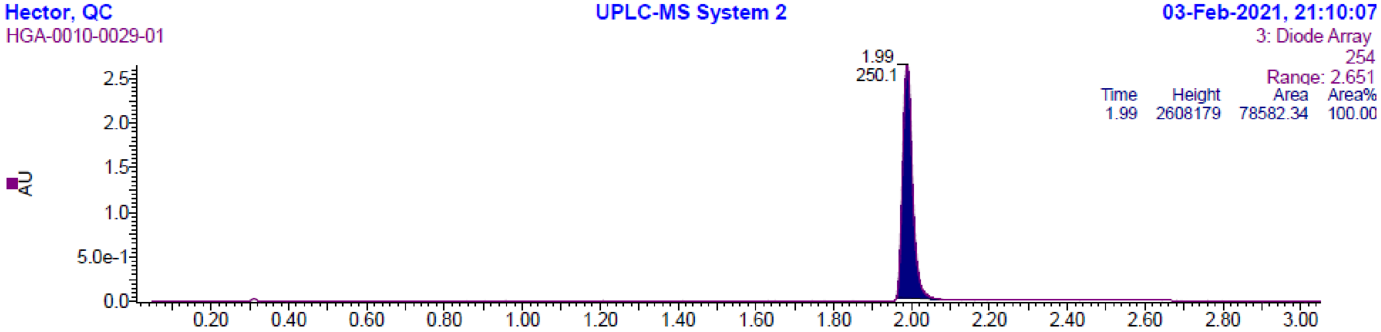


LC: Method 1, $t_R$= 1.99 min. MS (ESI): *m/z*= 402.27 $[M + H]^+$

## [4]

$^{1}$H NMR (500 MHz, DMSO-*d6*)

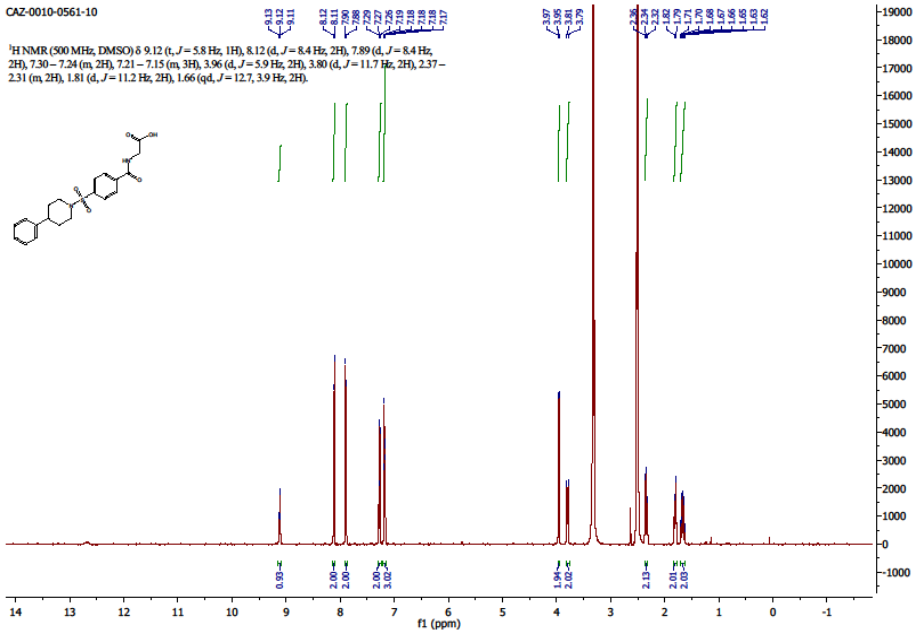


## [5]

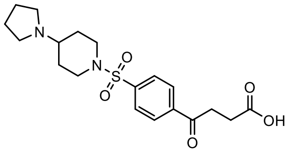

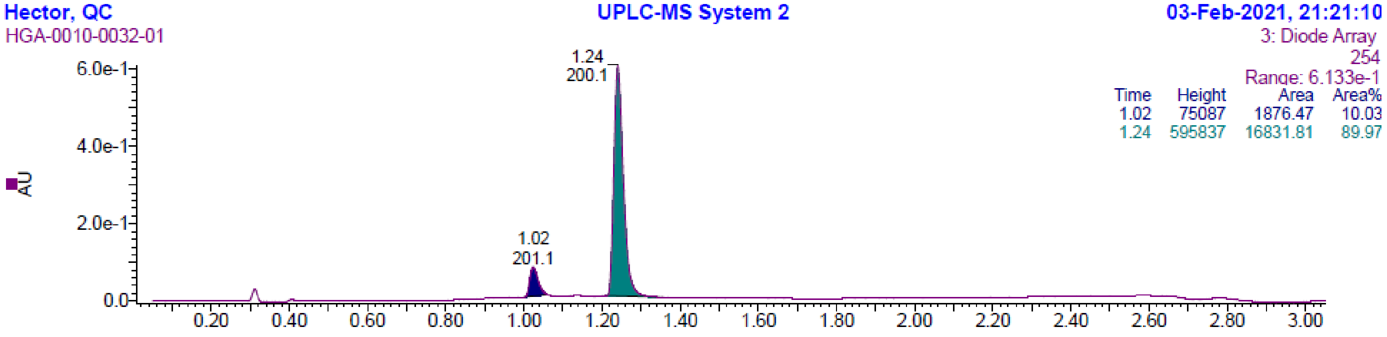


LC: Method 1, $t_R$ = 1.24 min. MS (ESI): m/z = 393.37 [M - H]$^-$

## [6]

$^{1}$H NMR (500 MHz, DMSO-*d6*)

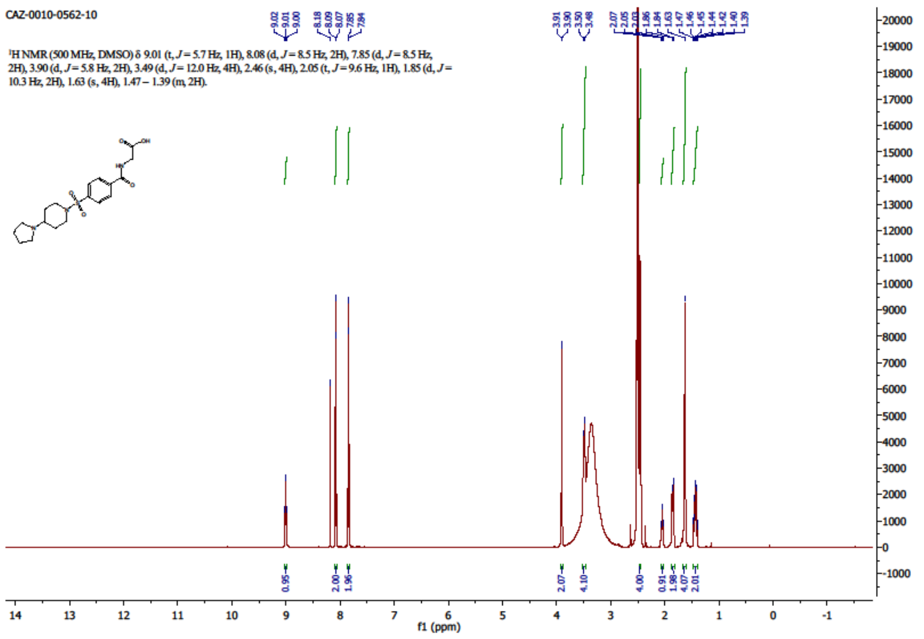


## [9]

$^{1}$H NMR (500 MHz, DMSO-*d6*)

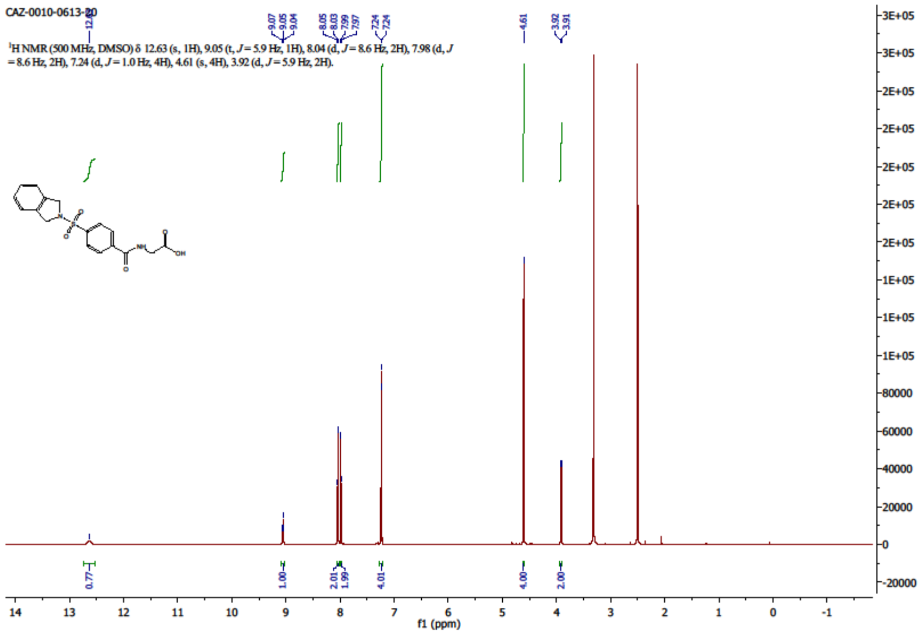

**[10]**

[1]H NMR (500 MHz, DMSO-*d6*)

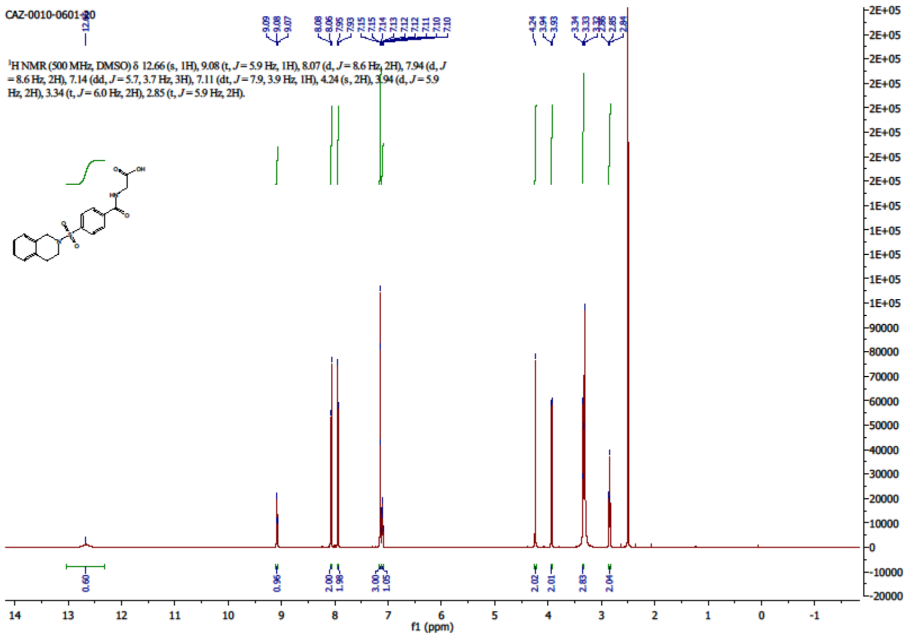


**[11]**

[1]H NMR (500 MHz, DMSO-*d6*)

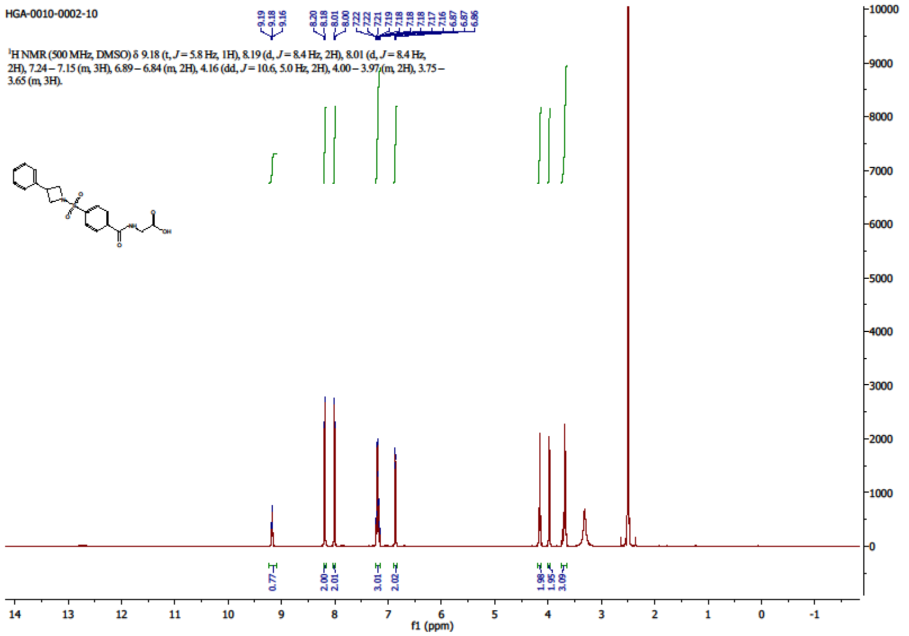

**[12]**

$^{1}$H NMR (500 MHz, DMSO-*d6*)

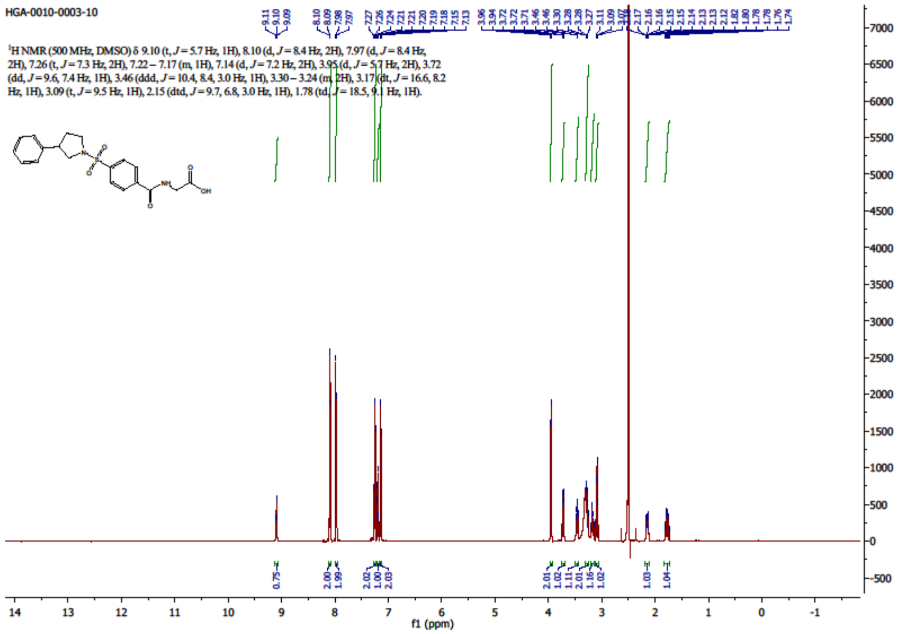


**[13]**

$^{1}$H NMR (500 MHz, DMSO-*d6*)

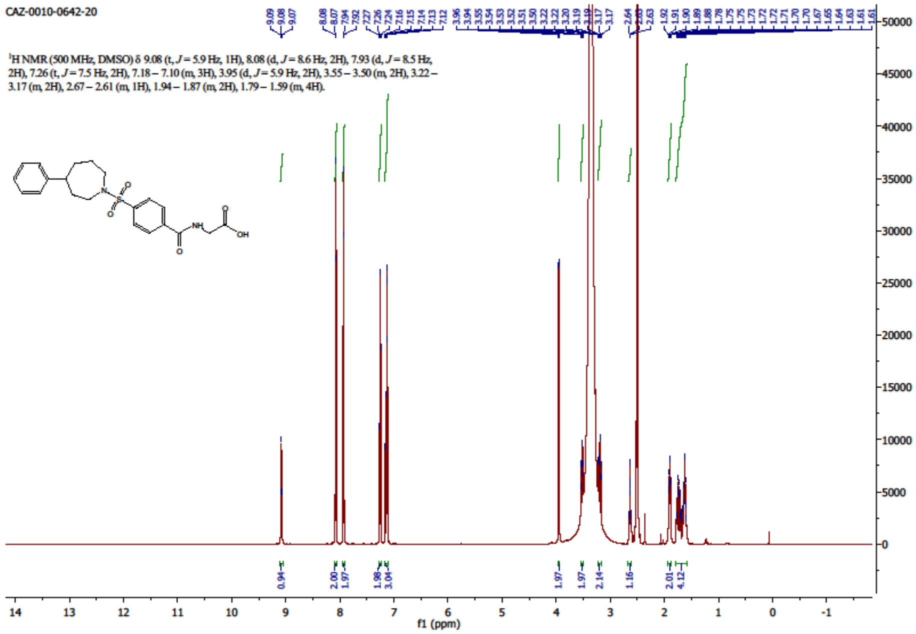

**[14]**

[1]H NMR (500 MHz, DMSO-*d6*)

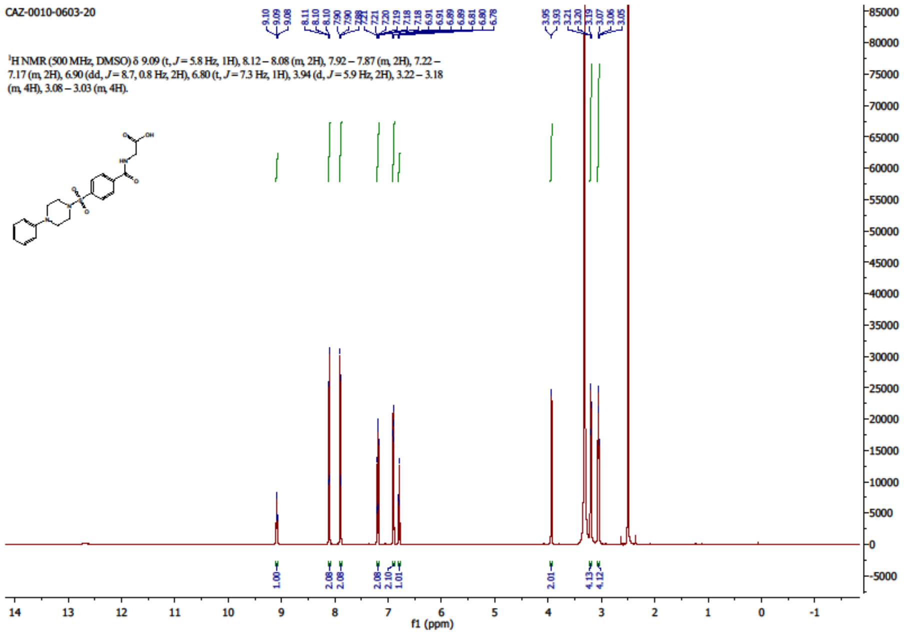

**[15]**

[1]H NMR (500 MHz, DMSO-*d6*)

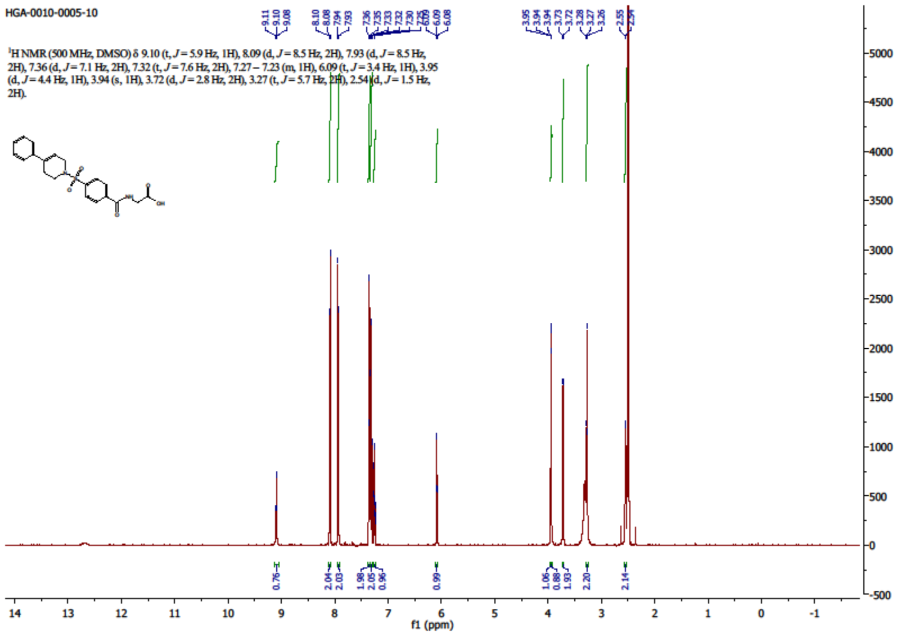

**[16]**

$^{1}$H NMR (500 MHz, DMSO-*d6*)

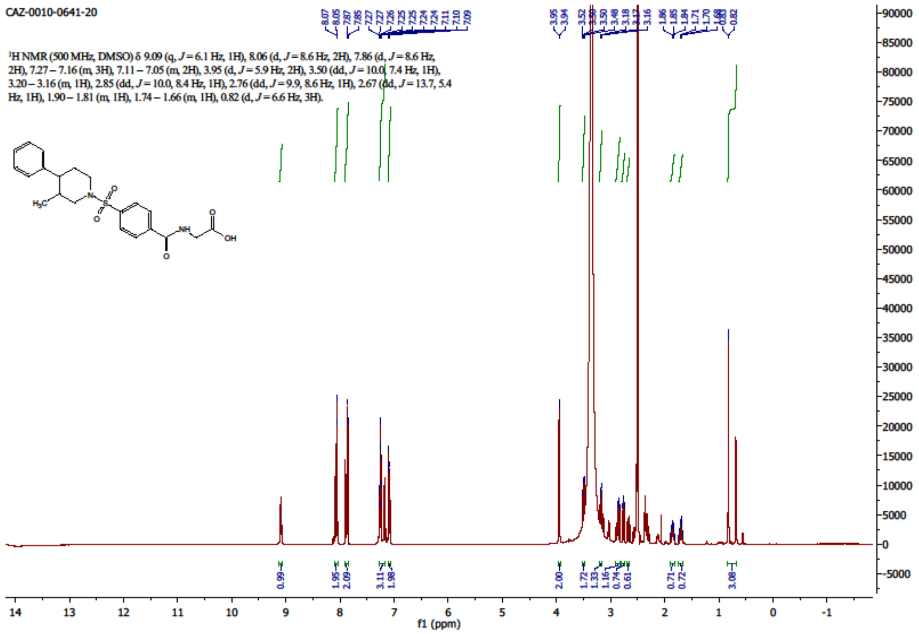


**[17]**

$^{1}$H NMR (500 MHz, DMSO-*d6*)

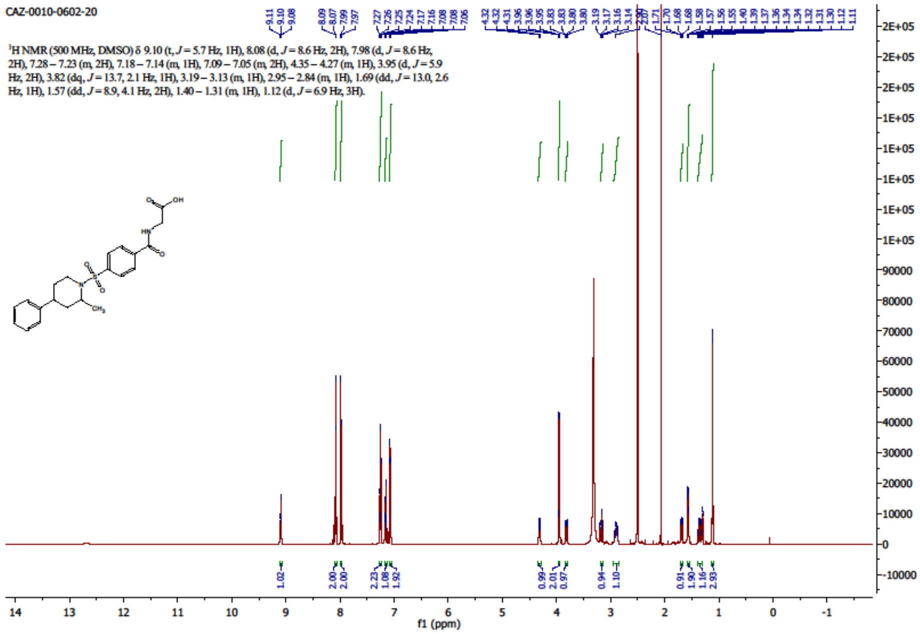

**[18]**

[1]H NMR (500 MHz, DMSO-*d6*)

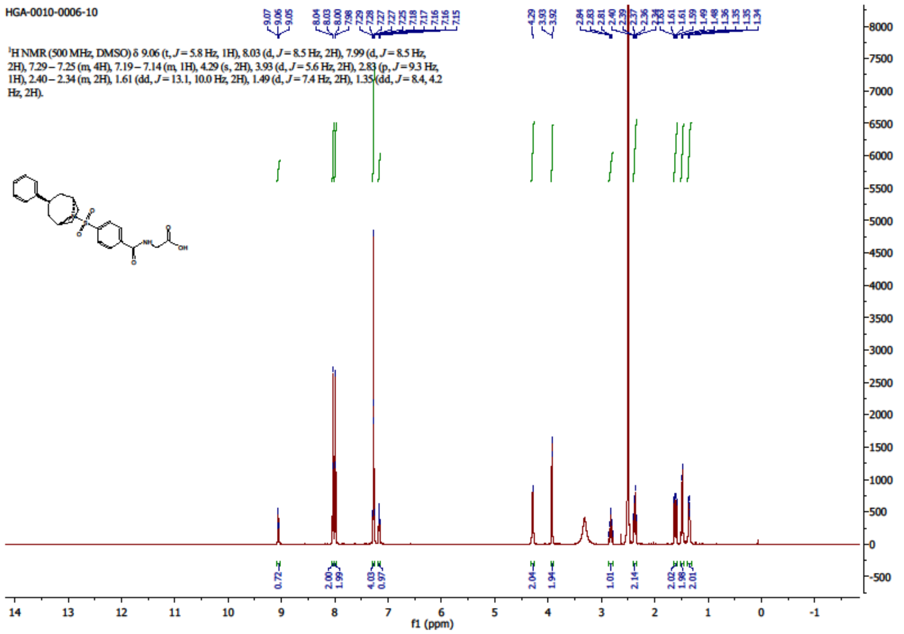


**[19]**

[1]H NMR (500 MHz, DMSO-*d6*)

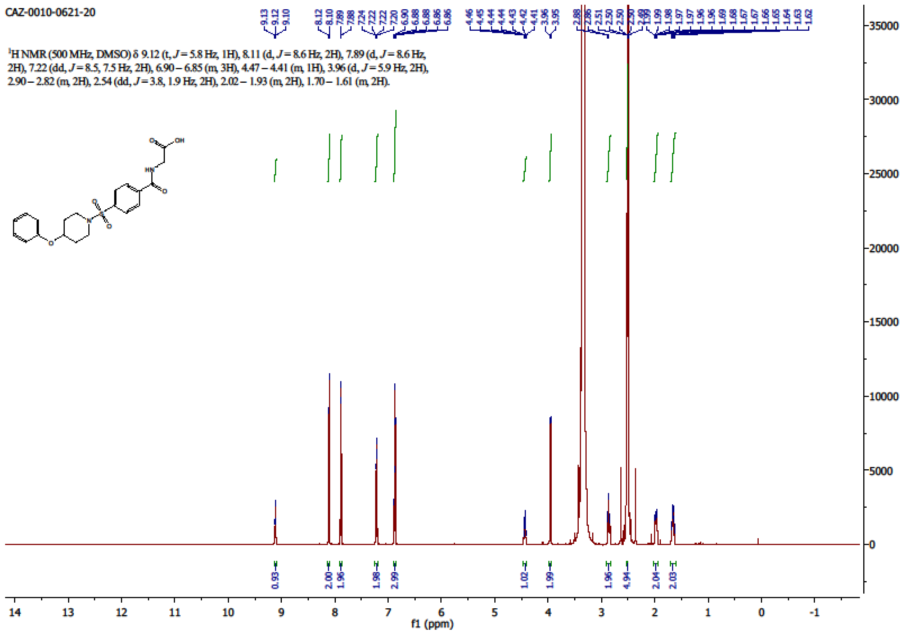

**[20]**

$^{1}$H NMR (500 MHz, DMSO-*d6*)

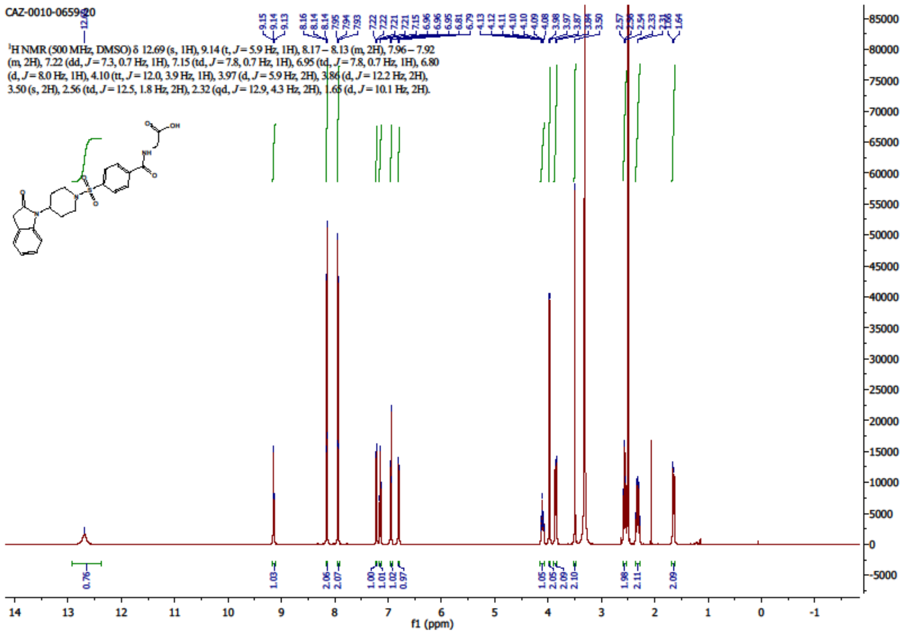


**[21]**

$^{1}$H NMR (500 MHz, DMSO-*d6*)

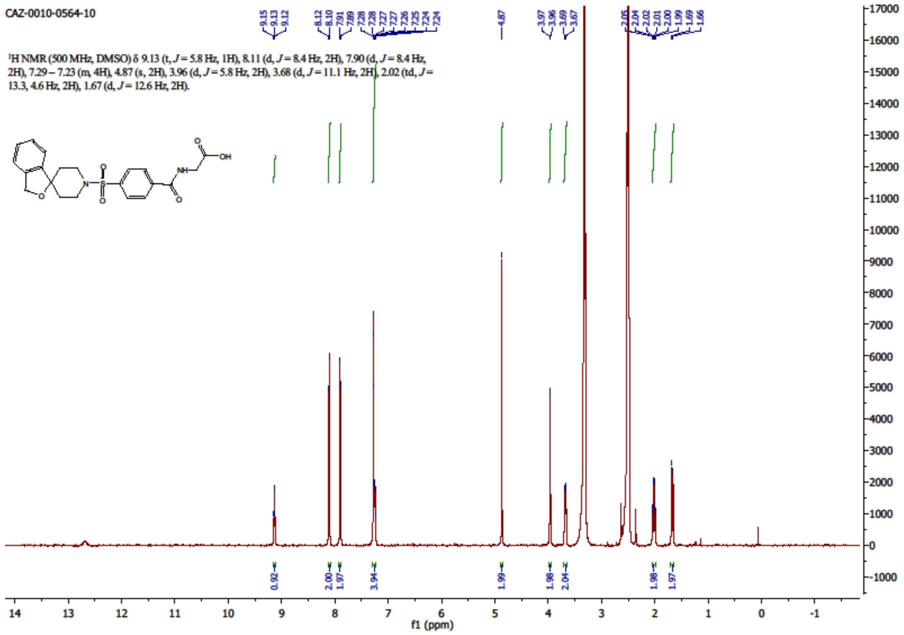

**[22]**

[1]H NMR (500 MHz, DMSO-*d6*) of compound **22**

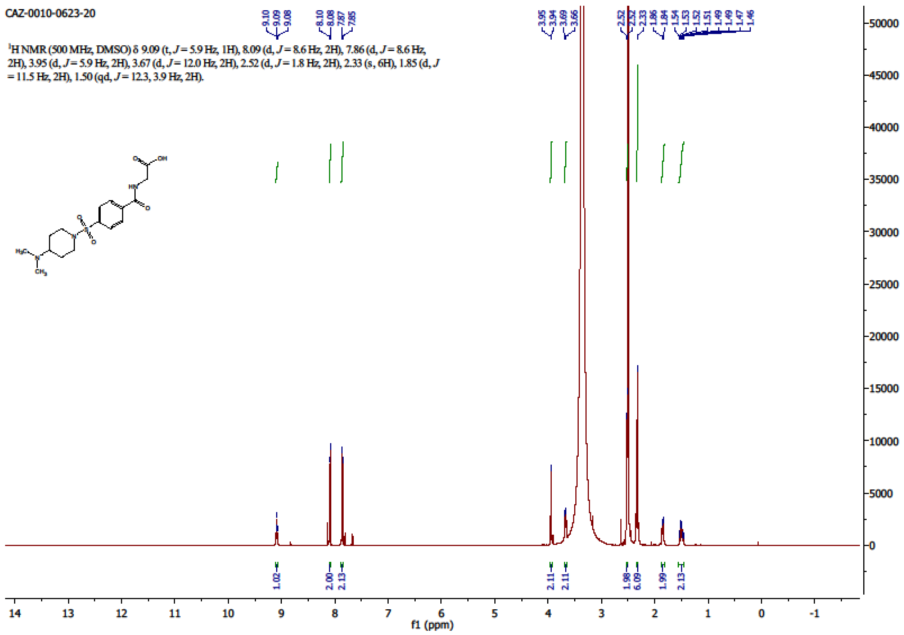


**[23]**

[1]H NMR (500 MHz, DMSO-*d6*)

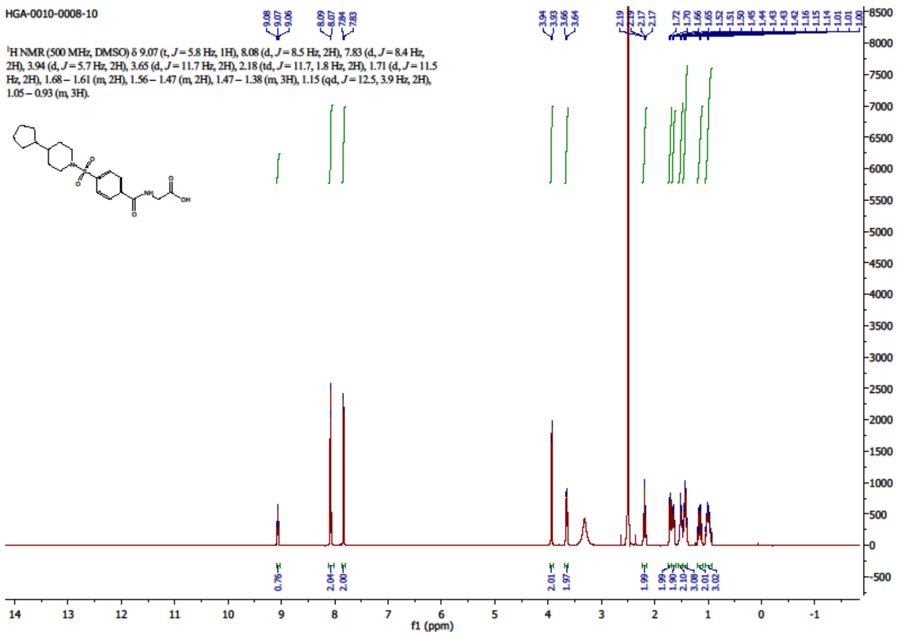

**[24]**

$^1$H NMR (500 MHz, DMSO-*d6*)

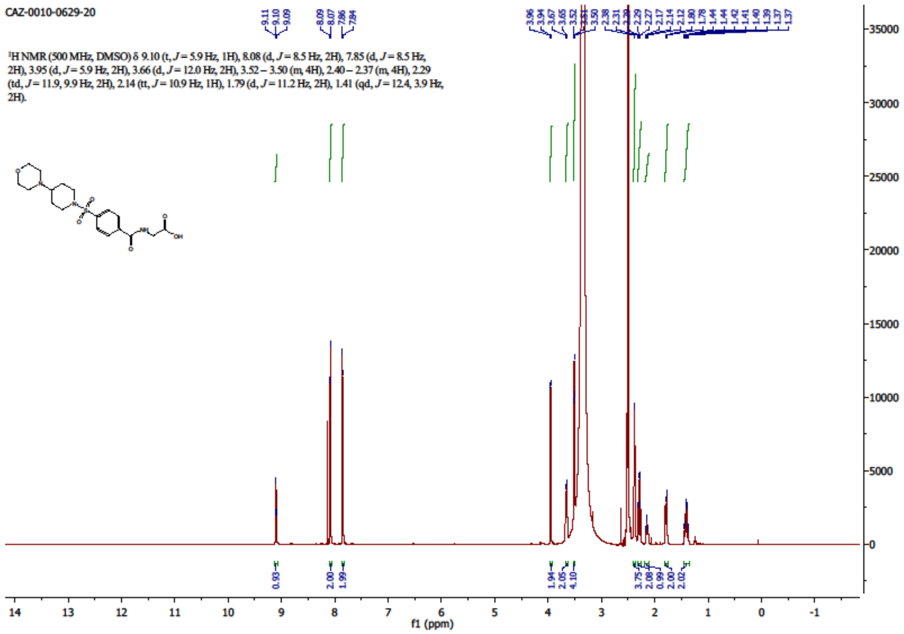


**[25]**

$^1$H NMR (500 MHz, DMSO-*d6*)

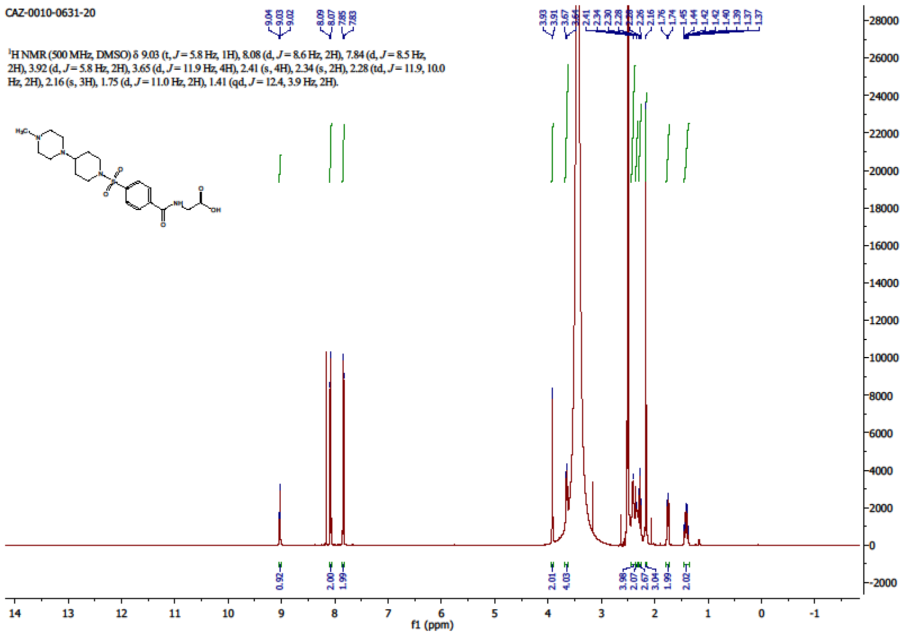

**[26]**

$^1$H NMR (500 MHz, DMSO-*d6*)

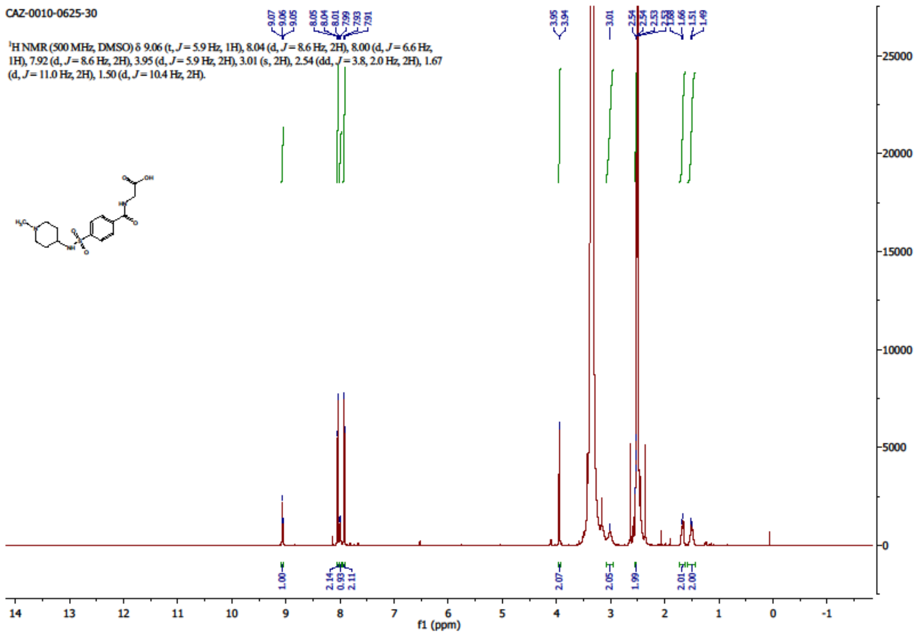


**[27]**

$^1$H NMR (500 MHz, DMSO-*d6*)

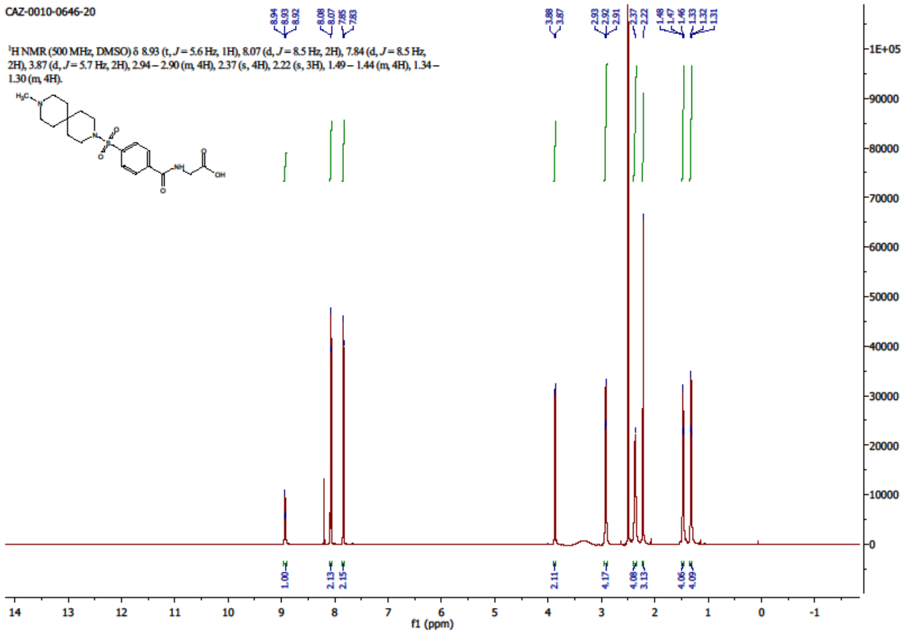

**[28]**

[1]H NMR (500 MHz, DMSO-*d6*)

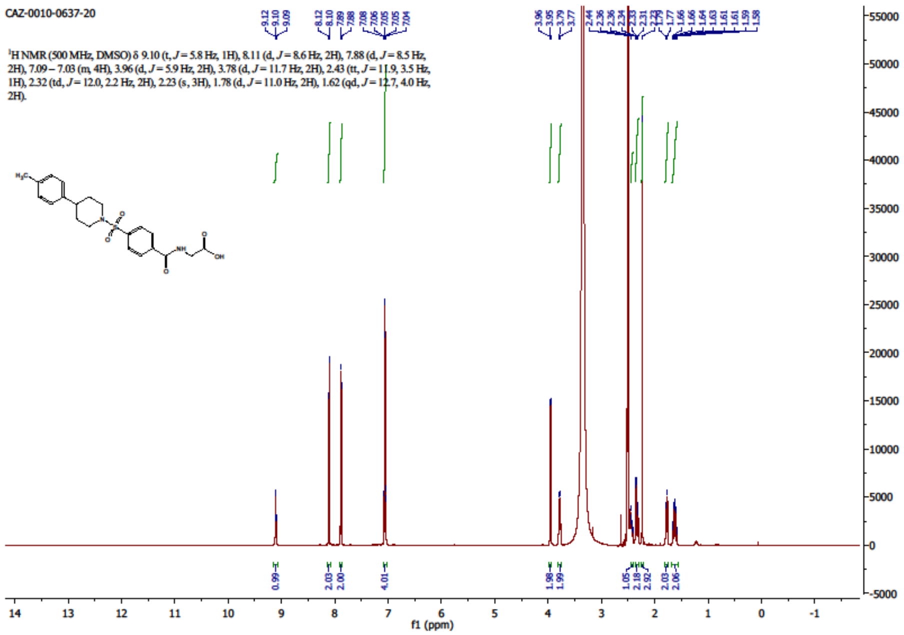


**[29]**

[1]H NMR (500 MHz, DMSO-*d6*)

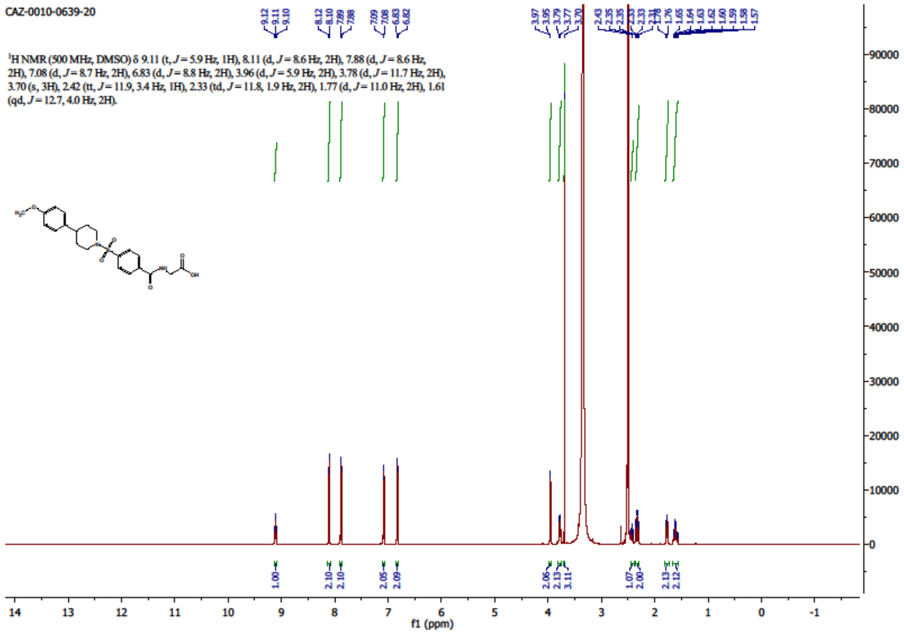

**[30]**

$^{1}$H NMR (500 MHz, DMSO-*d6*)

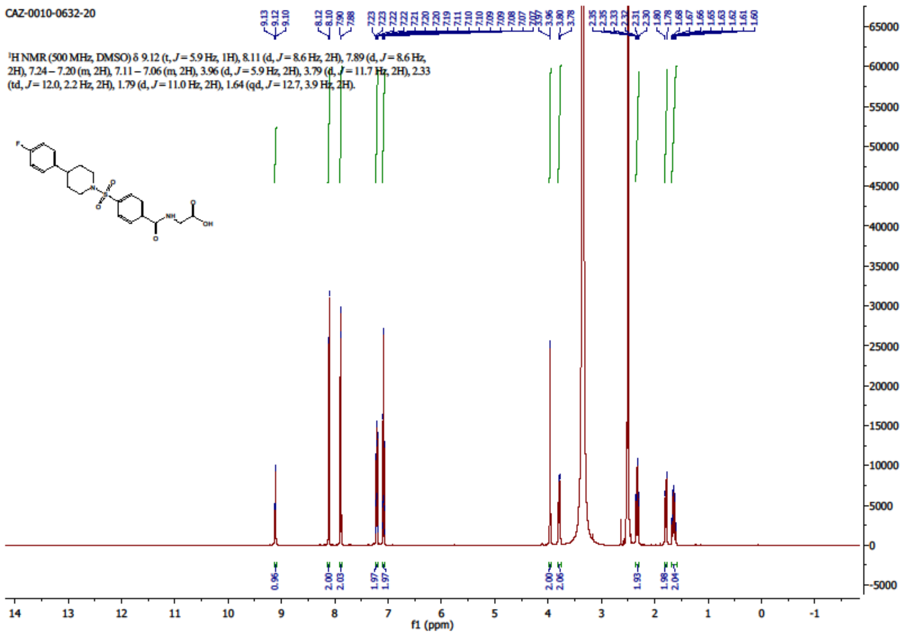

$^{19}$F NMR (471 MHz, DMSO-*d6*)

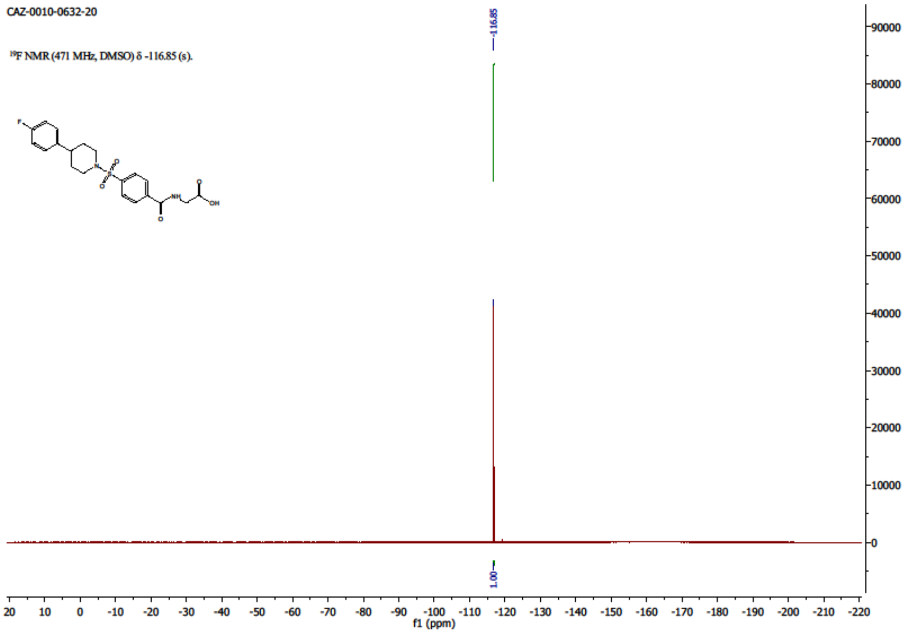

**[31]**

[1]H NMR (500 MHz, DMSO-*d6*)

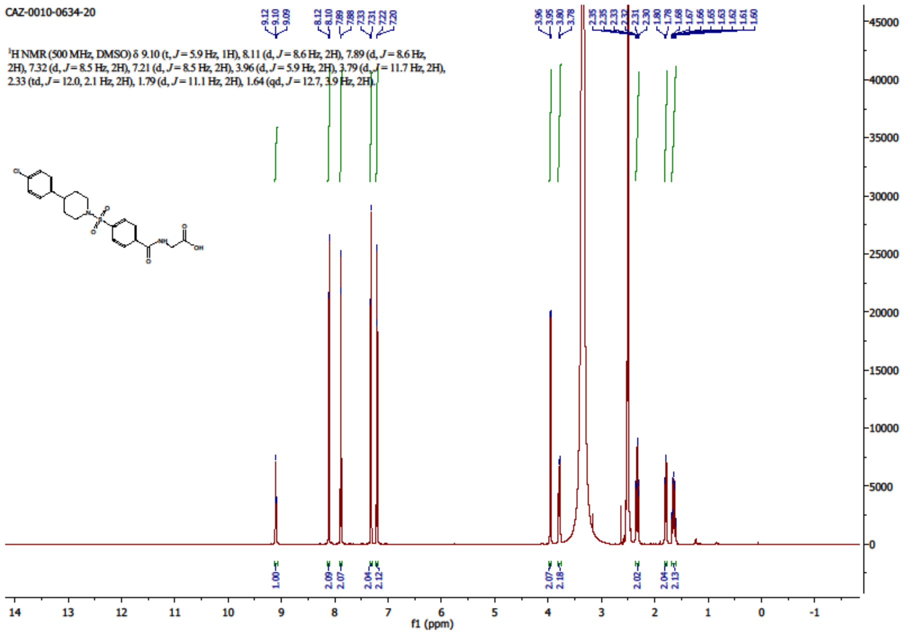


**[32]**

[1]H NMR (500 MHz, DMSO-*d6*)

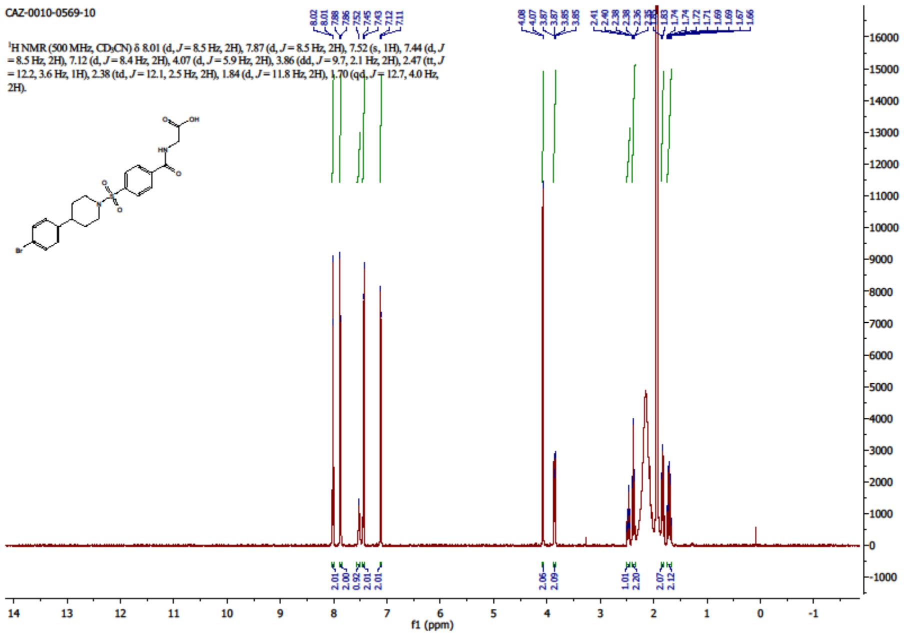

**[33]**

$^{1}$H NMR (500 MHz, DMSO-*d6*)

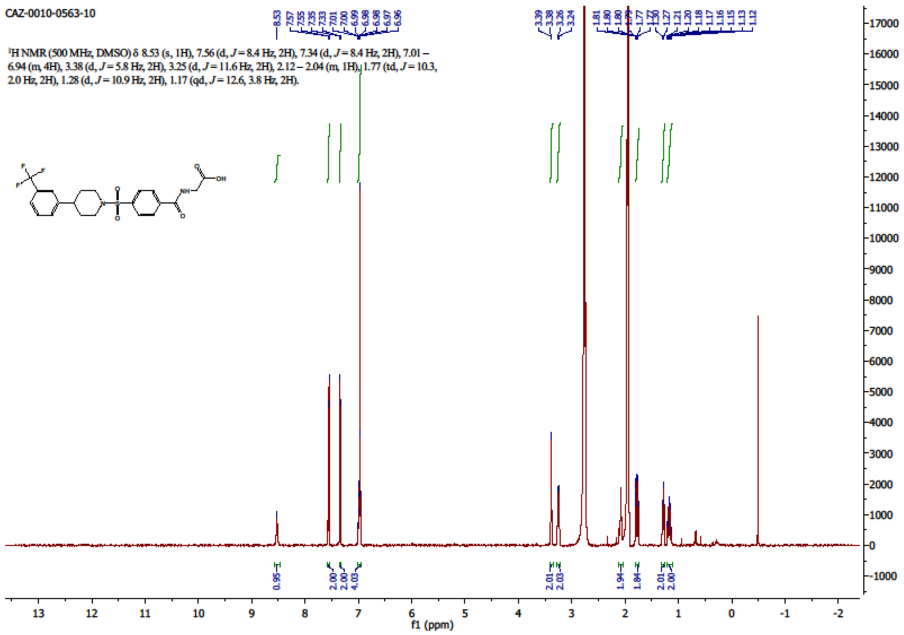


$^{19}$F NMR (471 MHz, DMSO-*d6*)

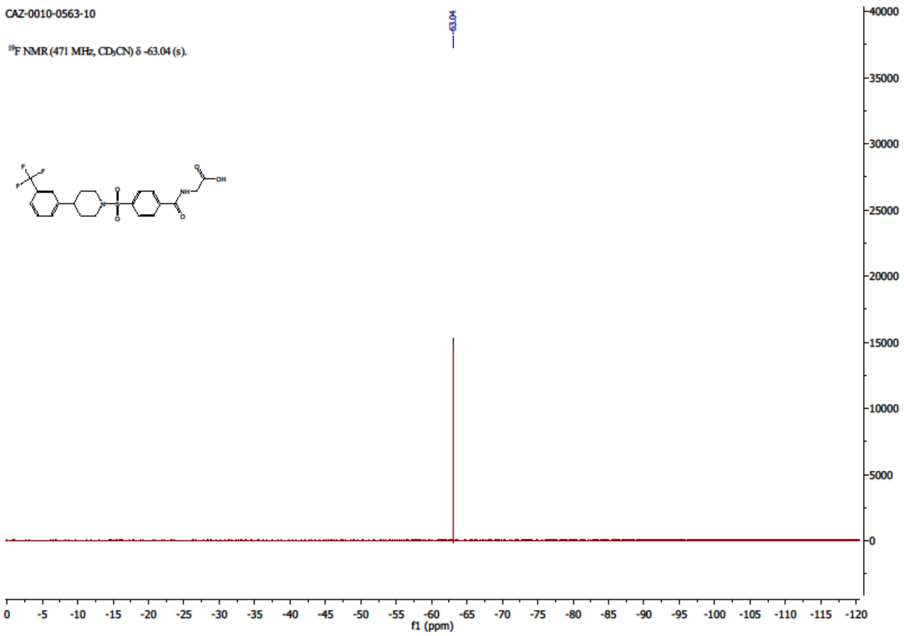

**[34]**

$^{1}$H NMR (500 MHz, DMSO-*d6*)

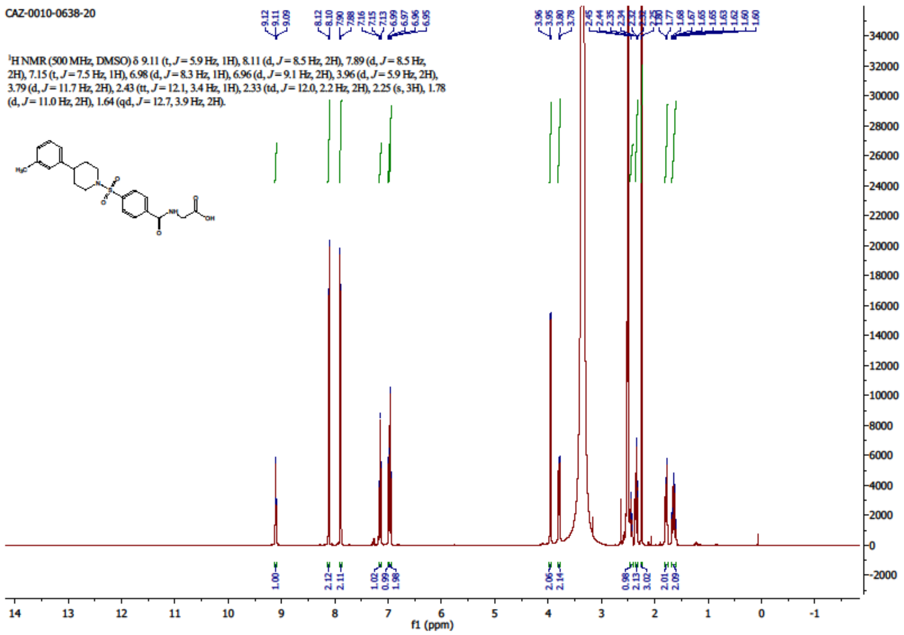


**[35]**

$^{1}$H NMR (500 MHz, DMSO-*d6*)

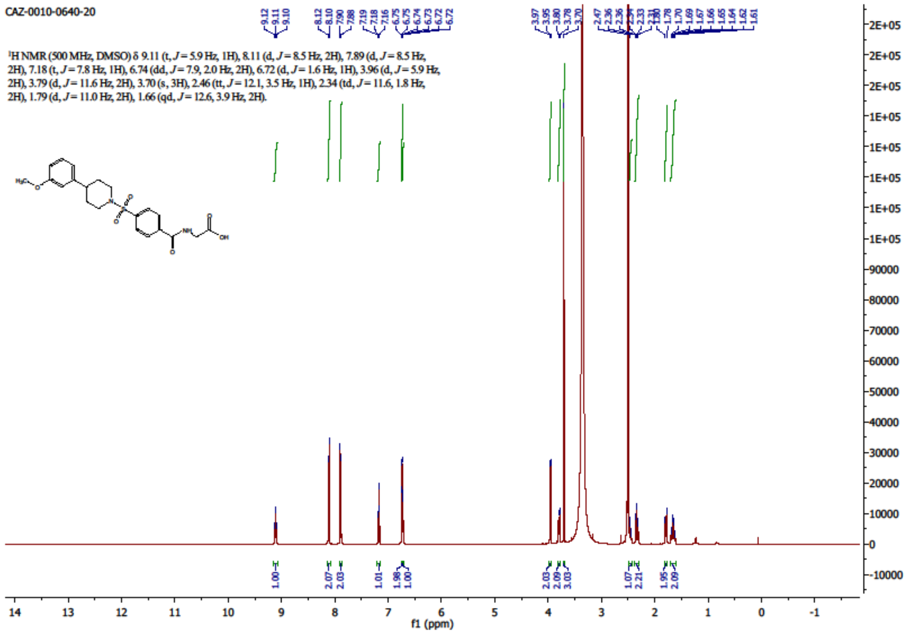

$^{1}$H NMR (500 MHz, DMSO-*d6*)

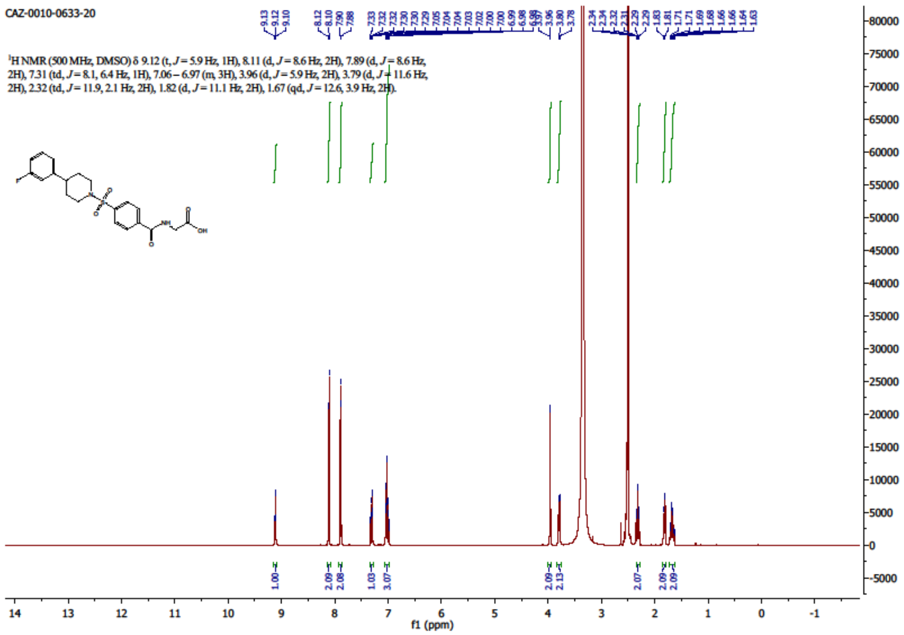

$^{19}$F NMR (471 MHz, DMSO-*d6*)

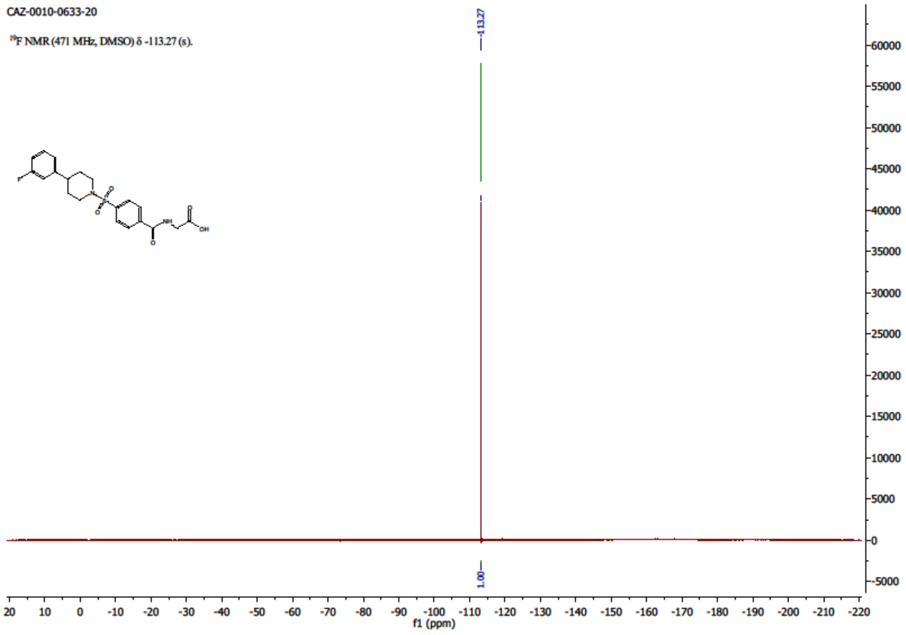

**[37]**

$^1$H NMR (500 MHz, DMSO-*d6*)

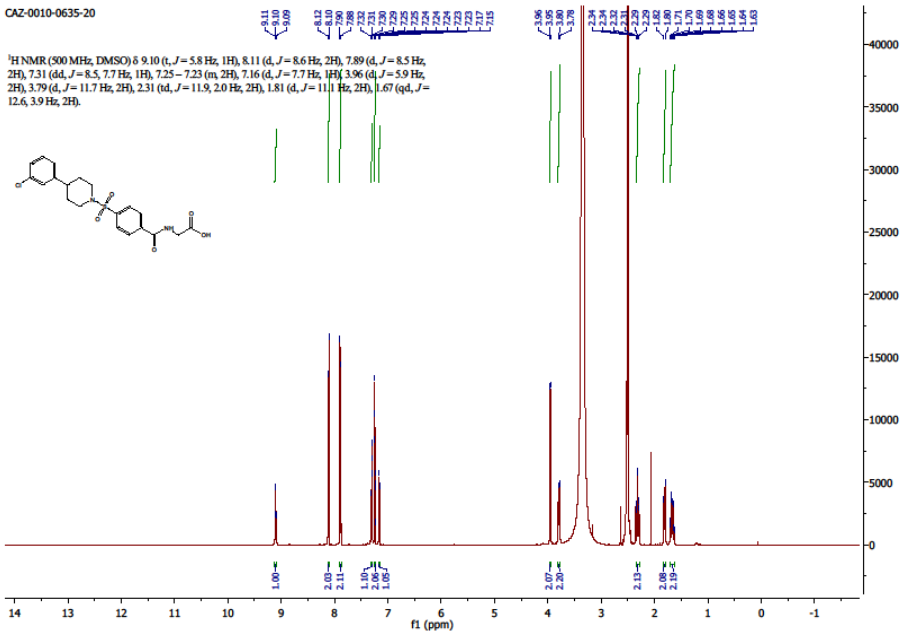

**[38]**

$^1$H NMR (500 MHz, DMSO-*d6*)

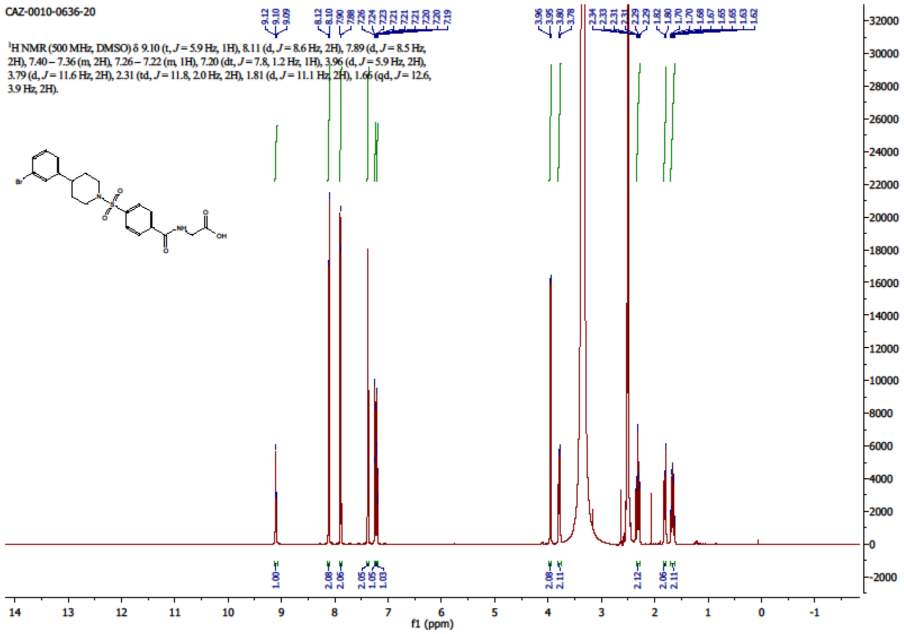

**[44]**

[1]H NMR (500 MHz, DMSO-*d6*)

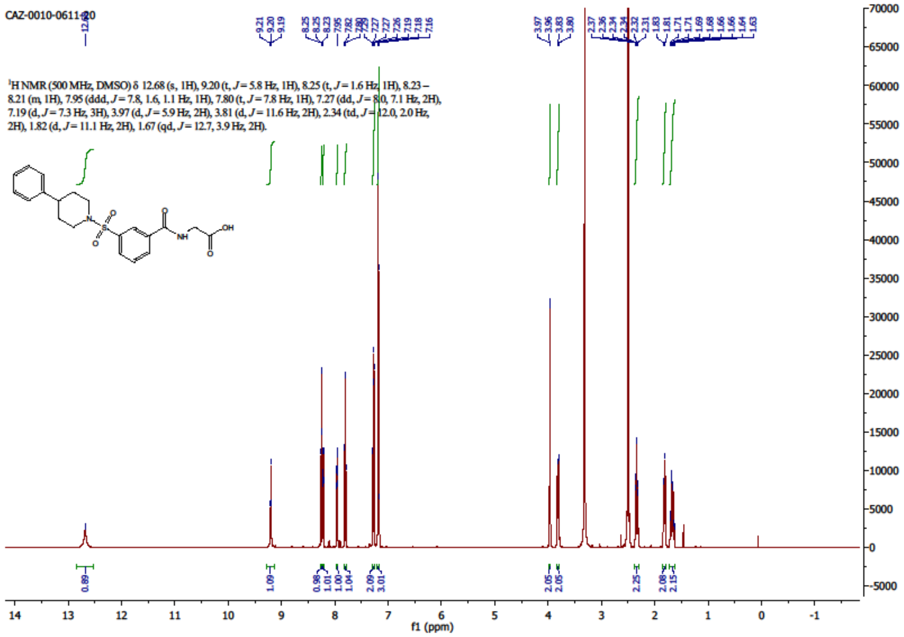

**[45]**

$^{1}$H NMR (500 MHz, DMSO-*d6*)

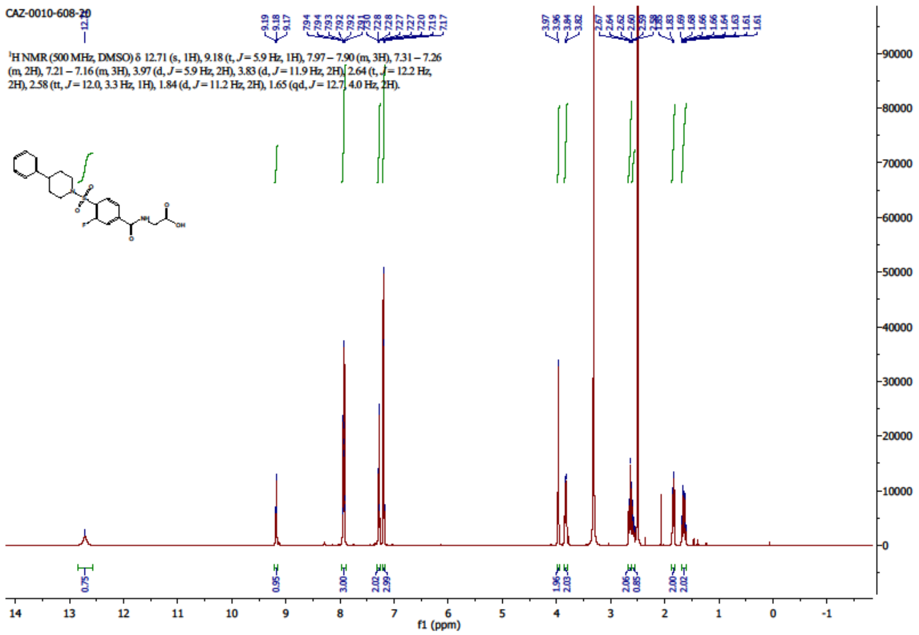


$^{19}$F NMR (471 MHz, DMSO-*d6*)

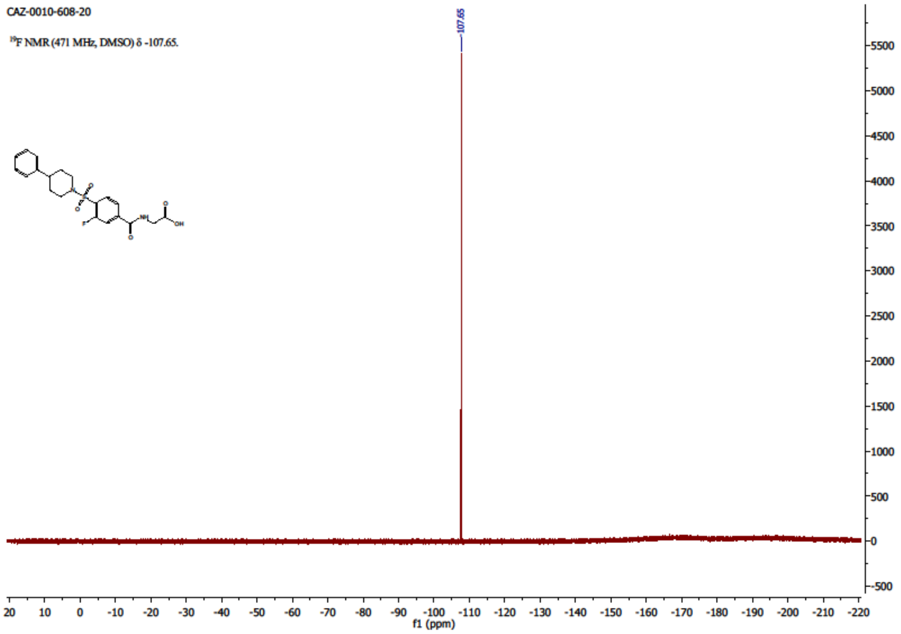

**[46]**

$^{1}$H NMR (500 MHz, DMSO-*d6*)

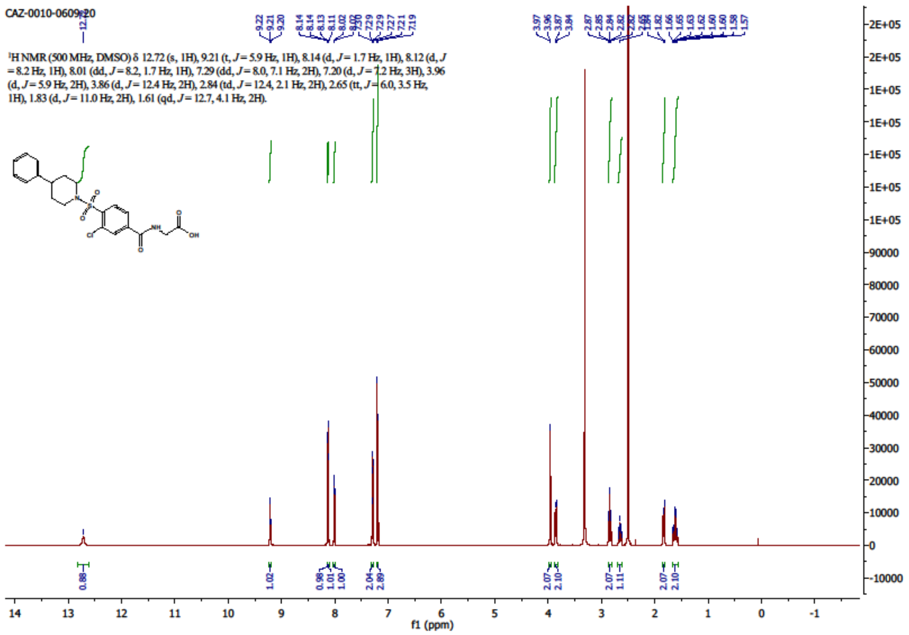

**[47]**

$^{1}$H NMR (500 MHz, DMSO-*d6*)

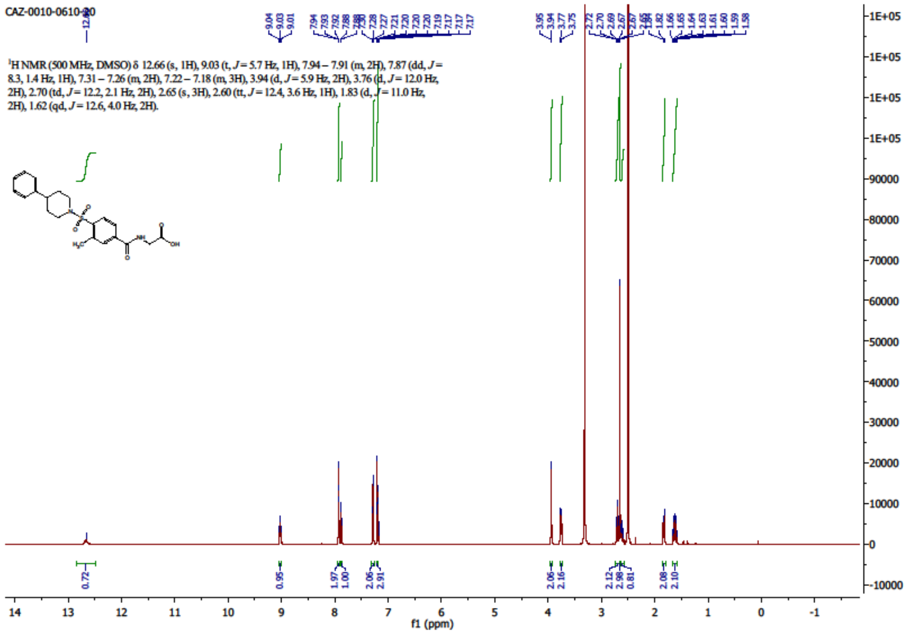

**[48]**

[1]H NMR (500 MHz, DMSO-*d6*)

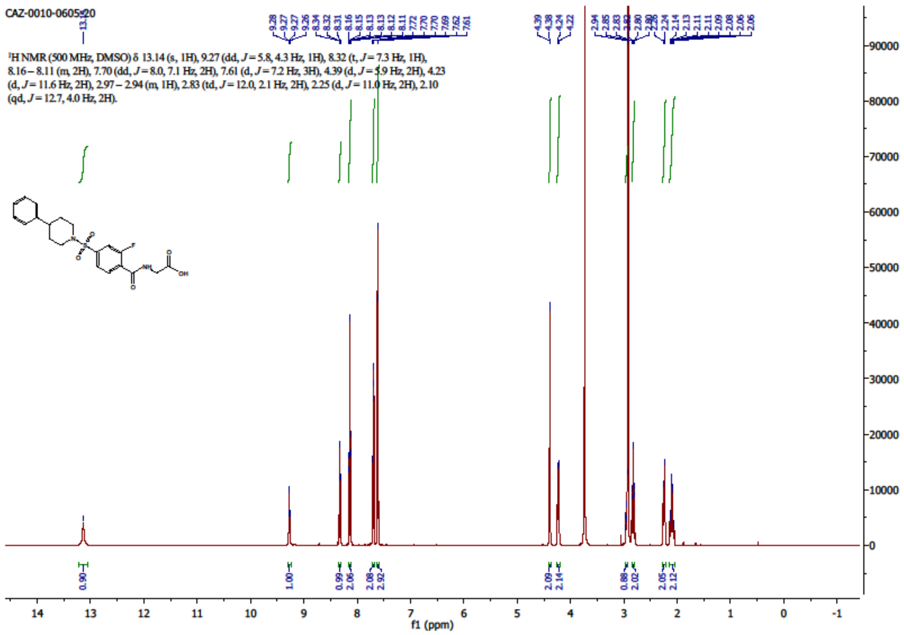


[13]C NMR (126 MHz, DMSO-*d6*)

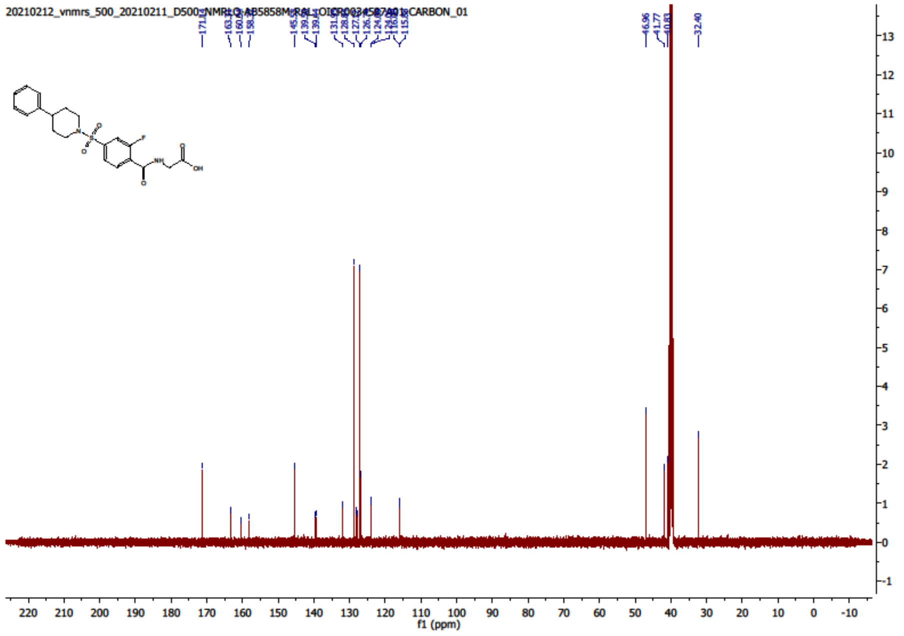

$^{19}$F NMR (471 MHz, DMSO-*d6*)

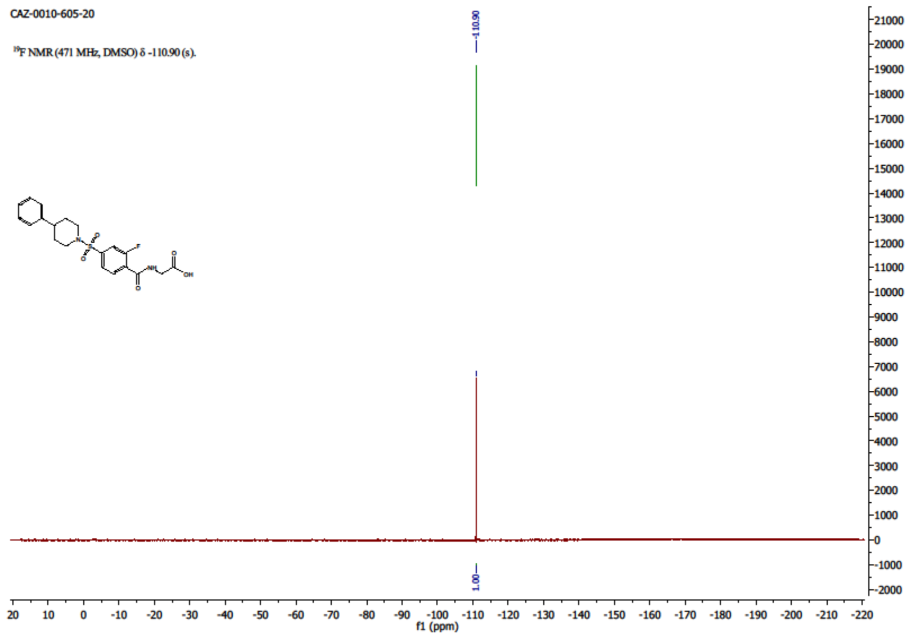


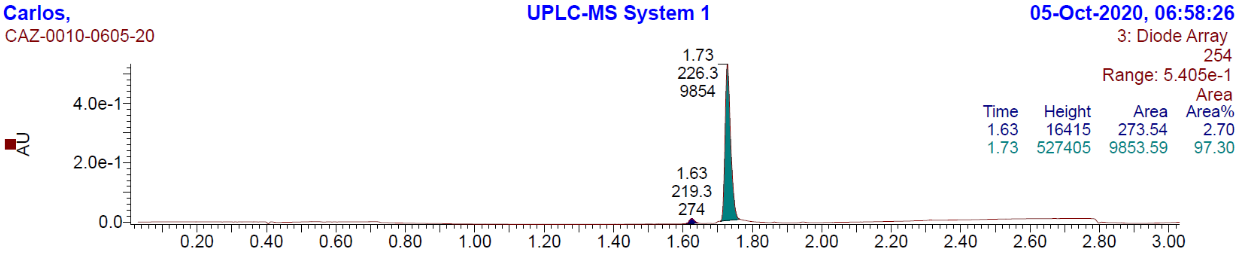

**[49]**

[1]H NMR (500 MHz, DMSO-*d6*)

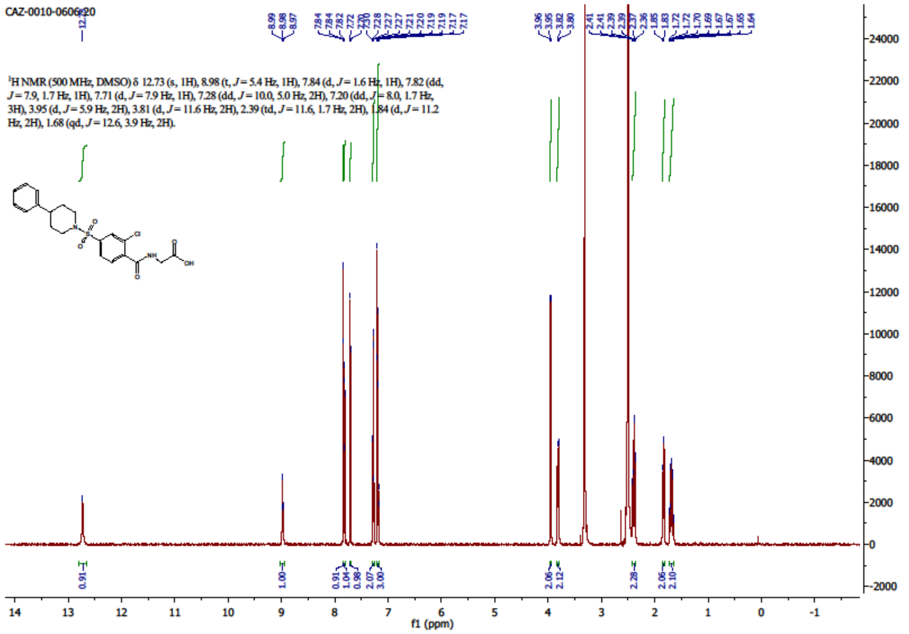

**[50]**

[1]H NMR (500 MHz, DMSO-*d6*)

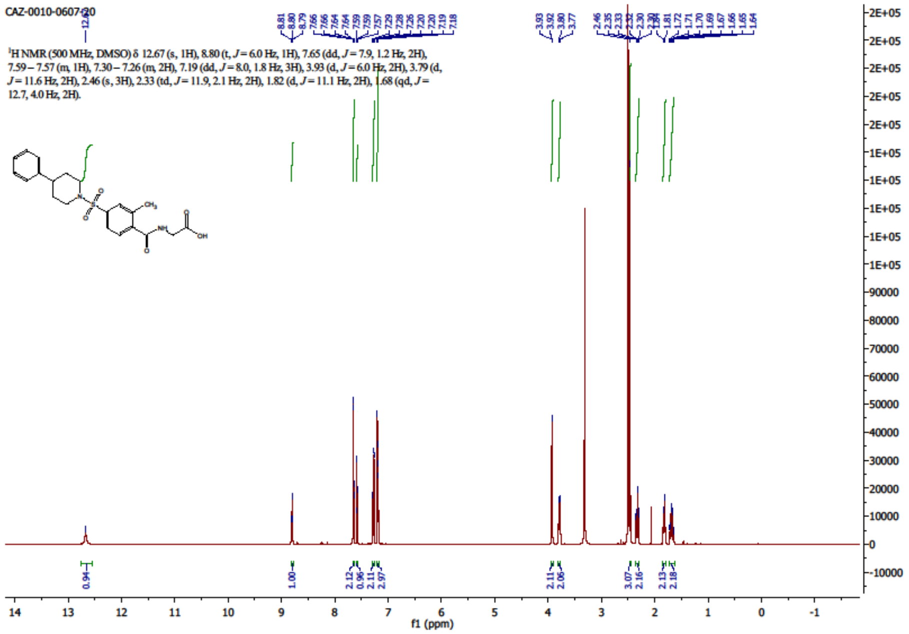

**[51]**

$^{1}$H NMR (500 MHz, DMSO-*d6*)

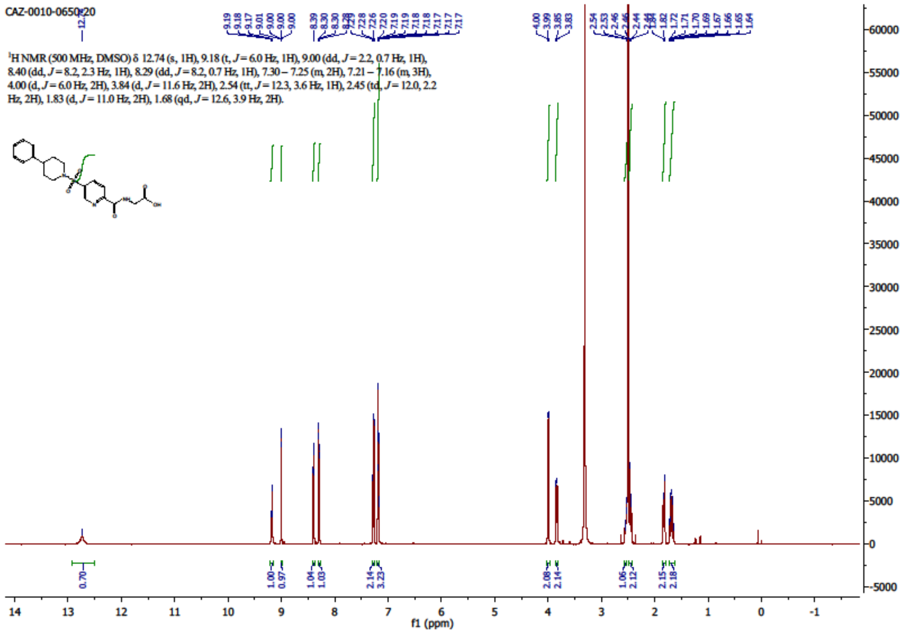


$^{13}$C NMR (126 MHz, DMSO-*d6*)

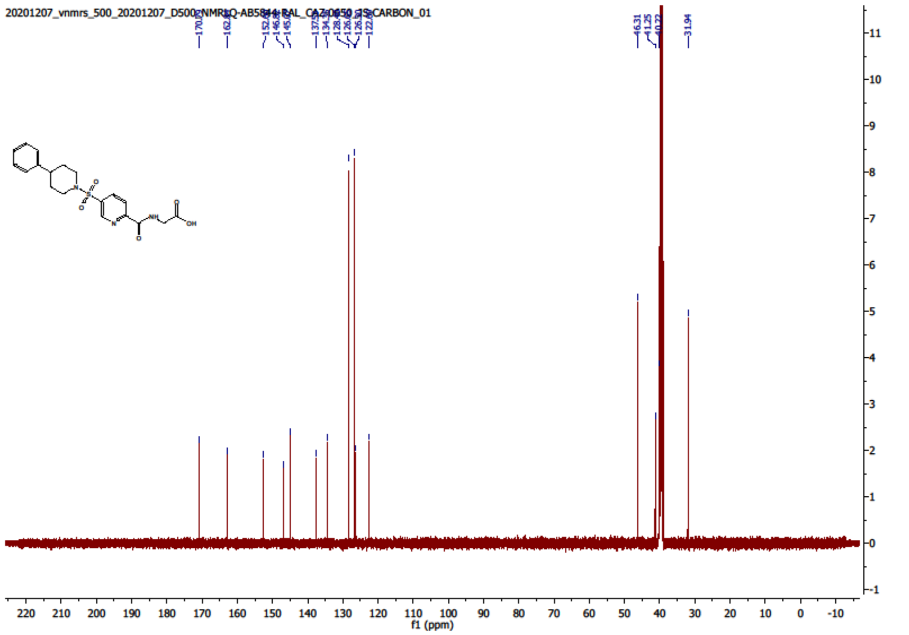

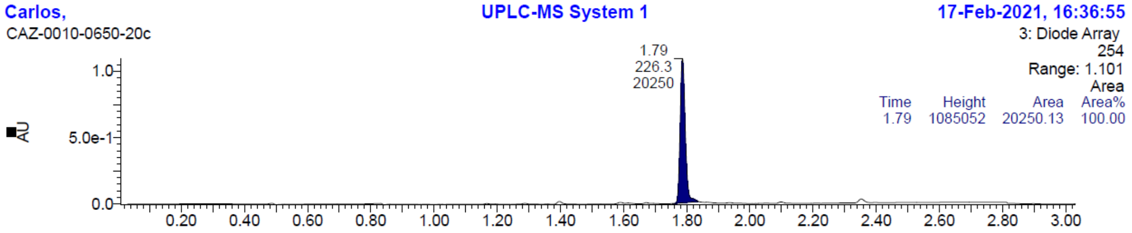


**[52]**

$^{1}$H NMR (500 MHz, DMSO-*d6*)

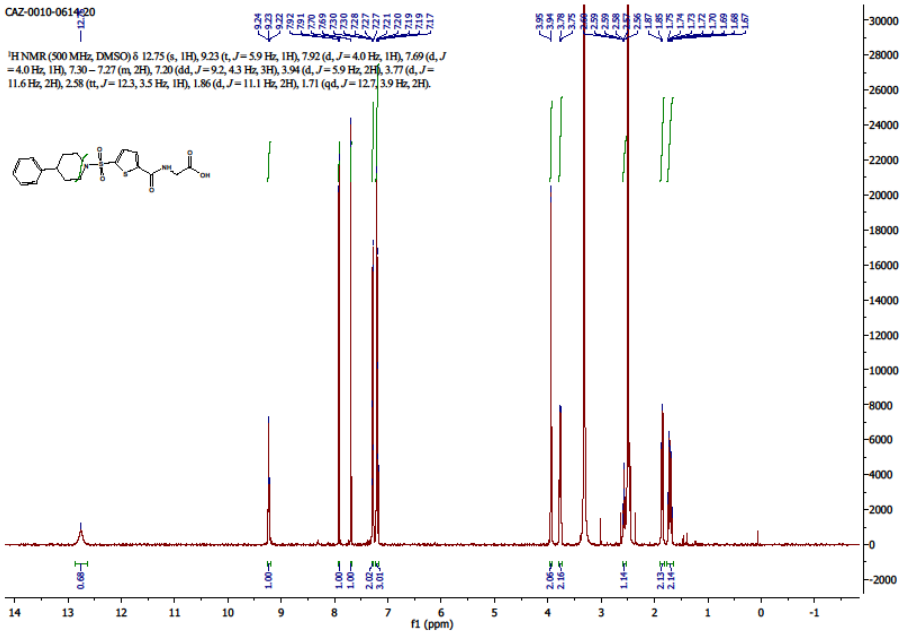


**[53]**

$^{1}$H NMR (500 MHz, DMSO-*d6*)

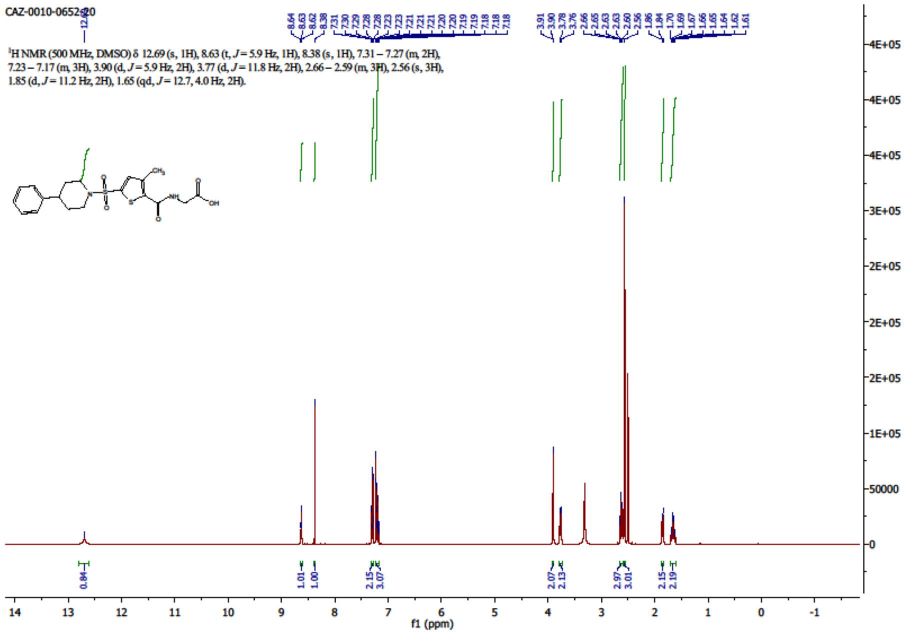

**[54]**

$^{1}$H NMR (500 MHz, DMSO-*d6*)

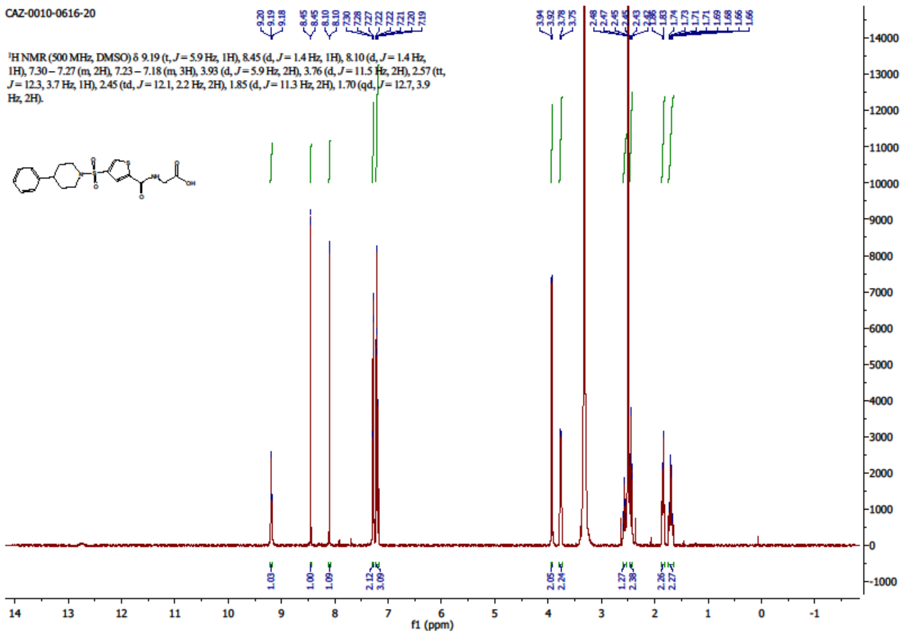

**[55]**

$^{1}$H NMR (500 MHz, DMSO-*d6*)

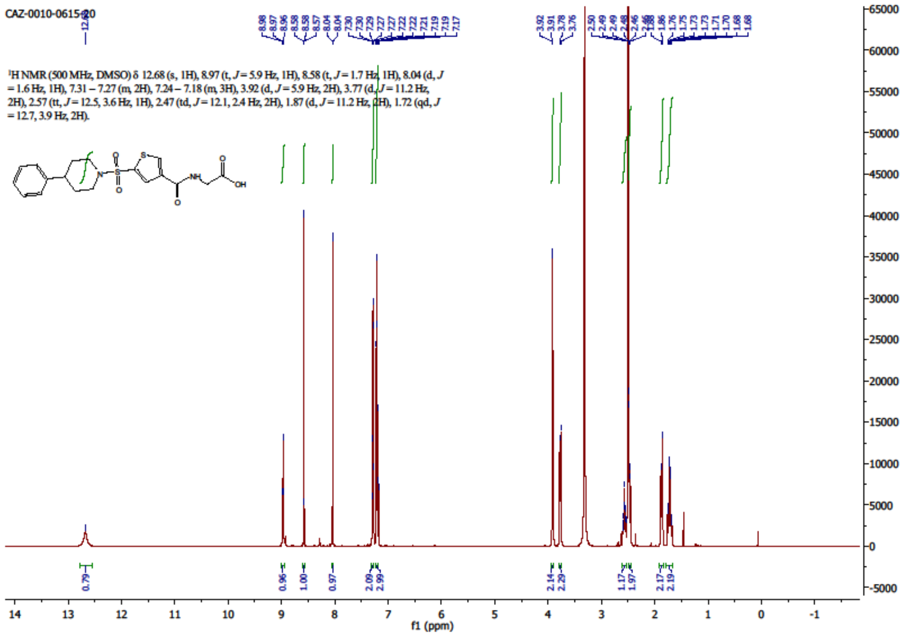

## [56]

$^{1}$H NMR (500 MHz, DMSO-*d6*)

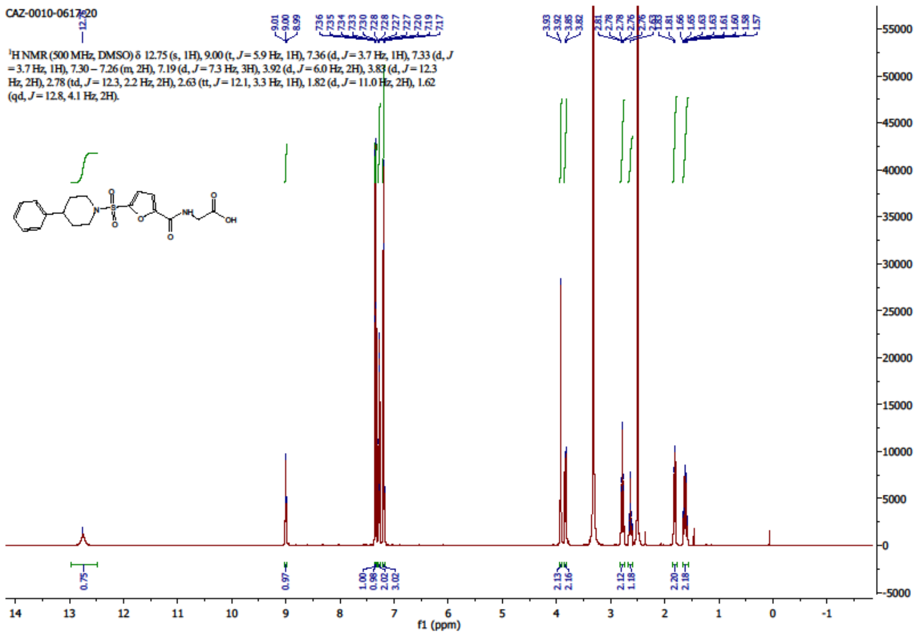


## [57]

$^{1}$H NMR (500 MHz, DMSO-*d6*)

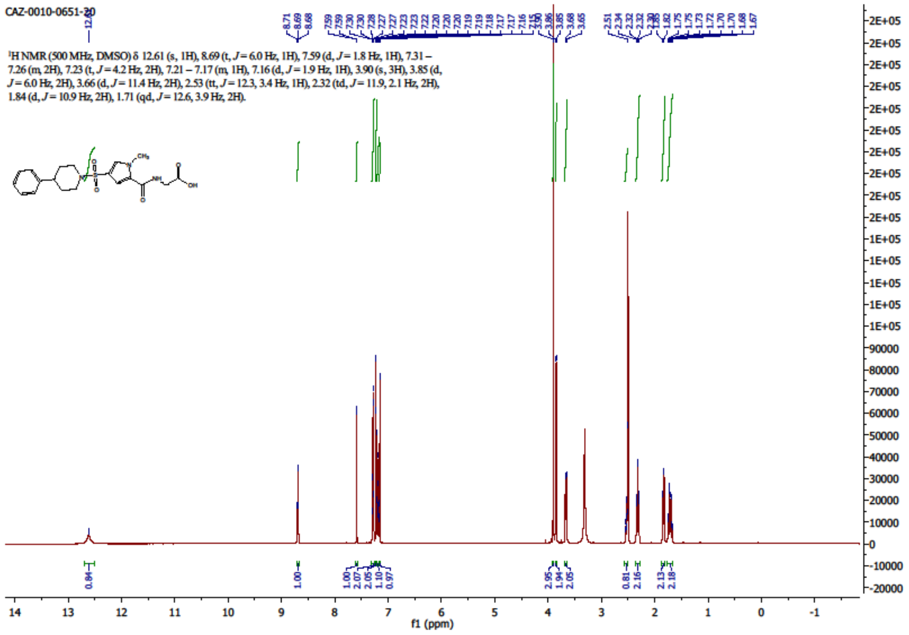

# [61]

[1]H NMR (500 MHz, DMSO-*d6*)

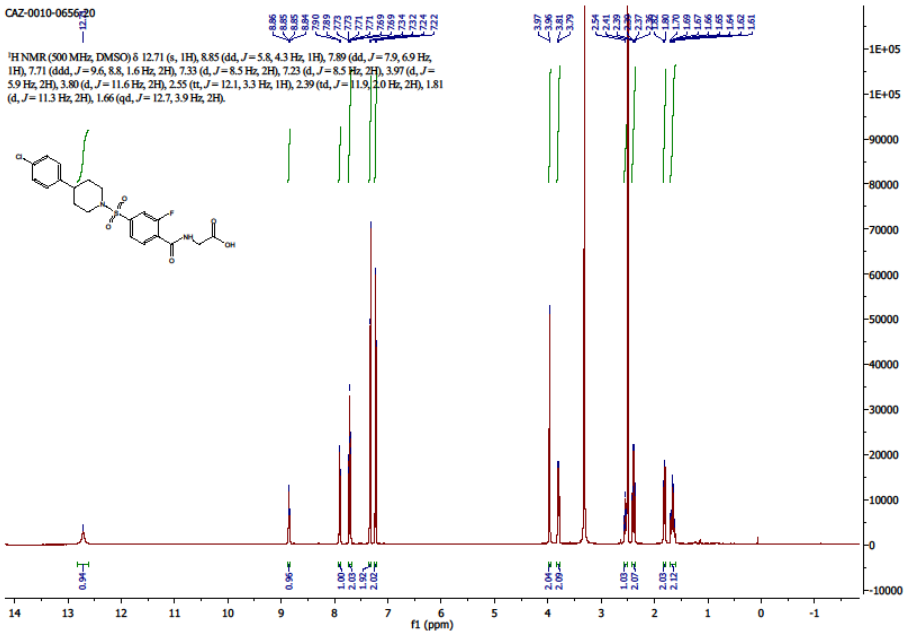


[13]C NMR (126 MHz, DMSO-*d6*)

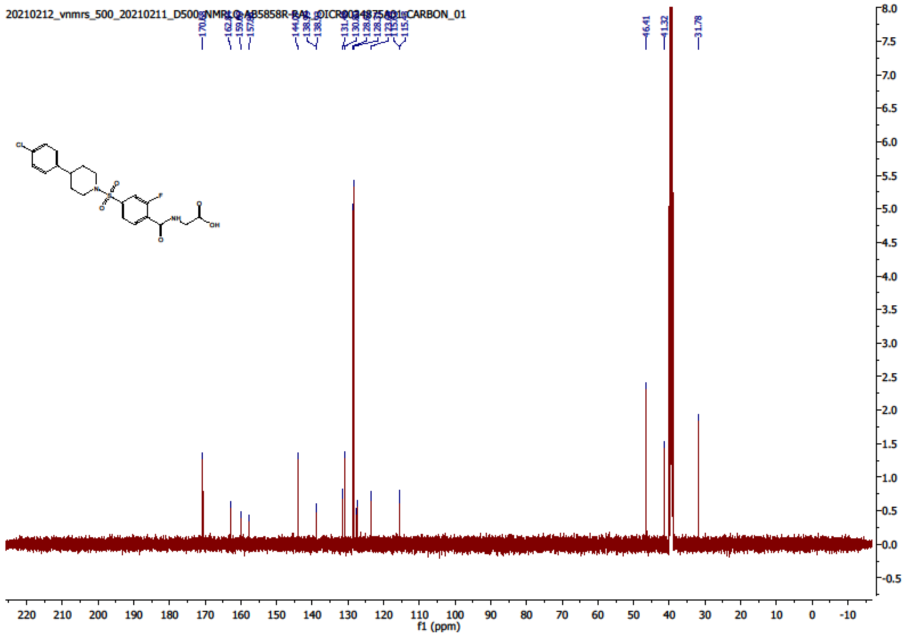

$^{19}F$ NMR (471 MHz, DMSO-*d6*)

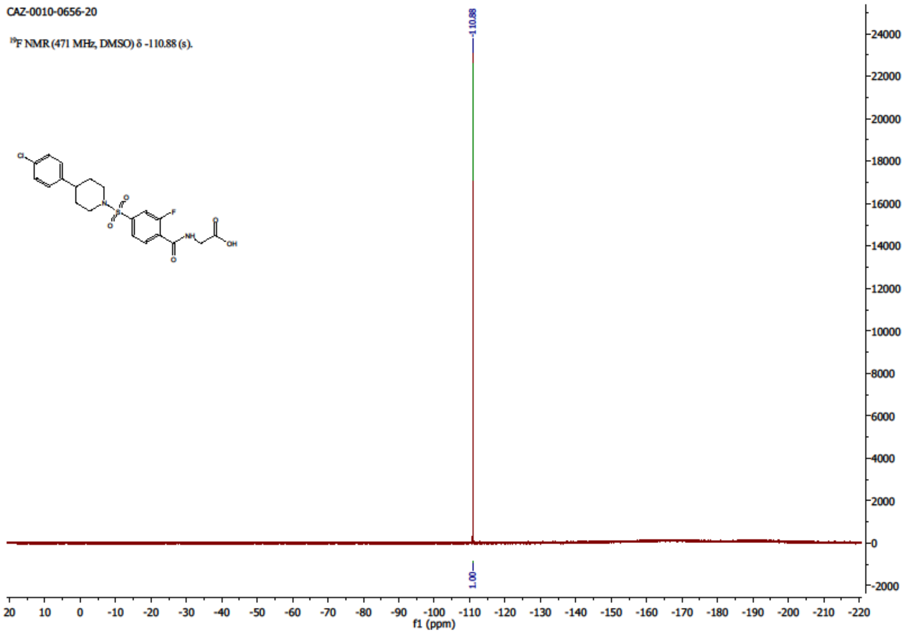


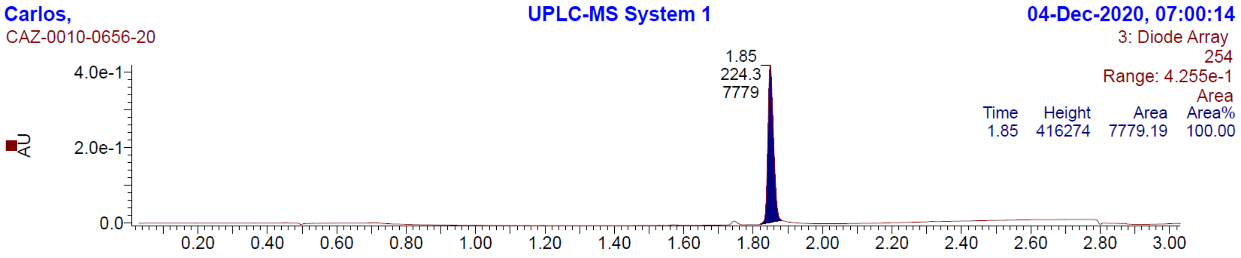

# [62]

$^{1}$H NMR (500 MHz, DMSO-*d6*)

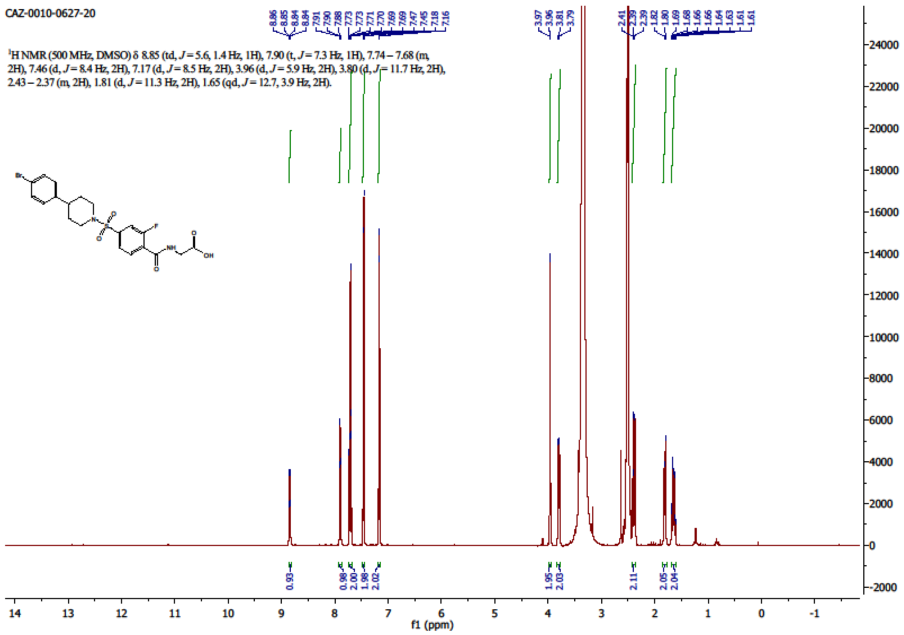


$^{13}$C NMR (126 MHz, DMSO-*d6*)

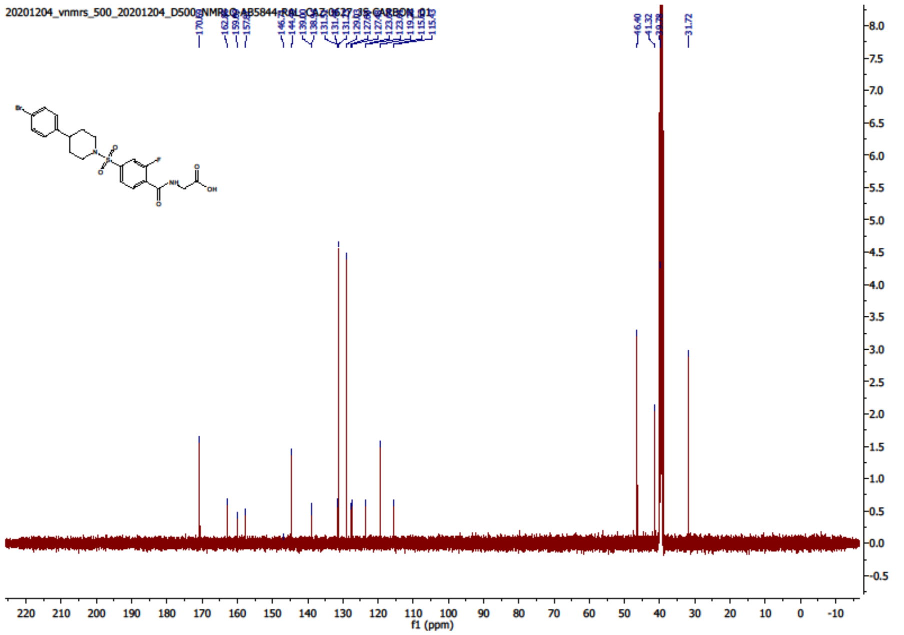

$^{19}$F NMR (471 MHz, DMSO-*d6*)

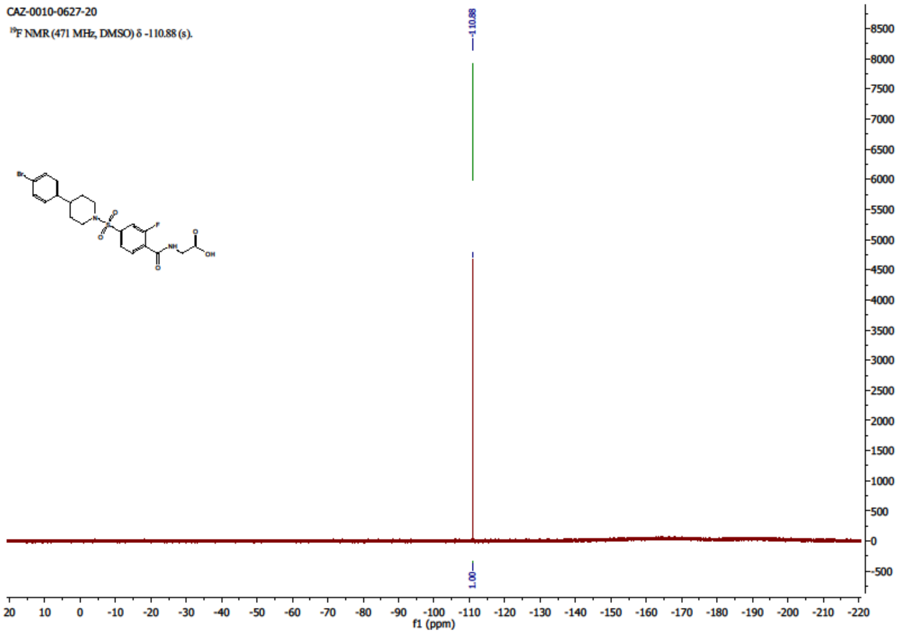


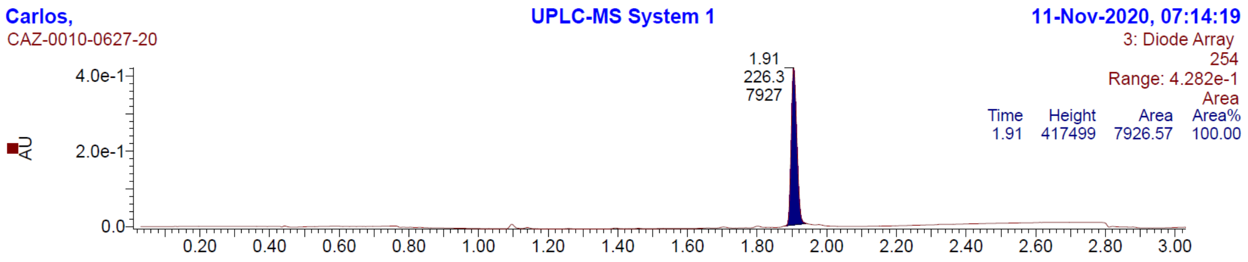

## [63]

$^{1}$H NMR (500 MHz, DMSO-*d6*)

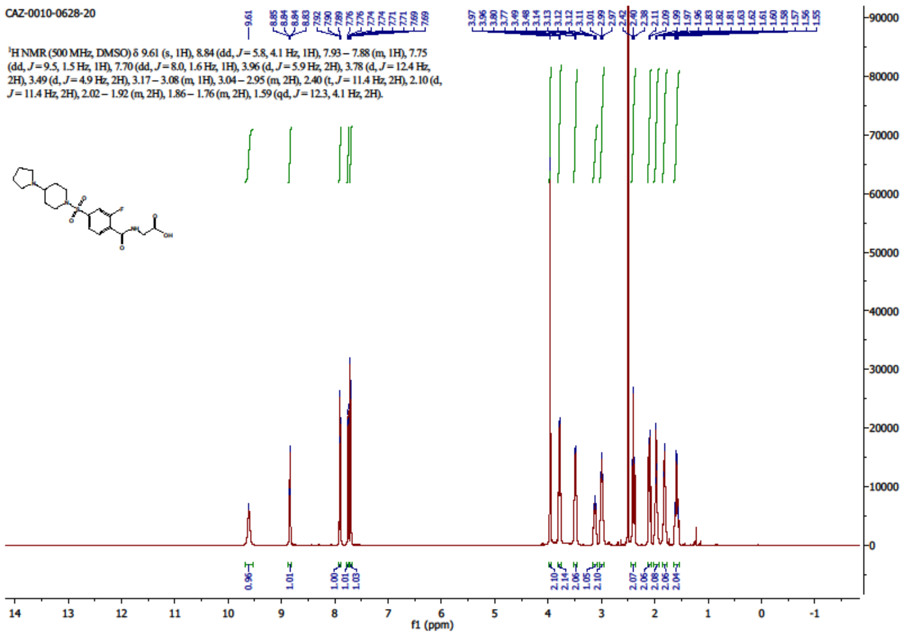


$^{13}$C NMR (126 MHz, DMSO-*d6*)

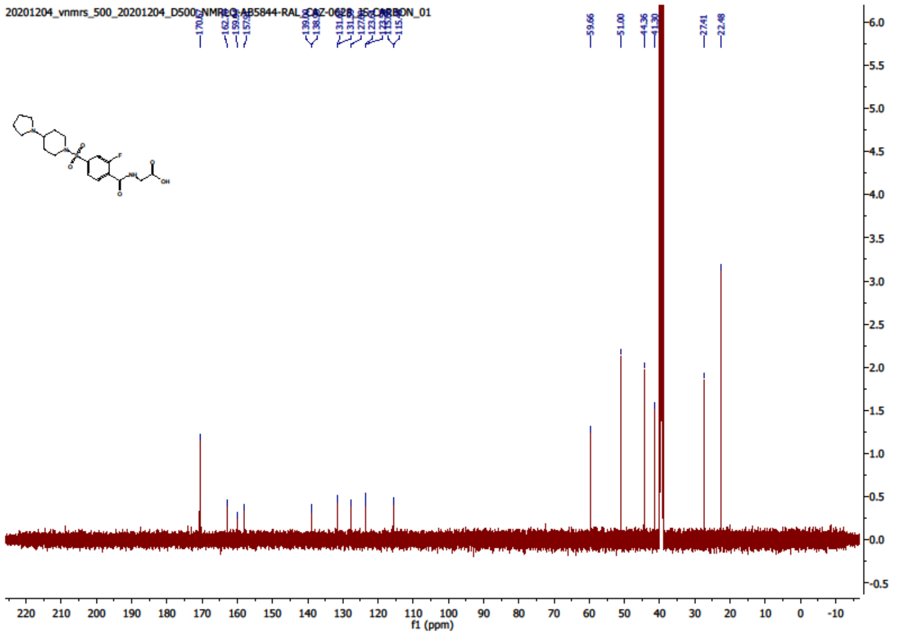

$^{19}$F NMR (471 MHz, DMSO-*d6*)

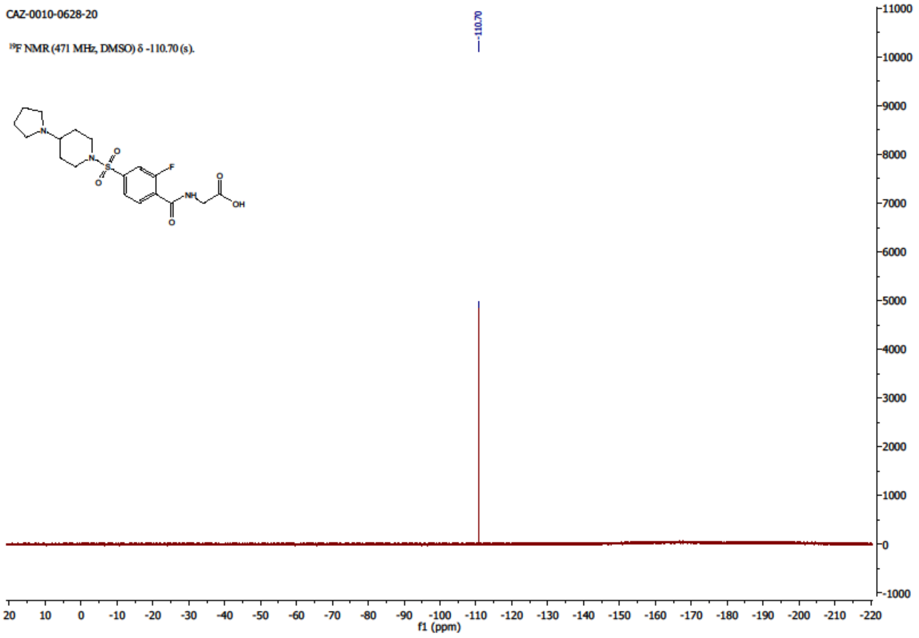


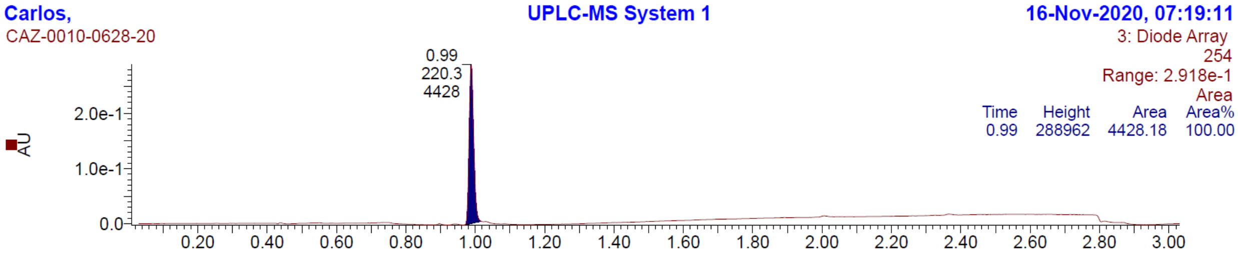

**[64]**

[1]H NMR (500 MHz, DMSO-*d6*)

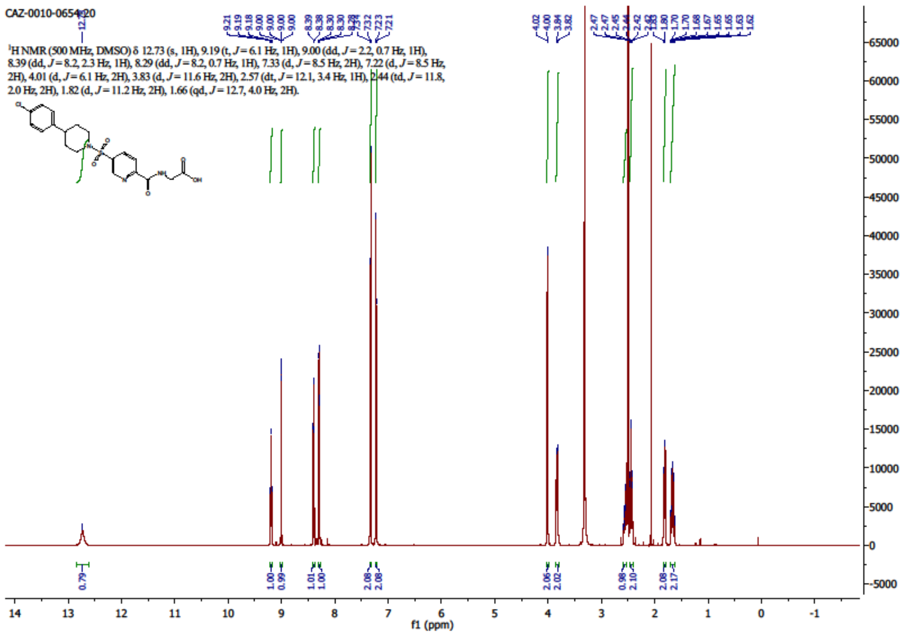


[13]C NMR (126 MHz, DMSO-*d6*)

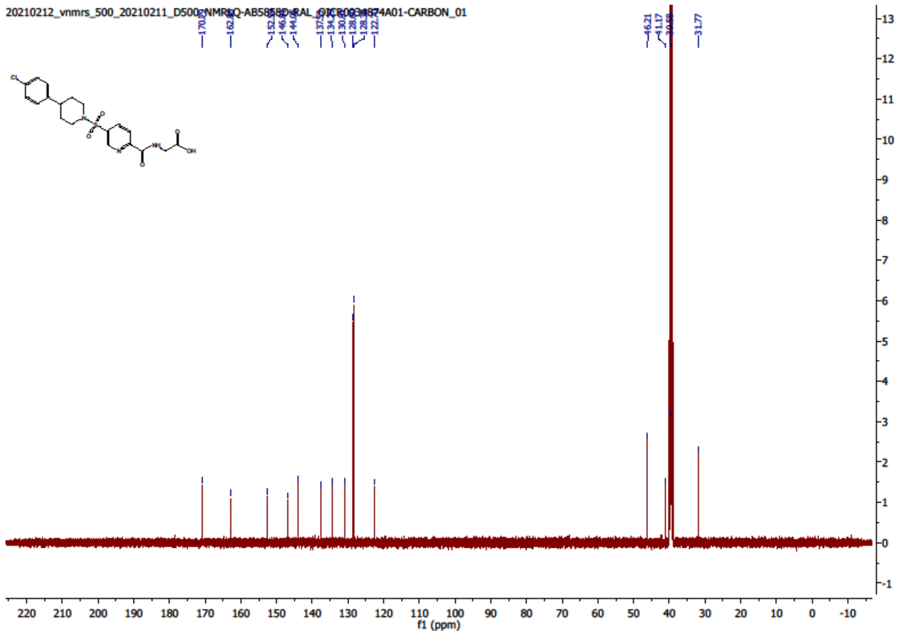

Carlos, UPLC-MS System 1 04-Dec-2020, 06:52:10

CAZ-0010-0654-20

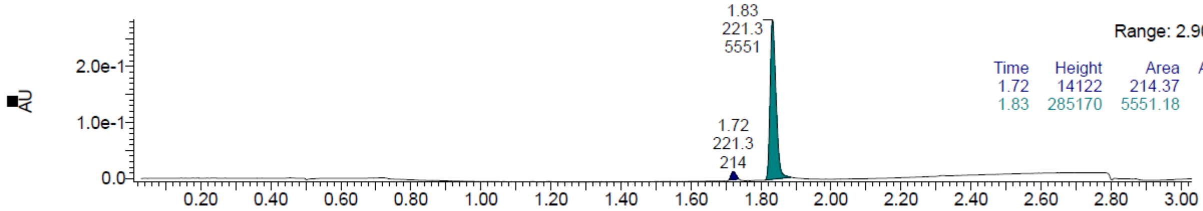


| Time | Height | Area | Area% |
| --- | --- | --- | --- |
| 1.72 | 14122 | 214.37 | 3.72 |
| 1.83 | 285170 | 5551.18 | 96.28 |

**[S1]**

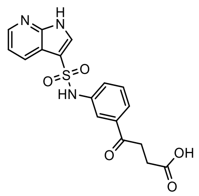

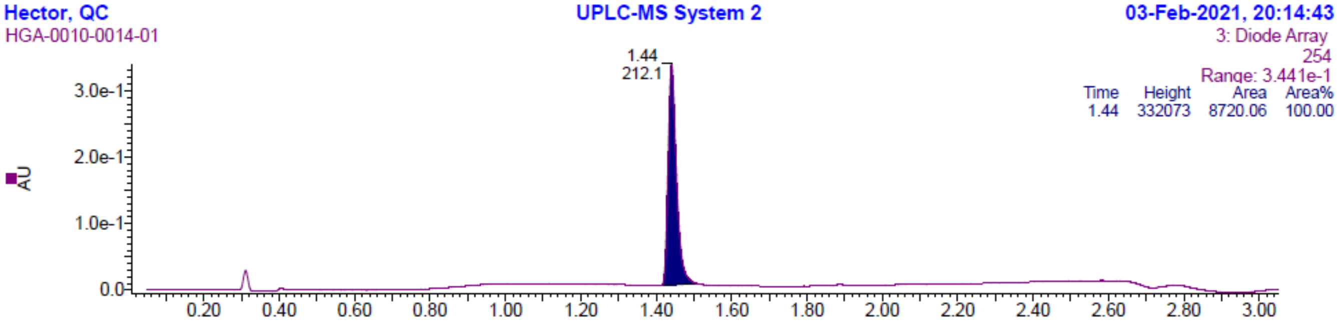


LC: Method 1, $t_R$= 1.44 min. MS (ESI): *m/z*= 373.88 $[M + H]^+$

**[S2]**

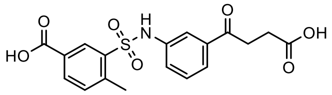

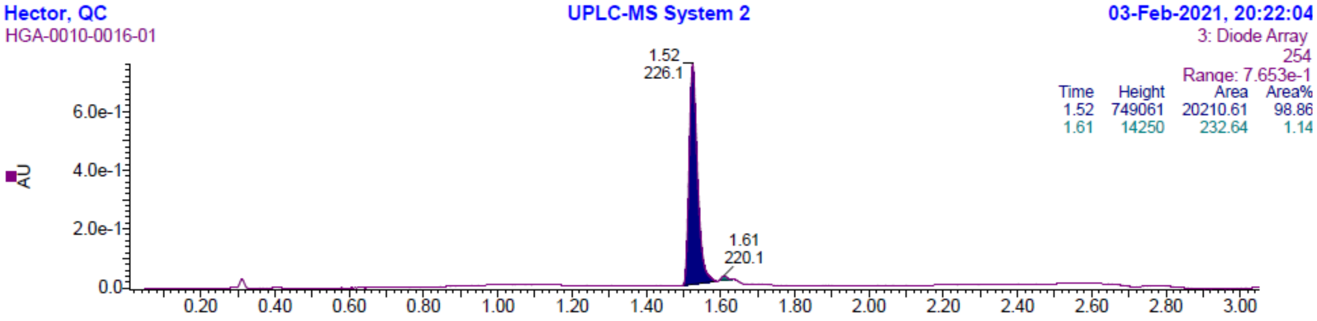


LC: Method 1, $t_R$ = 1.52 min. MS (ESI): m/z = 390.24 $[M - H]^-$

**[S3]**

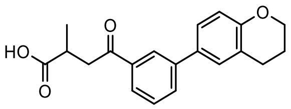

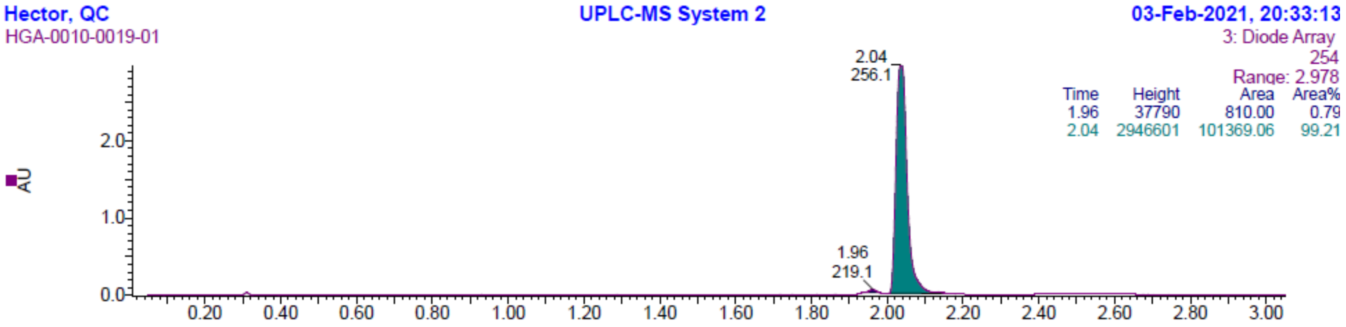


LC: Method 1, $t_R$ = 2.04 min. MS (ESI): m/z = 323.10 $[M - H]^-$

**[S4]**

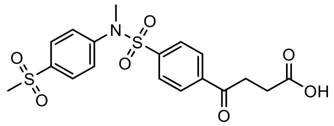

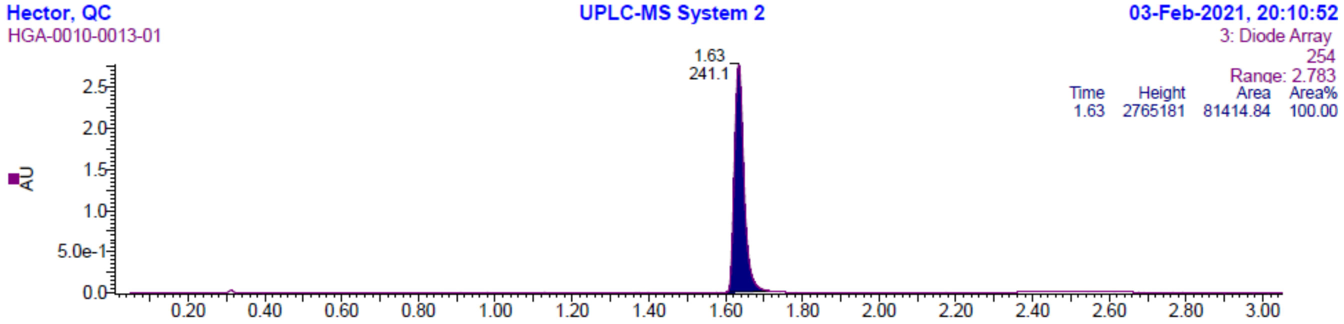


LC: Method 1, $t_R$ = 1.63 min. MS (ESI): m/z = 424.08 $[M - H]^-$

**[S5]**

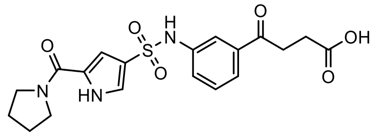

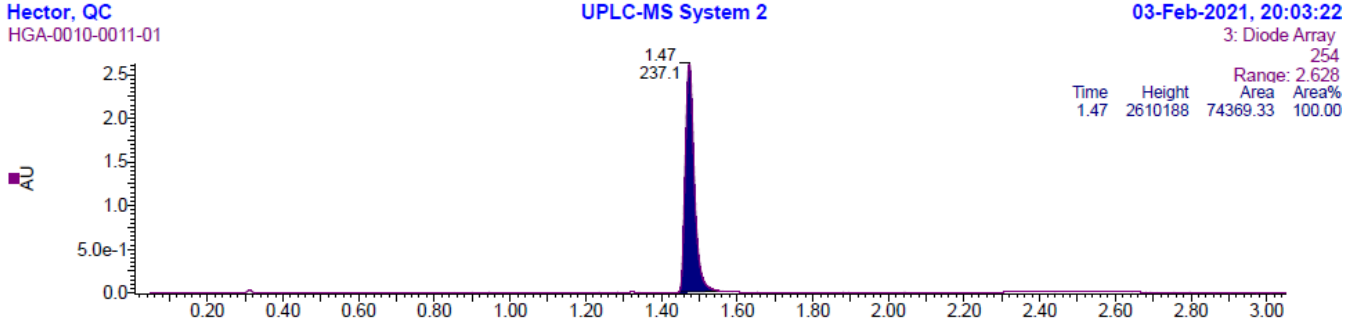


LC: Method 1, $t_R$= 1.47 min. MS (ESI): *m/z*= 420.07 $[M + H]^+$

**[S6]**

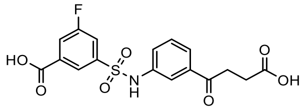

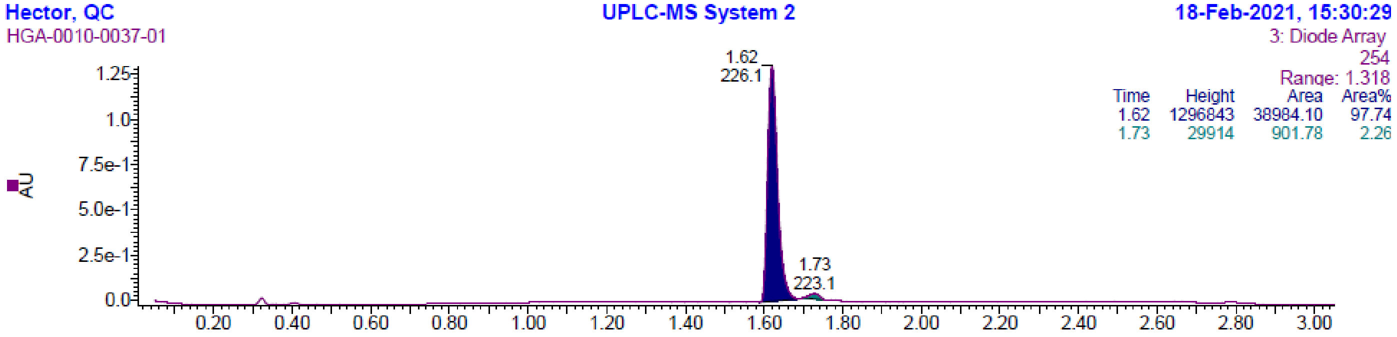


LC: Method 1, $t_R$ = 1.62 min. MS (ESI): m/z = 394.00 $[M - H]^-$

**[S7]**

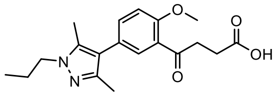

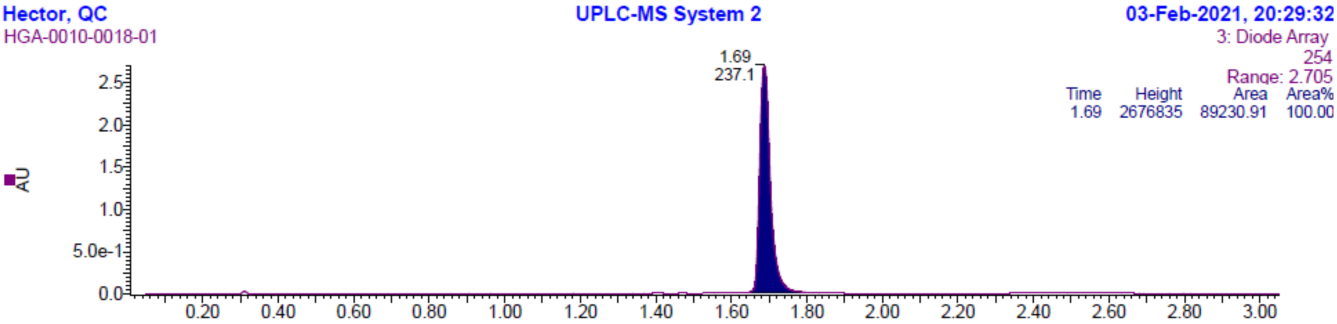


LC: Method 1, $t_R$ = 1.69 min. MS (ESI): m/z = 343.16 $[M - H]^-$

**[S8]**

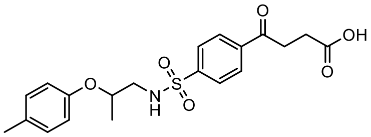

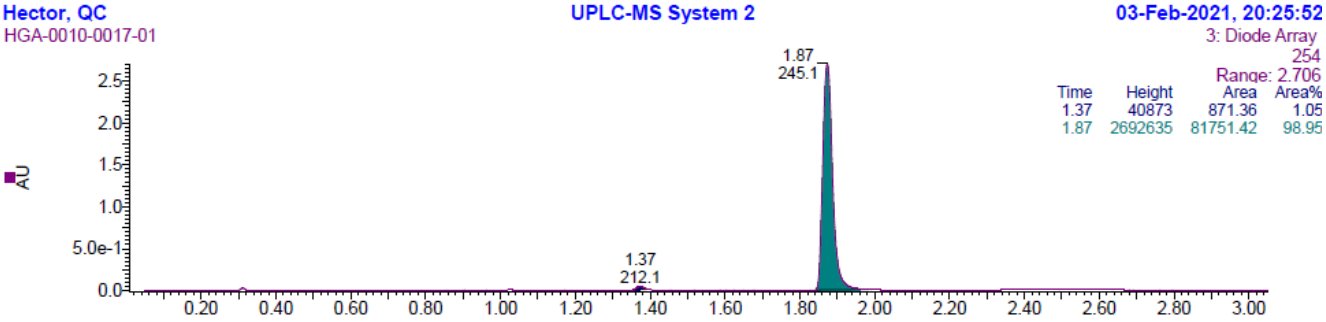


LC: Method 1, $t_R$ = 1.87 min. MS (ESI): m/z = 404.28 $[M - H]^-$

**[S9]**

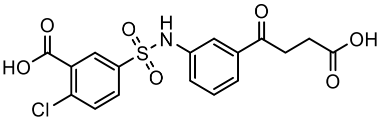

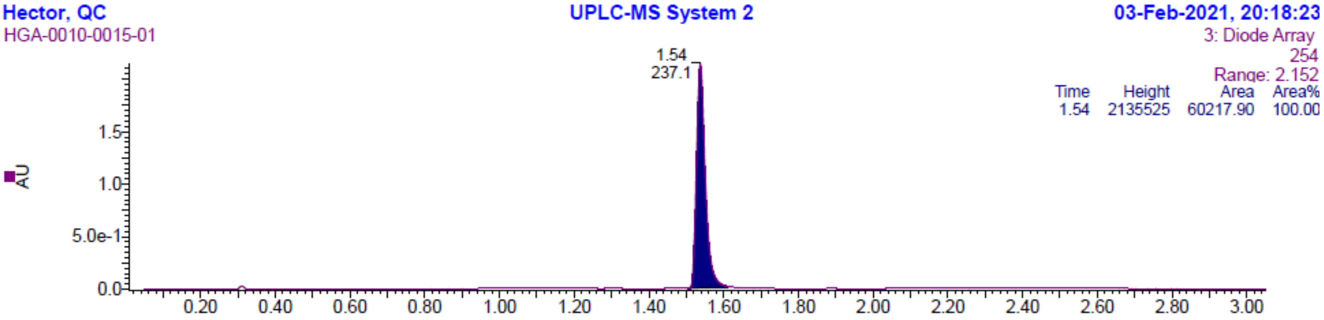


LC: Method 1, $t_R$ = 1.54 min. MS (ESI): m/z = 409.98 $[M - H]^-$, 410.17 $[M - 1]^-$ +2

**[S10]**

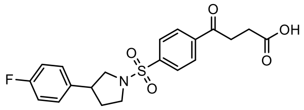


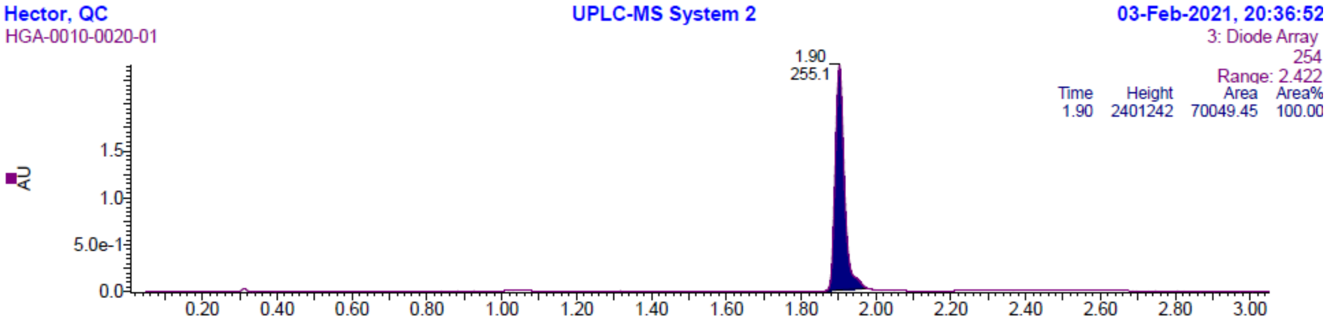


LC: Method 1, $t_R$ = 1.90 min. MS (ESI): m/z = 404.28 [M - H]$^-$

**[S11]**

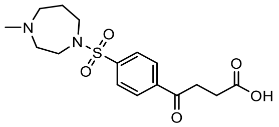


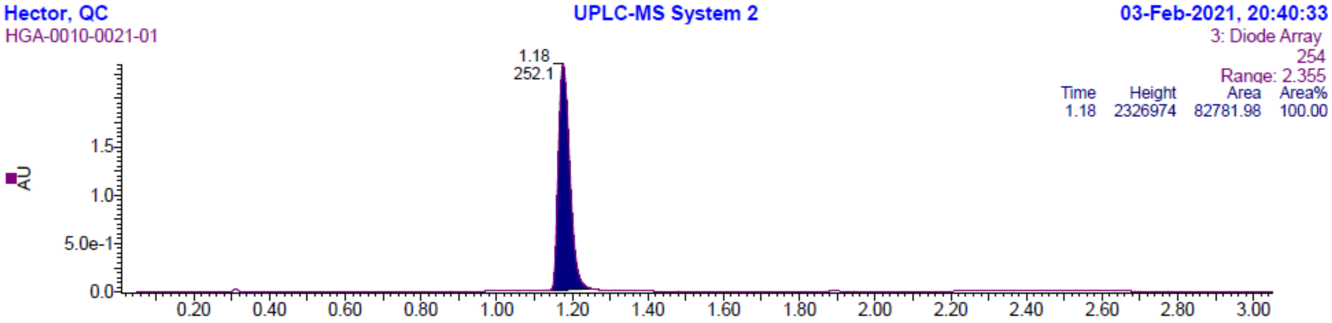


LC: Method 1, $t_R$ = 1.18 min. MS (ESI): m/z = 352.94 [M - H]$^-$

**[S12]**

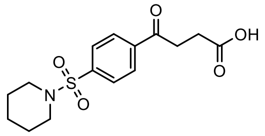


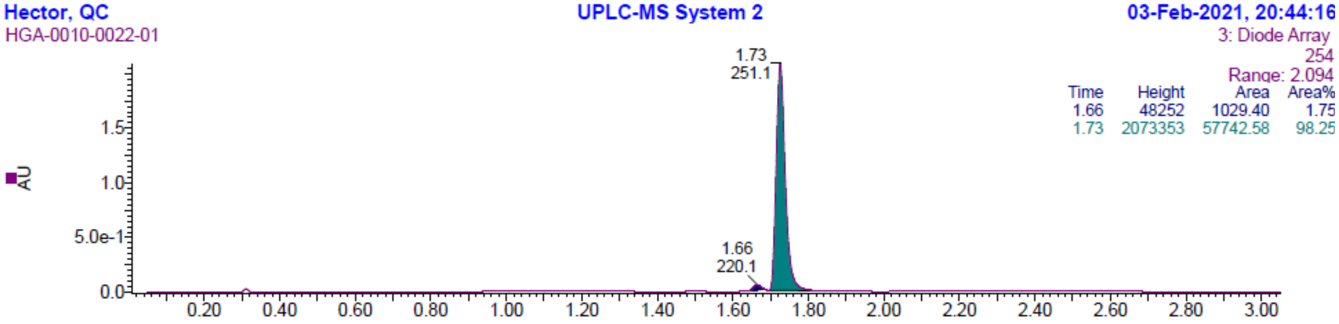


LC: Method 1, $t_R$ = 1.73 min. MS (ESI): m/z = 324.17 [M - H]$^-$

**[S13]**

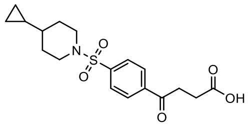

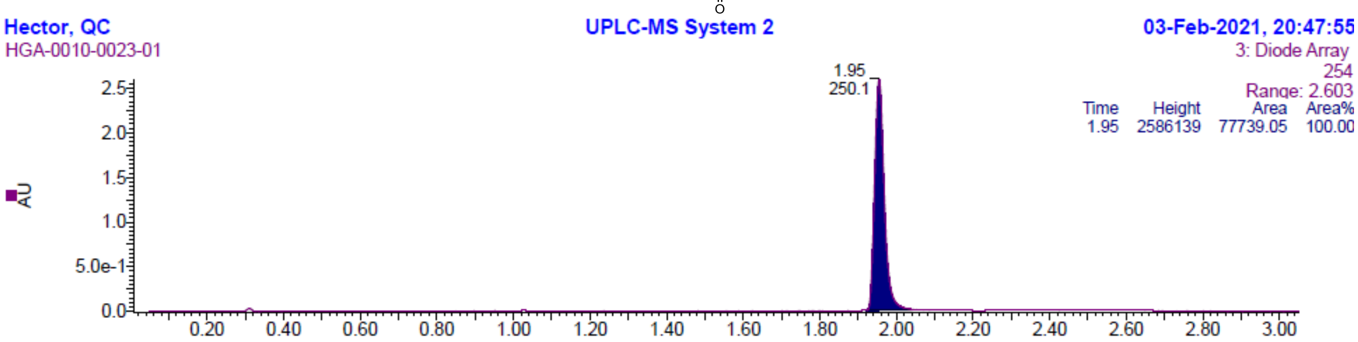


LC: Method 1, $t_R$= 1.95 min. MS (ESI): *m/z*= 366.10 $[M + H]^+$

**[S14]**

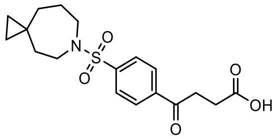

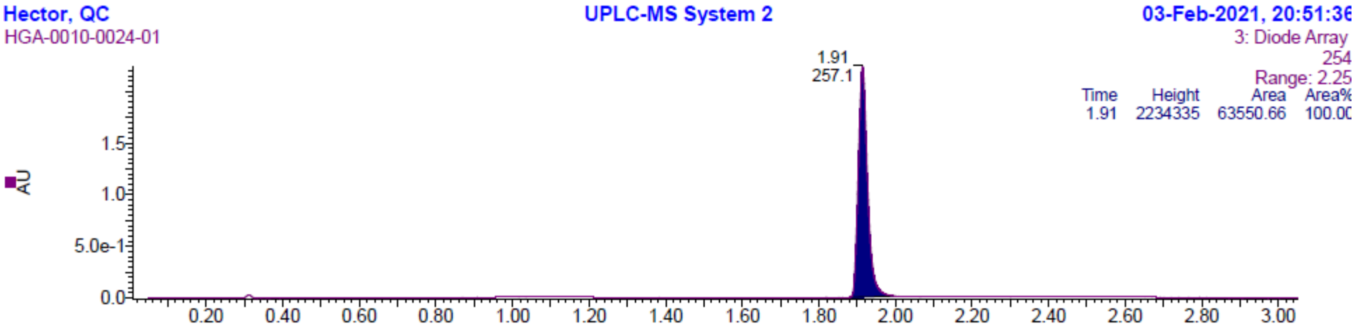


LC: Method 1, $t_R$= 1.91 min. MS (ESI): *m/z*= 365.92 $[M + H]^+$

**[S15]**

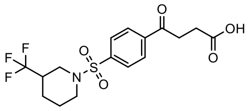

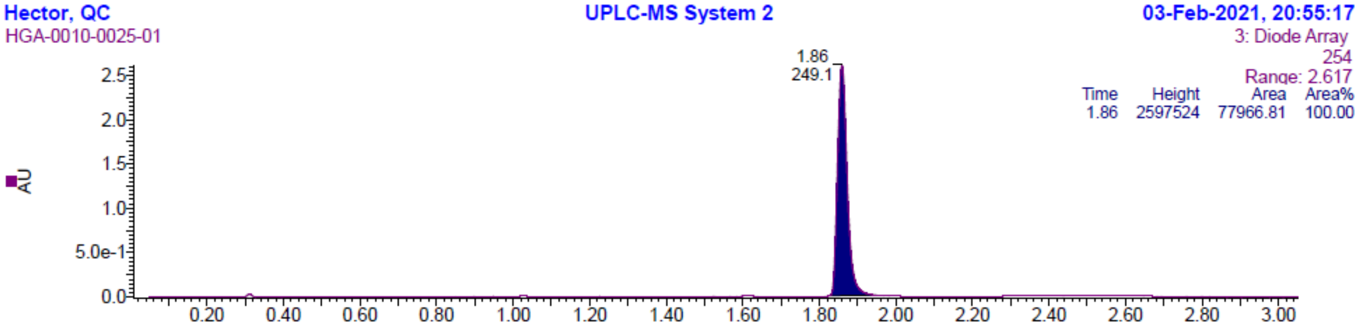


LC: Method 1, $t_R$ = 1.86 min. MS (ESI): m/z = 392.18 $[M - H]^-$

**[S16]**

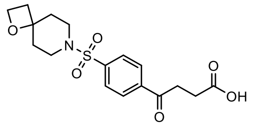

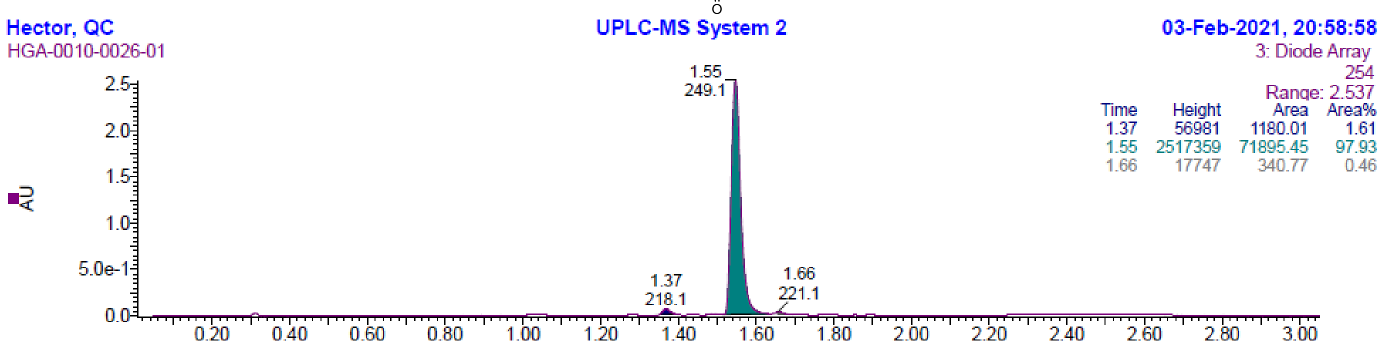


LC: Method 1, $t_R$ = 1.55 min. MS (ESI): m/z = 366.17 $[M - H]^-$

**[S17]**

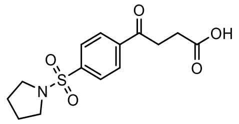

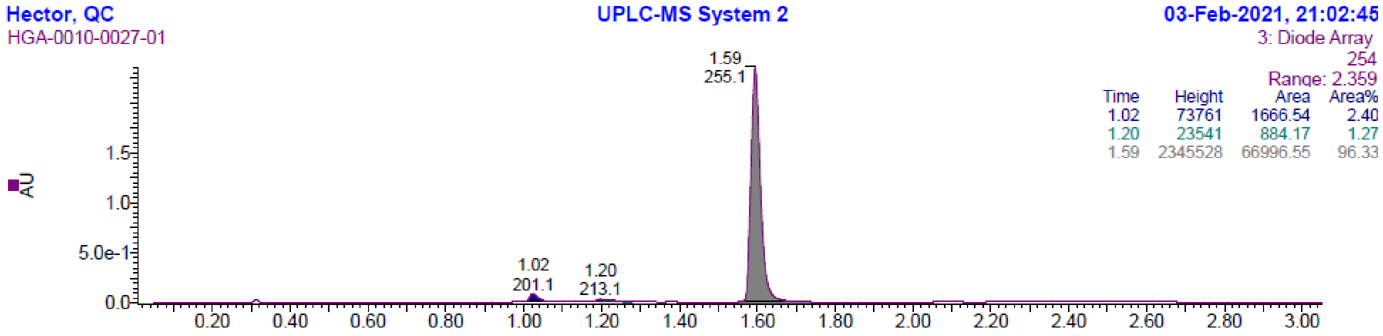


LC: Method 1, $t_R$ = 1.59 min. MS (ESI): m/z = 310.00 $[M - H]^-$

**[S18]**

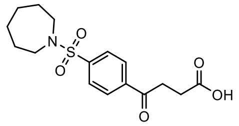

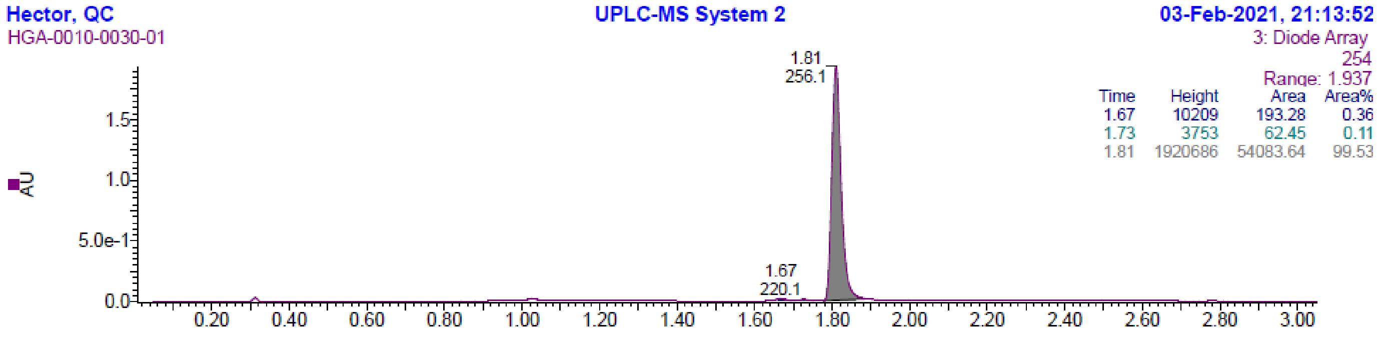


LC: Method 1, $t_R$ = 1.81 min. MS (ESI): m/z = 338.08 $[M - H]^-$

## [S19]

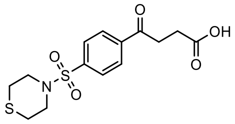

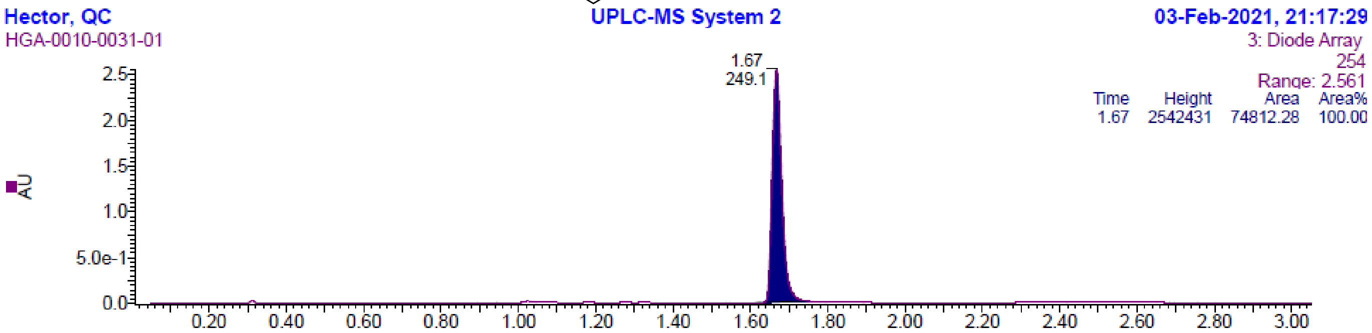


LC: Method 1, $t_R$ = 1.67 min. MS (ESI): m/z = 342.22 $[M - H]^-$

## [S20]

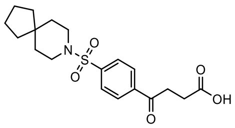

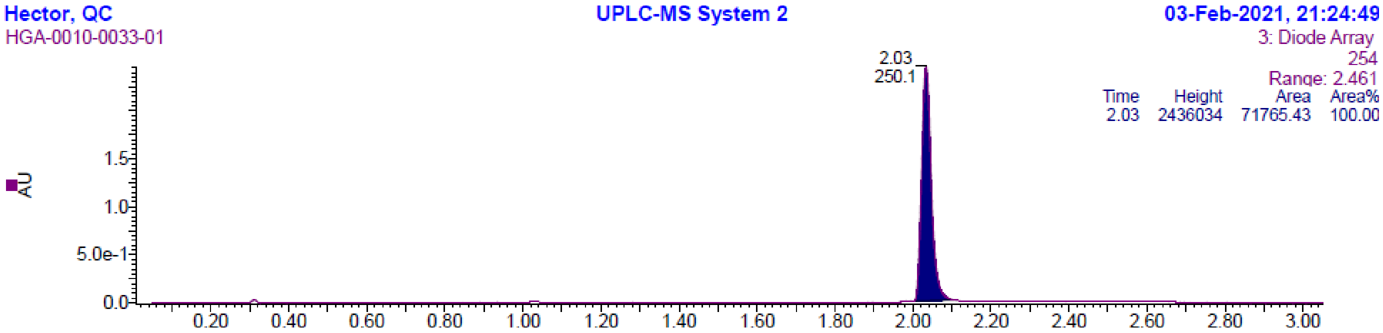


LC: Method 1, $t_R$ = 2.03 min. MS (ESI): m/z = 378.20 $[M - H]^-$

## [S21]

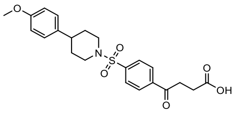

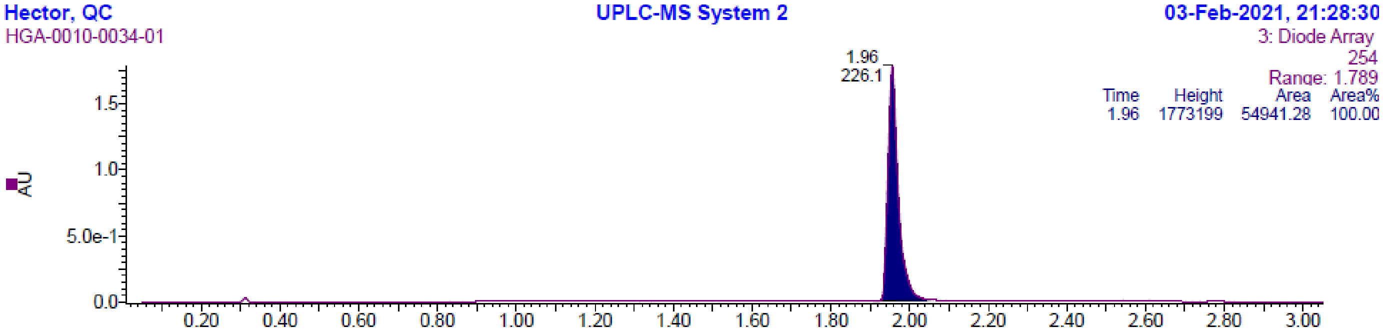


LC: Method 1, $t_R$= 1.96 min. MS (ESI): *m/z*= 432.30 $[M + H]^+$

**[S22]**

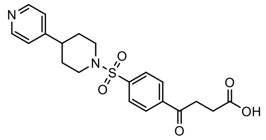

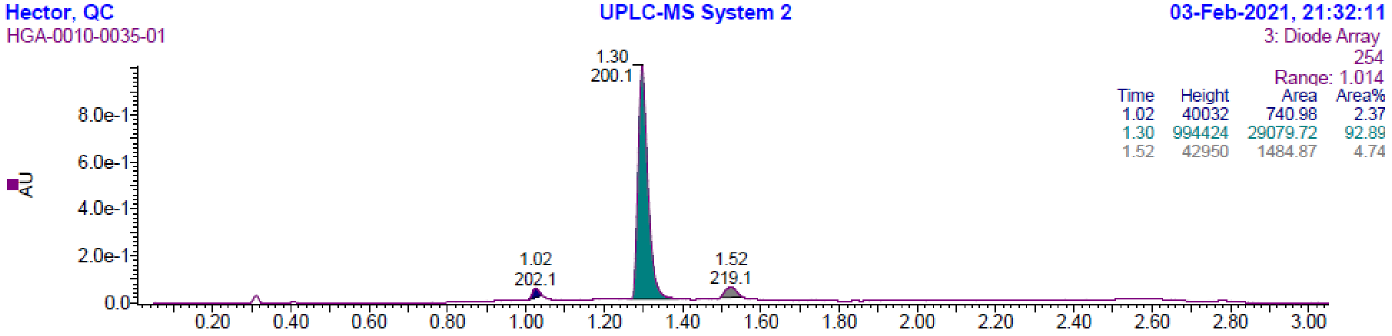


LC: Method 1, $t_R$ = 1.30 min. MS (ESI): m/z = 401.02 [M - H]$^-$

**[S24]**

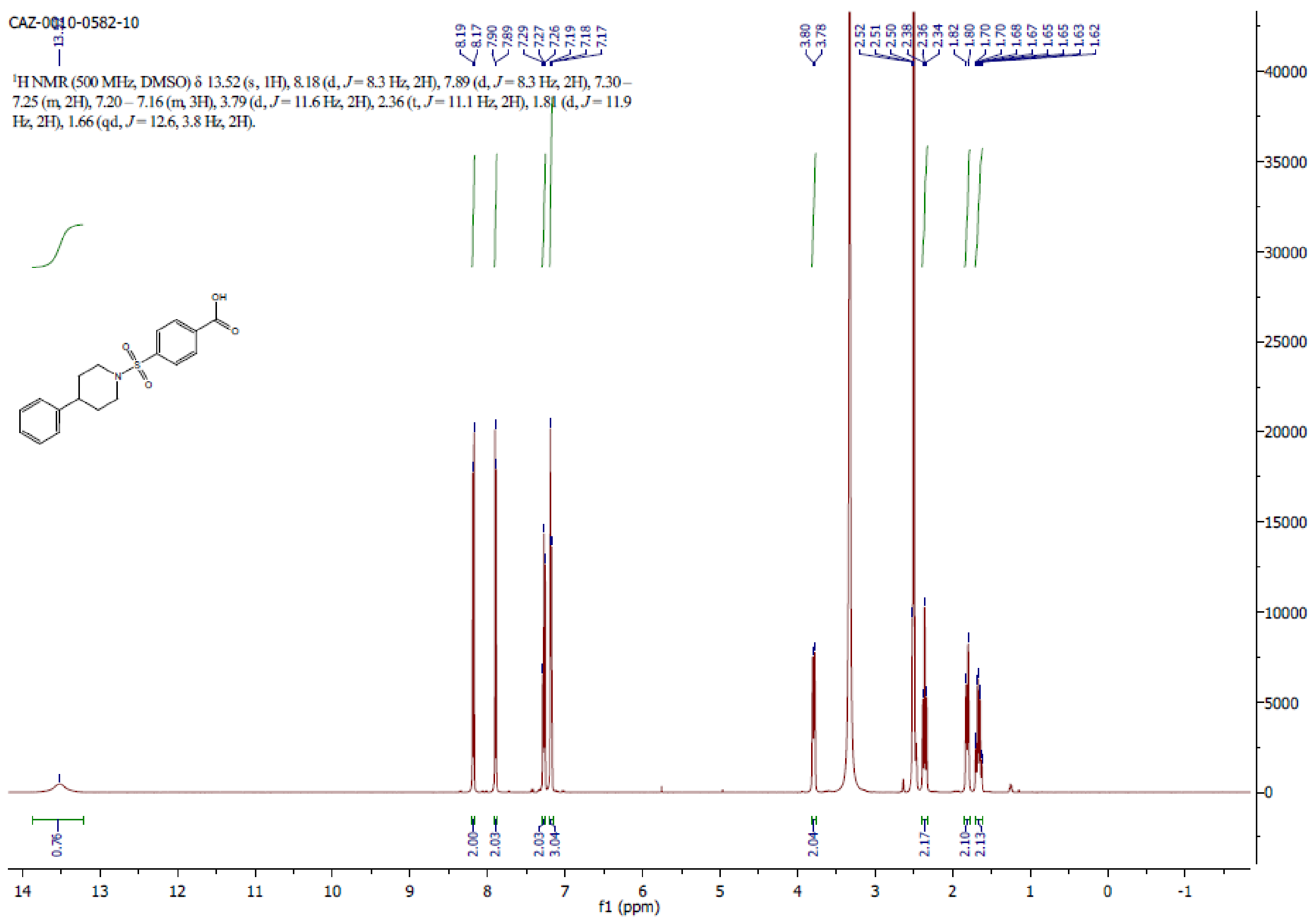

## [S25]

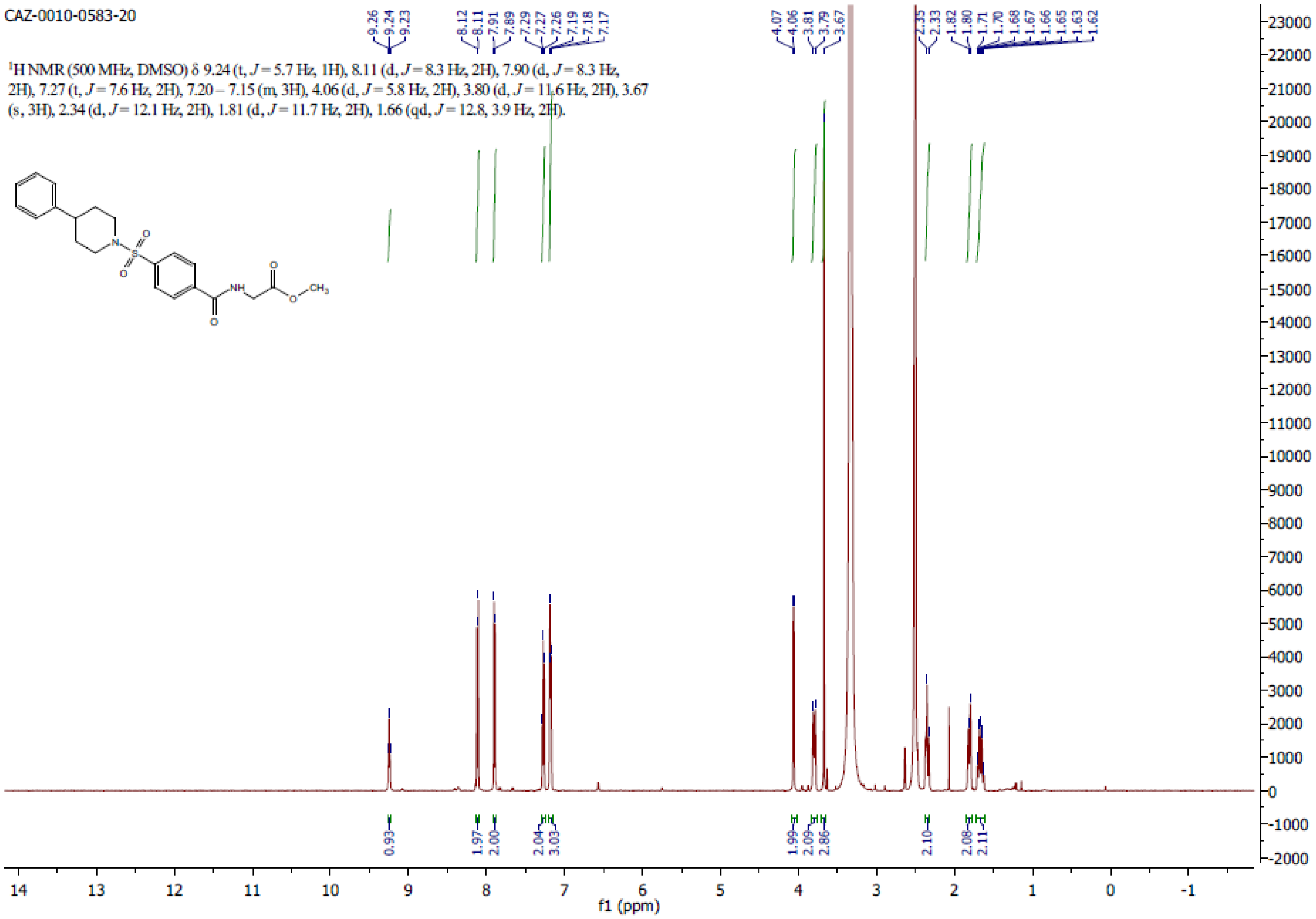
CAZ-0010-0583-20
1H NMR (500 MHz, DMSO) δ 9.24 (t, J = 5.7 Hz, 1H), 8.11 (d, J = 8.3 Hz, 2H), 7.90 (d, J = 8.3 Hz, 2H), 7.27 (t, J = 7.6 Hz, 2H), 7.20 – 7.15 (m, 3H), 4.06 (d, J = 5.8 Hz, 2H), 3.80 (d, J = 11.6 Hz, 2H), 3.67 (s, 3H), 2.34 (d, J = 12.1 Hz, 2H), 1.81 (d, J = 11.7 Hz, 2H), 1.66 (qd, J = 12.8, 3.9 Hz, 2H).
f1 (ppm)

## [S26]

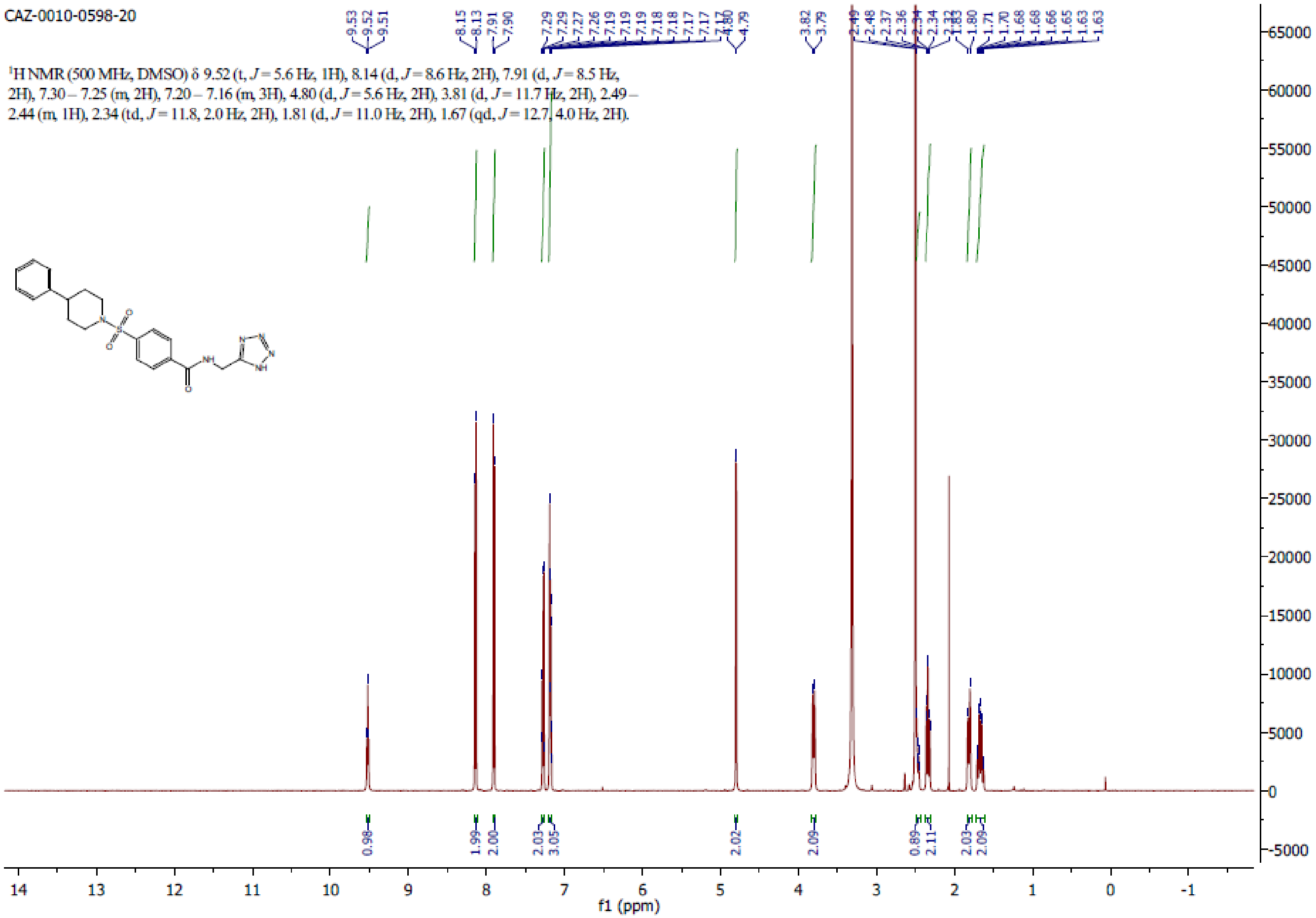

CAZ-0010-0598-20
[1]H NMR (500 MHz, DMSO) δ 9.52 (t, J = 5.6 Hz, 1H), 8.14 (d, J = 8.6 Hz, 2H), 7.91 (d, J = 8.5 Hz, 2H), 7.30 – 7.25 (m, 2H), 7.20 – 7.16 (m, 3H), 4.80 (d, J = 5.6 Hz, 2H), 3.81 (d, J = 11.7 Hz, 2H), 2.49 – 2.44 (m, 1H), 2.34 (td, J = 11.8, 2.0 Hz, 2H), 1.81 (d, J = 11.0 Hz, 2H), 1.67 (qd, J = 12.7, 4.0 Hz, 2H).
f1 (ppm)

**[S27]**

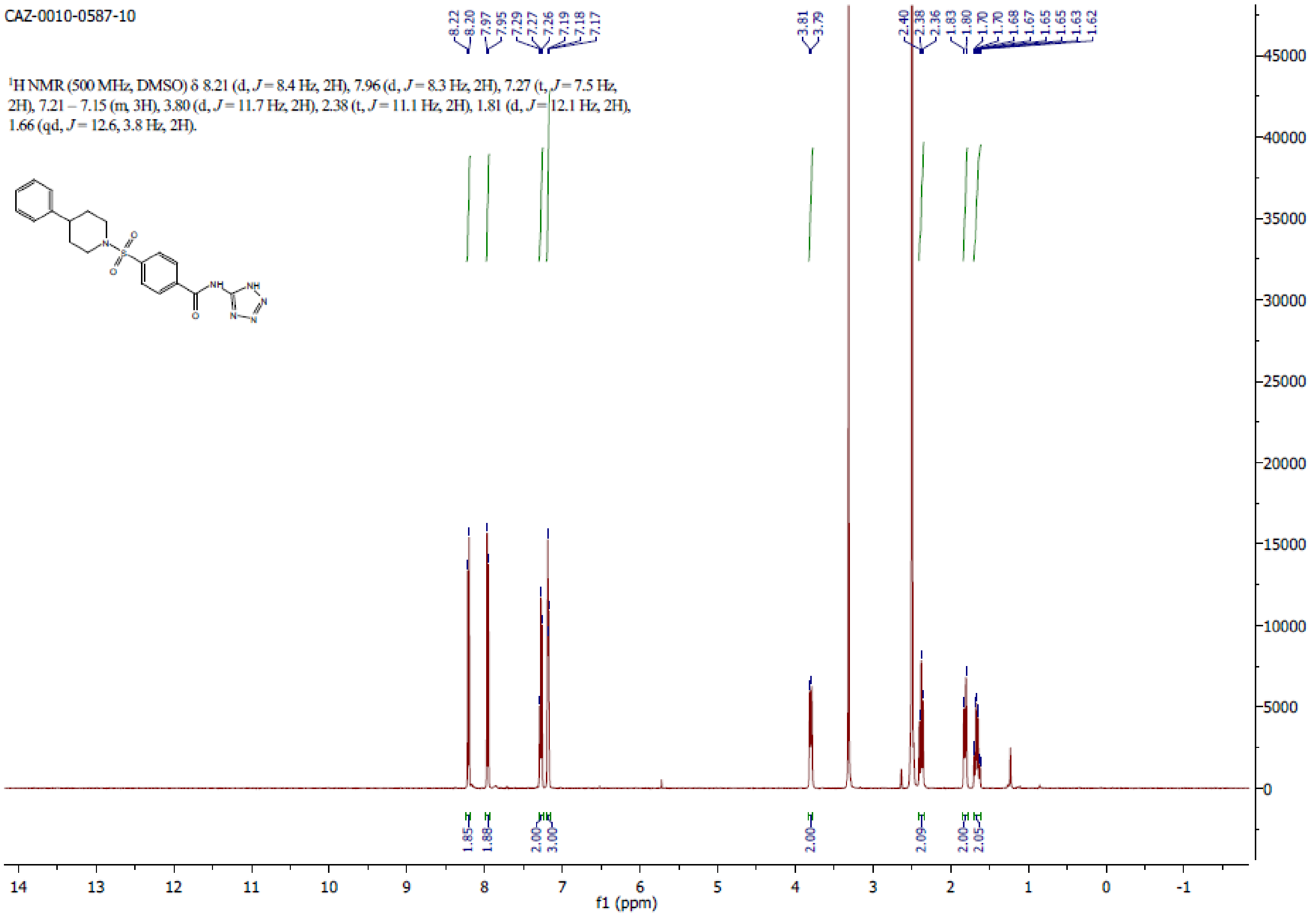

CAZ-0010-0587-10
¹H NMR (500 MHz, DMSO) δ 8.21 (d, J = 8.4 Hz, 2H), 7.96 (d, J = 8.3 Hz, 2H), 7.27 (t, J = 7.5 Hz, 2H), 7.21 – 7.15 (m, 3H), 3.80 (d, J = 11.7 Hz, 2H), 2.38 (t, J = 11.1 Hz, 2H), 1.81 (d, J = 12.1 Hz, 2H), 1.66 (qd, J = 12.6, 3.8 Hz, 2H).
8.22 8.20 7.97 7.95 7.29 7.27 7.26 7.19 7.18 7.17
3.81 3.79
2.40 2.38 2.36 1.83 1.80 1.70 1.70 1.68 1.67 1.65 1.65 1.63 1.62
1.85 1.88 2.00 3.00 2.00 2.09 2.00 2.05
f1 (ppm)

**[S28]**

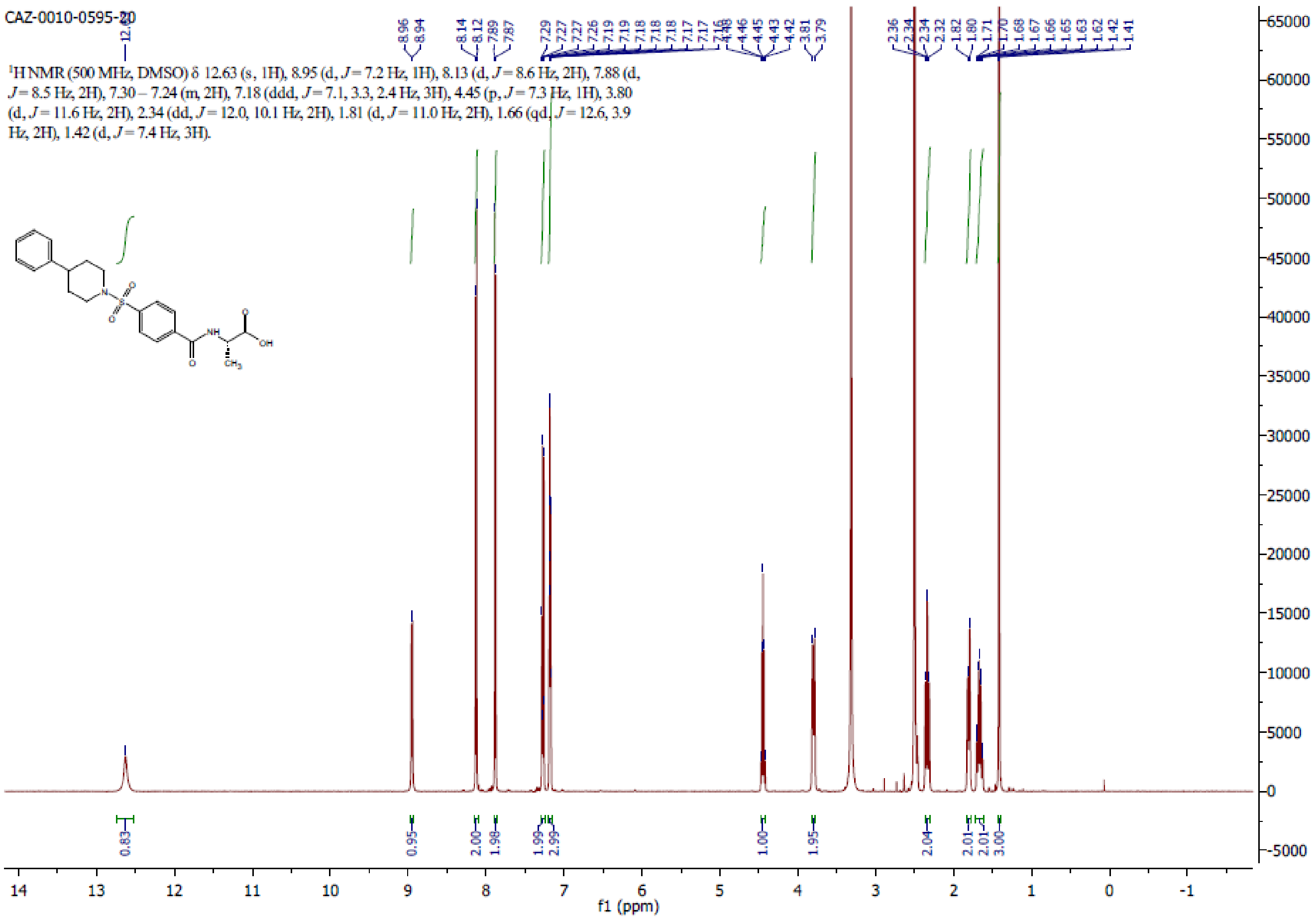
CAZ-0010-0595-20
¹H NMR (500 MHz, DMSO) δ 12.63 (s, 1H), 8.95 (d, J = 7.2 Hz, 1H), 8.13 (d, J = 8.6 Hz, 2H), 7.88 (d, J = 8.5 Hz, 2H), 7.30 – 7.24 (m, 2H), 7.18 (ddd, J = 7.1, 3.3, 2.4 Hz, 3H), 4.45 (p, J = 7.3 Hz, 1H), 3.80 (d, J = 11.6 Hz, 2H), 2.34 (dd, J = 12.0, 10.1 Hz, 2H), 1.81 (d, J = 11.0 Hz, 2H), 1.66 (qd, J = 12.6, 3.9 Hz, 2H), 1.42 (d, J = 7.4 Hz, 3H).
f1 (ppm)

**[S29]**

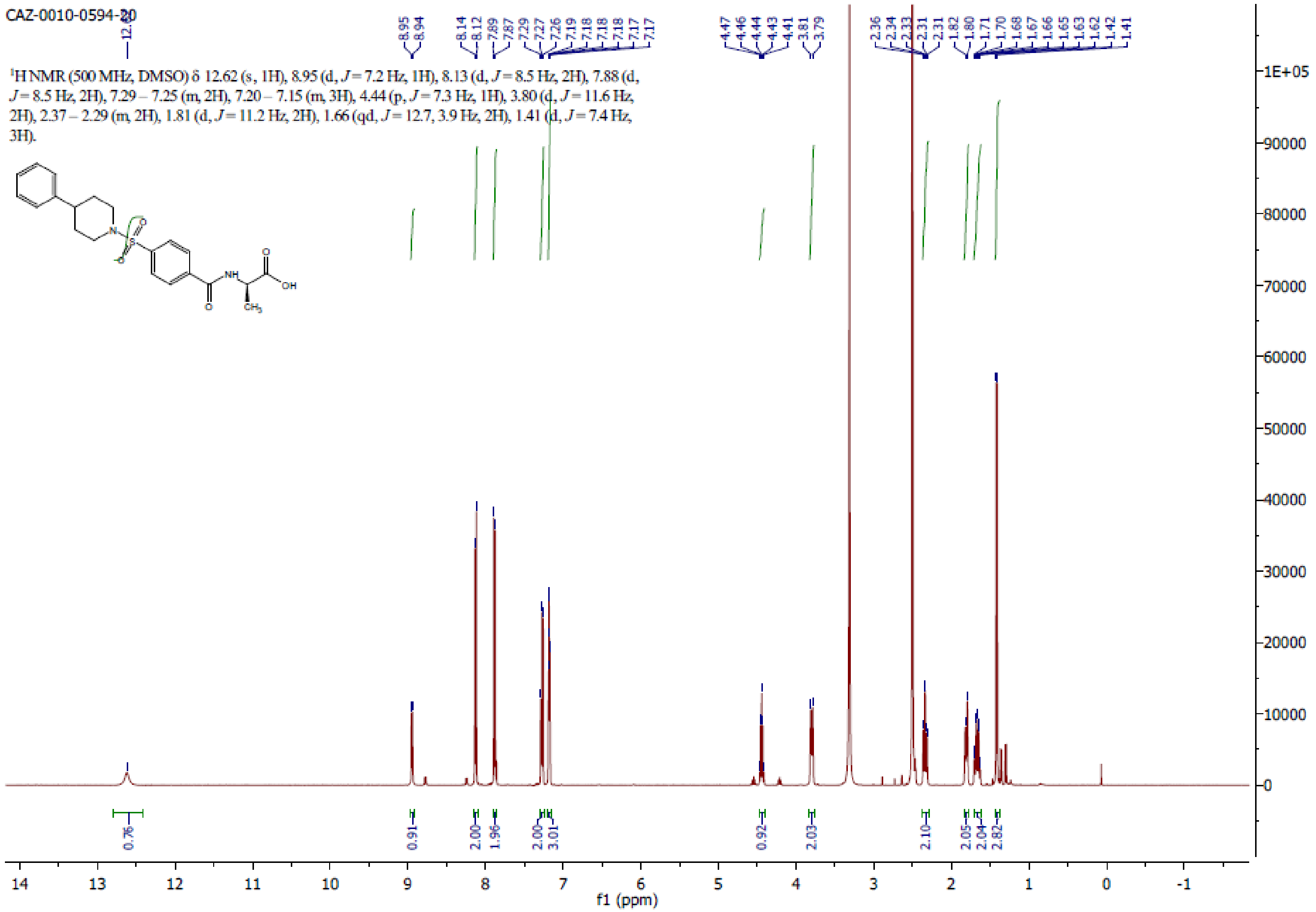

CAZ-0010-0594-20
¹H NMR (500 MHz, DMSO) δ 12.62 (s, 1H), 8.95 (d, J = 7.2 Hz, 1H), 8.13 (d, J = 8.5 Hz, 2H), 7.88 (d, J = 8.5 Hz, 2H), 7.29 – 7.25 (m, 2H), 7.20 – 7.15 (m, 3H), 4.44 (p, J = 7.3 Hz, 1H), 3.80 (d, J = 11.6 Hz, 2H), 2.37 – 2.29 (m, 2H), 1.81 (d, J = 11.2 Hz, 2H), 1.66 (qd, J = 12.7, 3.9 Hz, 2H), 1.41 (d, J = 7.4 Hz, 3H).
f1 (ppm)

# [S30]

CAZ-0010-0596-20

[1]H NMR (500 MHz, DMSO) δ 12.27 (s, 1H), 8.74 (s, 1H), 8.09 (d, $J$ = 8.6 Hz, 2H), 7.86 (d, $J$ = 8.5 Hz, 2H), 7.29 – 7.24 (m, 2H), 7.20 – 7.16 (m, 3H), 3.79 (d, $J$ = 11.6 Hz, 2H), 2.47 (tt, $J$ = 12.2, 3.5 Hz, 1H), 2.32 (td, $J$ = 11.9, 2.1 Hz, 2H), 1.81 (d, $J$ = 11.0 Hz, 2H), 1.67 (qd, $J$ = 12.7, 4.0 Hz, 2H), 1.48 (s, 6H).

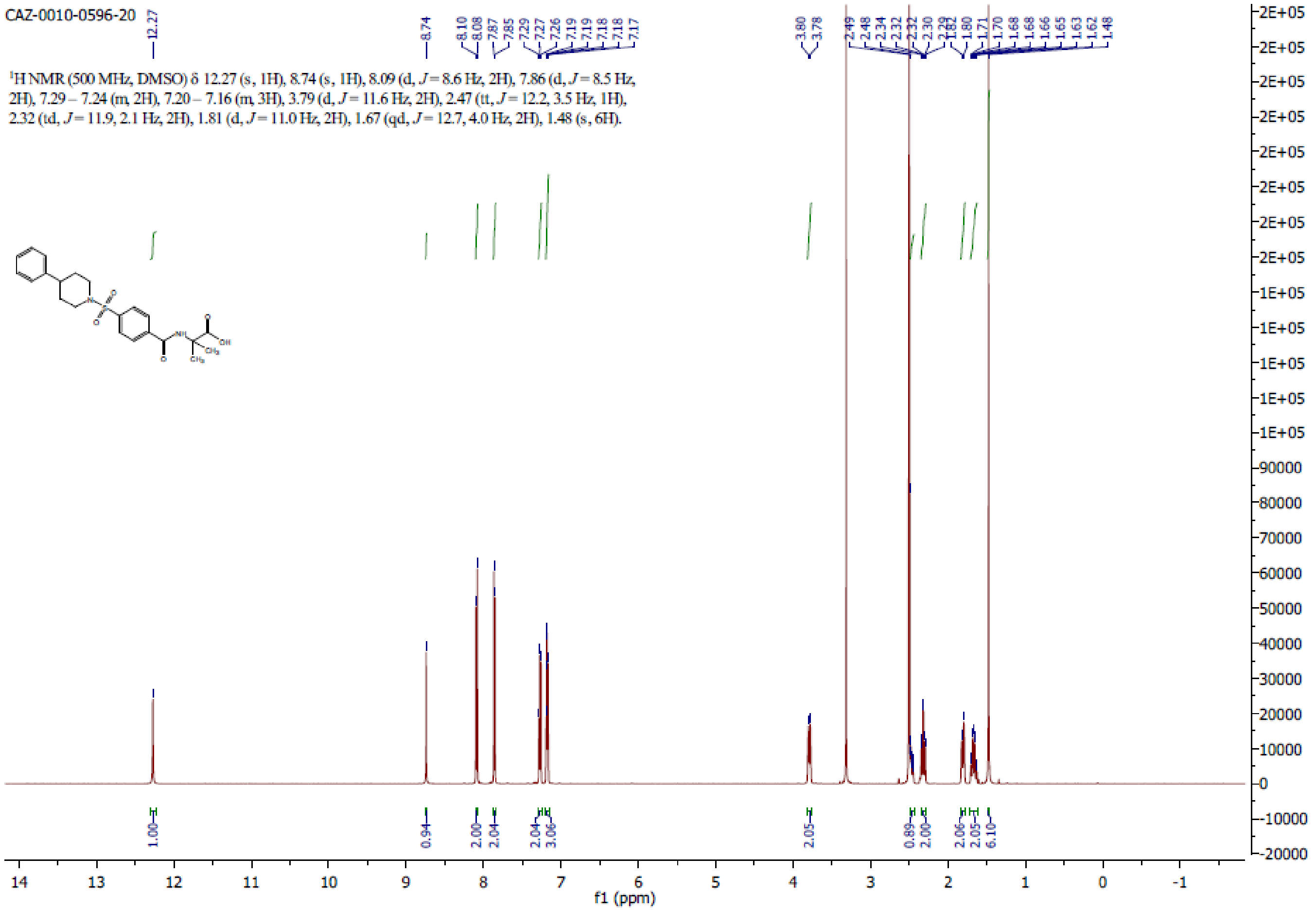

**[S31]**

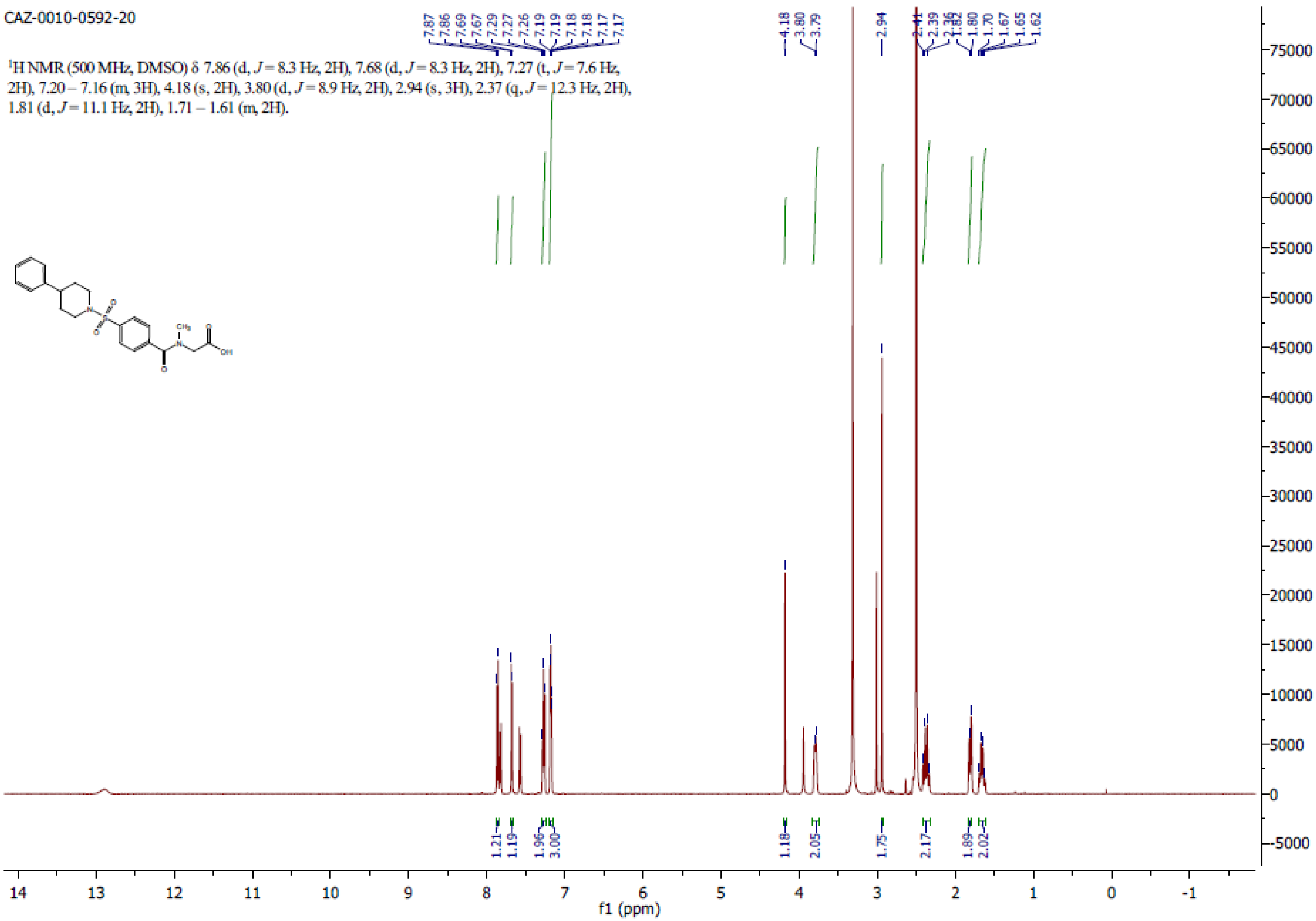
CAZ-0010-0592-20
¹H NMR (500 MHz, DMSO) δ 7.86 (d, J = 8.3 Hz, 2H), 7.68 (d, J = 8.3 Hz, 2H), 7.27 (t, J = 7.6 Hz, 2H), 7.20 – 7.16 (m, 3H), 4.18 (s, 2H), 3.80 (d, J = 8.9 Hz, 2H), 2.94 (s, 3H), 2.37 (q, J = 12.3 Hz, 2H), 1.81 (d, J = 11.1 Hz, 2H), 1.71 – 1.61 (m, 2H).
f1 (ppm)

**[S32]**

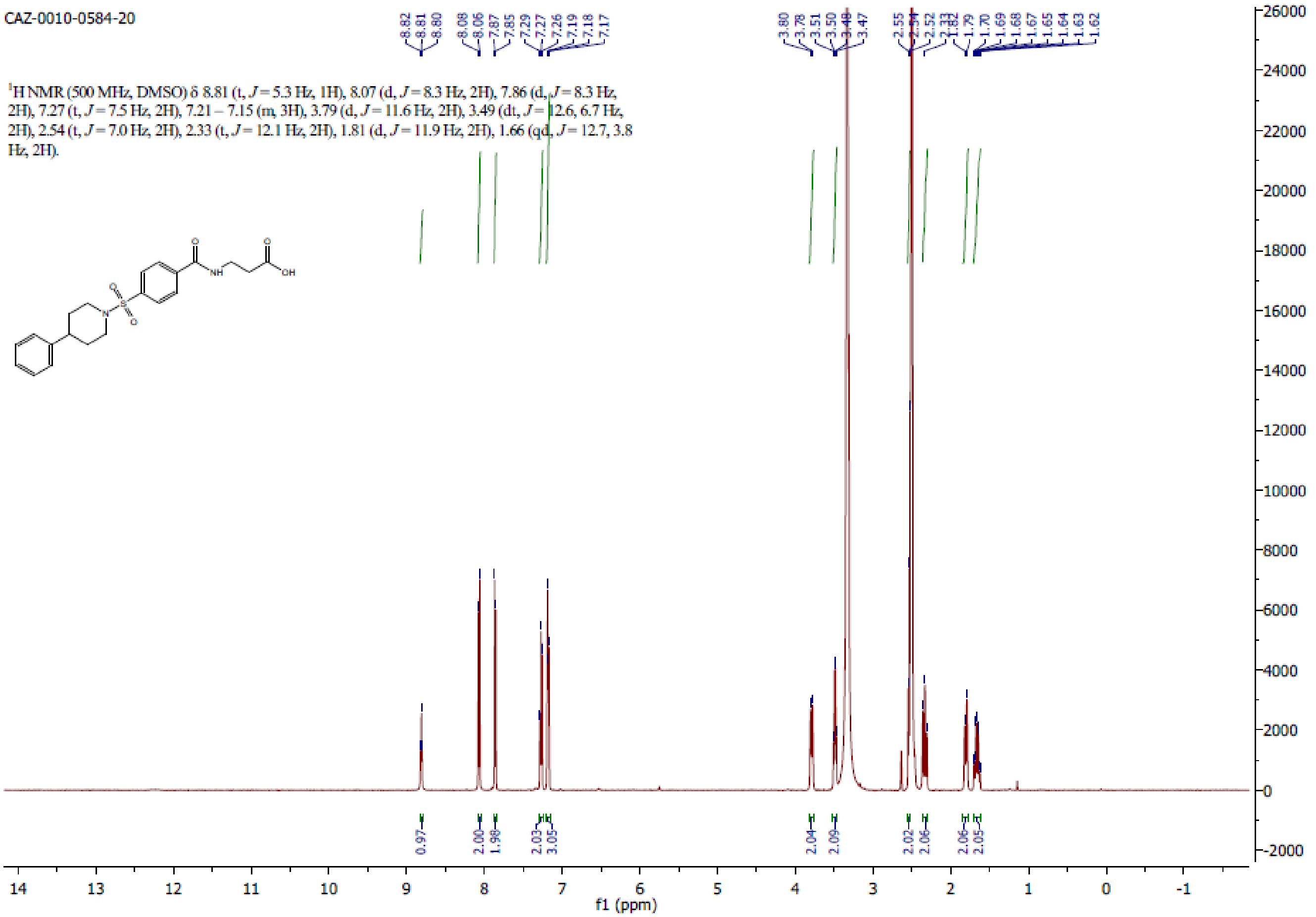
CAZ-0010-0584-20
1H NMR (500 MHz, DMSO) δ 8.81 (t, J = 5.3 Hz, 1H), 8.07 (d, J = 8.3 Hz, 2H), 7.86 (d, J = 8.3 Hz, 2H), 7.27 (t, J = 7.5 Hz, 2H), 7.21 – 7.15 (m, 3H), 3.79 (d, J = 11.6 Hz, 2H), 3.49 (dt, J = 12.6, 6.7 Hz, 2H), 2.54 (t, J = 7.0 Hz, 2H), 2.33 (t, J = 12.1 Hz, 2H), 1.81 (d, J = 11.9 Hz, 2H), 1.66 (qd, J = 12.7, 3.8 Hz, 2H).
f1 (ppm)
